# CryoFold: determining protein structures and ensembles from cryo-EM data

**DOI:** 10.1101/687087

**Authors:** Mrinal Shekhar, Genki Terashi, Chitrak Gupta, Daipayan Sarkar, Gaspard Debussche, Nicholas J. Sisco, Jonathan Nguyen, Arup Mondal, James Zook, John Vant, Petra Fromme, Wade D. Van Horn, Emad Tajkhorshid, Daisuke Kihara, Ken Dill, Alberto Perez, Abhishek Singharoy

## Abstract

Cryo-EM is a powerful method for determining protein structures. But it requires computational assistance. Physics-based computations have the power to give low-free-energy structures and ensembles of populations, but have been computationally limited to only small soluble proteins. Here, we introduce CryoFold. By integrating data of varying sparsity from electron density maps of 3–5 Å resolution with coarse-grained physical knowledge of secondary and tertiary interactions, CryoFold determines ensembles of protein structures directly from sequence. We give six examples showing its broad capabilities, over proteins ranging from 72 to 2000 residues, including membrane and multi-domain proteins, and including results from two EMDB competitions. The ensembles CryoFold predicts starting from the density data of a single known protein conformation encompass multiple low-energy conformations, all of which are experimentally validated and biologically relevant.

---

Cryo-electron microscopy (cryo-EM) is a powerful tool for determining the structures of biomolecules. It serves a niche – such as large complexes or membrane proteins or molecules that are not easily crystallizable – that traditional methods, such as X-ray diffraction, electron or neutron scattering, or NMR often cannot handle. Routine cryo-EM structure determination has a number of components: the experiment produces raw data in the form of single-particle images, correction and processing of this data recovers an electron density map, and finally molecular modeling is required to determine structures from the map. Currently, there are two broad classes of methods for molecular modeling. First, established algorithms for refining X-ray crystallography or NMR structures, such as *Phenix*.*realspaceref ine* or REFMAC (1), are often used, even for ensemble determination (2), but offer complete models with the highest-resolution density data. Cryo-EM studies commonly produce lower-resolution data. Second, *integrative approaches* that leverage data from multiple types of experiments (3) to find structures compatible with the data. The challenge here is that cryo-EM data is often *heterogeneous*, meaning that some parts of a protein structure are well-determined by the data while others are more poorly defined.

For computational modeling, the changing resolution poses the need for extensive conformational sampling and the need to identify which conformations amongst all that fit the lower resolution regions are most biophyically relevant. The size of the search space is large and grows non-linearly with system size (4). Physics-based modeling, such as molecular dynamics (MD) simulations, can give proper thermodynamic weights for choosing among the different conformational populations. But, we need efficient ways of sampling using physics based approaches. Most MD is used for exploring dynamics around an experimental structure and for automated model refinement (5, 6). Yet, large conformational changes, such as those relevant in many biological processes, remain inaccessible to MD(7–9) - it is computationally expensive. In structure determination, the end structure is unknown, so collective variables to accelerate the process are not an option (10). Therefore, MD is augmented with external information such as evolutionary covariance (11, 12) and homology-based starting models (13, 14), or with advanced sampling methods based on Bayesian inference (15–17) and specialized hardware (18), which improve the speed of structure prediction by 10 to 100-fold over brute-force simulations. Notwithstanding this improvement, the prediction of protein fragments beyond 115 residues remain a bottleneck for physics-based methods (19). Fragment search and fitting schemes are successful in resolving the EM map(20), but they require at least 70% of the C*α* atoms placed correctly (21–23), and for membrane systems, such refinements also leverage MD simulations (24). However, the bioinformatic augmentations to MD introduce new discrepancies that are often refractory to automated fixes (23, 25), warranting our developments.

Here, we describe CryoFold, an integrative atomistic-physical algorithm that derives ensemble of folded protein structures from cryo-EM data. Illustrated in **(Fig. 1)**, CryoFold is a combination of three methods: **(1)** MAINMAST(26), MAINchain Model trAcing from Spanning Tree – a method that generates the trace of the connected peptide chain when provided with EM data, **(2)** ReMDFF(27), Resolution exchange Molecular Dynamics Flexible Fitting – a MD method for refining protein conformations from electron-density maps, and **(3)** MELD(15, 28), Modeling Employing Limited Data – a Bayesian folding and refolding engine that can work from insufficient data to accelerate the MD sampling of rare events such as those needed for protein folding. The guidance from experimental data allows MD simulations to fold models with well beyond 115 residues, including transmembrane systems and asymmetric multi-protein complexes. More importantly, the free energy description of folded and unfolded populations accessible to MELD enables the exhaustive sampling of structures that are representative of different metastable states. Thus, starting with the structural data from a particular protein conformation, CryoFold predicts on one hand, the energetically favorable ensemble of structures that are consistent with the data, while on the other hand, discovers multiple new low-energy protein states in the vicinity of the fitted model. Going beyond the determination of one stationary structure, CryoFold offers the opportunity to combine all the predicted structures into a model conformational transition pathway, where the new states are also validated and re-refined against orthogonal NMR, X-ray crystallography or cryo-EM data.

**Fig. 1.**
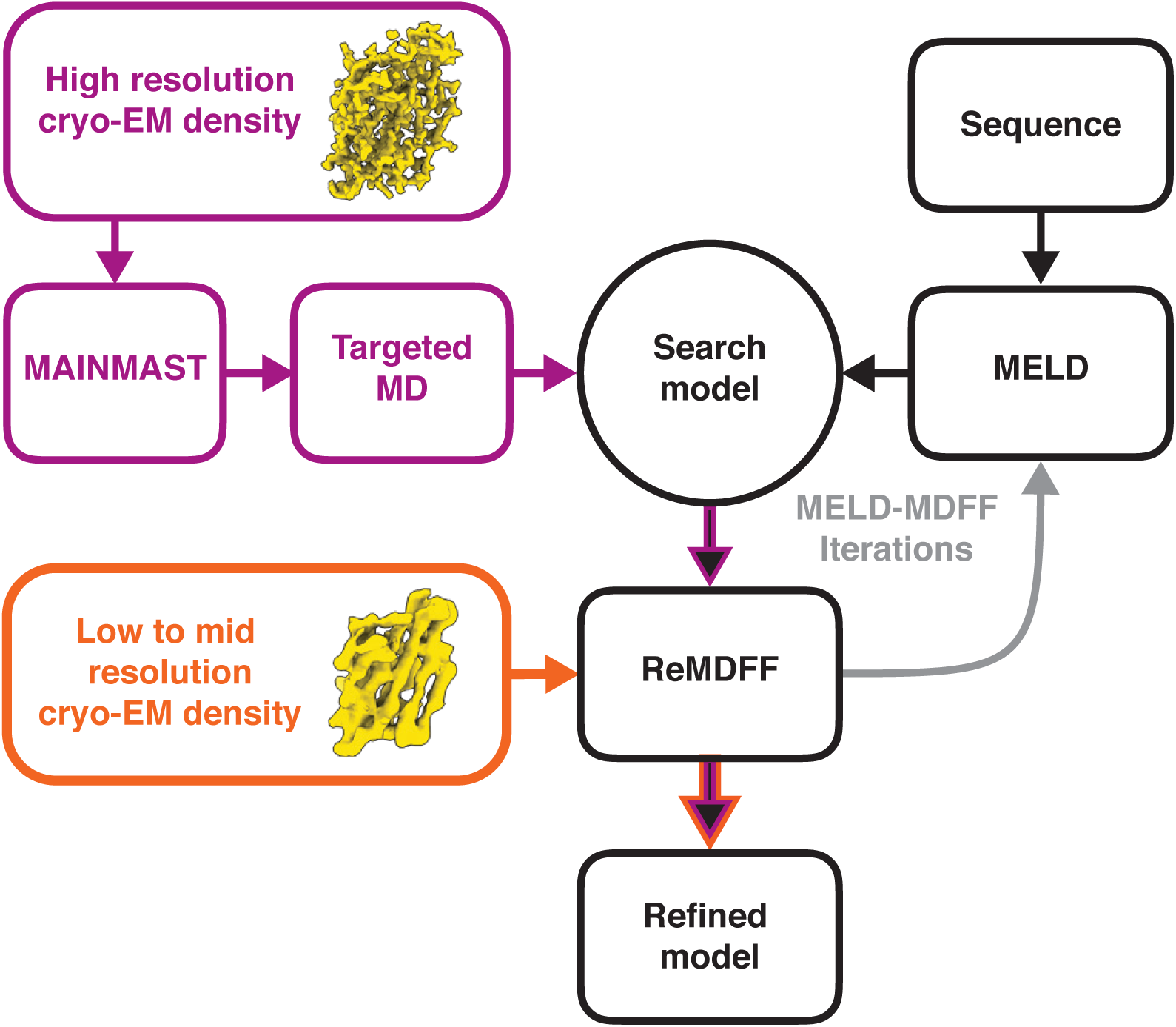
An overview of the CryoFold protocol. For a high-resolution density map (data-rich case), backbone tracing is performed using MAINMAST to determine C*α* positions, and a random coil is fitted to these positions using targeted MD. This fitted protein model is subjected to the next MELD-ReMDFF cycles as a search model. For a low or medium resolution density map (data-poor case), a search model is constructed from primary sequence using MELD. This search model is fitted into the electron density using ReMDFF. The ReMDFF output is fed back to MELD for the next iteration, and the cycle continues until convergence.

Starting with density maps of resolution 5.0 Å and higher, first, MAINMAST is employed to derive a chain trace of C_*α*_ atoms. Then we use this trace as a template to iterate between MELD and ReMDFF. While MELD explores a large conformational space, visiting multiple plausible secondary structures consistent with the MAINMAST template, ReMDFF simulations refine the protein backbone and sidechain conformations to fit to the density map for each one of the assumed secondary structures (6). Taken alone, ReMDFF fits models into electron density features, but fails to explore the variations in secondary structures(27). MELD addresses this issue by partial folding, unfolding and reformation of secondary structures(15, 28), using the coarse physical information (CPI) available on webservers(15, 29); for example, based on their sequences, proteins prefer specific fractions of hydrophobic interactions, *β*-strand pairing and secondary structures to minimize frustration **(Fig. 2A)**. Consequently, a hybrid iterative MELD-ReMDFF approach allows the determination of complete all-atom models from sequence information merged with available structural data of varying coarseness. For intermediate to low-resolution data (lesser than 5 Å) wherein C-alpha tracing is unreliable (30), the MAINMAST step can be avoided. Nonetheless, if successful, the search template derived from backbone tracing almost always accelerates convergence of CryoFold.

**Fig. 2.**
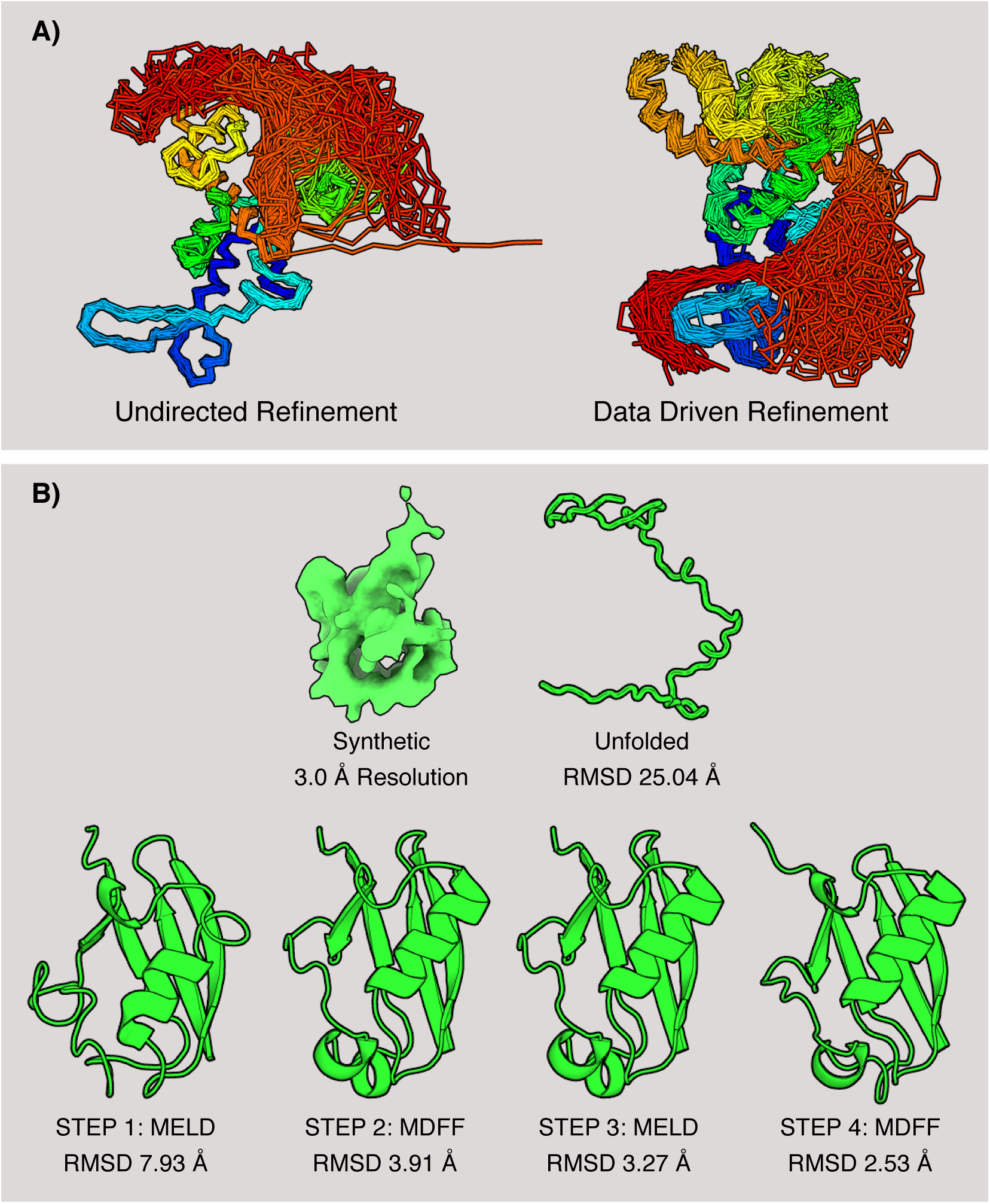
Ensemble models for TRPV1 and the refinement protocol for ubiquitin. **(A)** Ensemble refinement with CryoFold showcased for the soluble domain of TRPV1. Several conformations from the TRPV1 ensemble are superimposed; color coding from blue (N-terminal) to red (C-terminal). In a MELD-only simulation, a soluble loop (indicated in red) artifactually interacted with the transmembrane domains. Following the data-guidance from ReMDFF, this loop interacted with the soluble domains and a more focused ensemble is derived that agrees with the electron density. **(B)** Stages of the refinement protocol for a test case, ubiquitin. The initial model is an unfolded coil. MELD was used to generate 50 search models from just the amino acid sequence, and no usage of the electron density data. Then, these models were rigid-fitted into the electron density using Chimera(59), and ranked based on their global cross-correlation. ReMDFF refined the best rigid-fitted model even further. The ReMDFF model with the highest Cross Correlation (CC) to the density map served as a template for the subsequent iteration with MELD. In two consecutive MELD-ReMDFF iterations the RMSD of the folded model relative to the crystal structure (1UBQ) attenuated from 25.04 Å to 2.53 Å

We report data-guided structural ensembles for six different examples here, for proteins from 72 to 618 residues, extending to multi-protein complexes of up to 2000 residues, and across both soluble and membrane systems. CryoFold overcomes the sampling limitations of traditional MD predictions, producing high-quality structural models: it offers a high radius of convergence in the range of 50 Å, refining soluble and transmembrane structures with consistently *>* 90% favored backbone and sidechain statistics, and high EMRinger scores (31). The results are independent of the initial estimated conformation and consistent with physics and stereochemistry, highlighted through results in 2016 and 2019 EMDB competitions. The hybrid protocol is available through a python-based graphical user interface with a video tutorial.

## Results

We describe six systems, chosen to represent the different bottlenecks in the three component methods of the CryoFold pipeline. At any given resolution, the accuracy of CryoFold predictions depends on: **(1)** quality of C-_*α*_ traces by MAIN-MAST, **(2)** variations in secondary structure within the MELD ensemble, and **(3)** convergence of ReMDFF. Three are soluble proteins, with varying degrees of local resolution in the density maps. One is from the 2019 EMDB competition challenge, in which the data was provided at three different resolutions. One was a large asymmetric multi-protein complex that allowed us to test how big a structure we could handle. And, one was a transmembrane system, to see if MELD’s aqueous implicit-solvent model would be sufficient for the membrane environment.

### A. Proof of principle on a small known protein

In this case, we began with a synthetic map of ubiquitin, a small 72-residue protein. Ubiquitin is a good test system because, on the one hand, it is small enough to fold computationally, and yet on the other hand its experimental folding time is in the millisecond range, so it been hard to fold by brute force MD (32), and even, to a lesser extent, by the MELD approach (15). From the known X-ray crystal structure of ubiquitin, we generated a synthetic electron density at 3.0 Å resolution (33), and asked if CryoFold could correctly recover the X-ray structure. We found that only two MELD-ReMDFF iterations **(Fig. 2B)** were needed to give a model having an RMSD difference of 2.53 Å from the crystal structure (PDB id: 1UBQ, see Table S1).

### B. Test on a soluble lipoprotein with a uniformly high-resolution data

Francisella lipoprotein Flpp3 is a 108 amino acids long membrane-interacting protein that serves as a target for drug development against tularemia(34). In this case, we had two datasets: one at high resolution (1.8 Å) from our Serial Femtosecond X-ray (SFX) crystallography experiments of Flpp3 (See Supplementary Information and (35), and a synthetic one at low resolution (5.0 Å). The point of this test was to see if we could use the low-resolution data to achieve the high-resolution structure. For both sets, we used MAINMAST (26) to introduce the C*α* traces as constraints for MELD **(Fig. 3A,B**). Convergent ensembles derived from this MAINMAST-guided MELD step were then refined by ReMDFF to improve the sidechains until the density was resolved with models of reliable geometry.

**Fig. 3.**
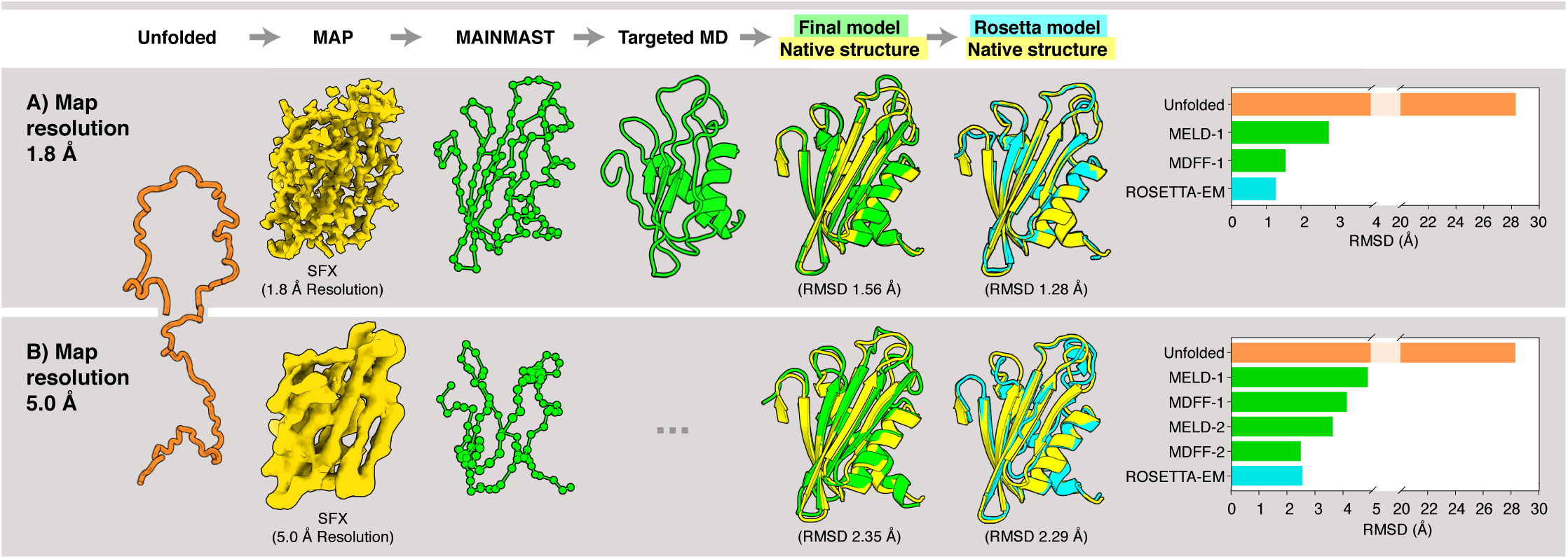
Hybrid structure determination of Flpp3. **(A)** High-resolution density map at 1.8 Å resolution. An unfolded structure was used as the initial model. A SFX density map at 1.8 Å resolution was employed to generate the C*α* position (green spheres) using MAINMAST, and the initial model was fitted into these positions by targeted MD. The resulting structure (green cartoon model) was then subjected to MELD-ReMDFF refinement. This procedure yielded a structure with RMSD of 1.56 Å relative to the native SFX structure (yellow). The Rosetta-EM model (cyan) has an RMSD of 1.28 Å with respect to the SFX structure. **(B)** Lower-resolution density map at 5 Å resolution. An initial C*α* trace in the map was computed using MAINMAST. Subsequent MELD-ReMDFF refinement resulted in a structure (green cartoon model) with an RMSD of 2.29 Å from the SFX structure (yellow). The best Rosetta-EM model has (cyan) an RMSD of 2.35 Å to the SFX structure. Barplots depict the evolution of RMSD of the CryoFold models with each subsequent MELD-ReMDFF refinement.

We found that one iteration of the MELD-ReMDFF cycle sufficed to resolve an all-atom model of Flpp3 from the SFX density, with accurate sidechain conformations, secondary and tertiary structure assignments (structural statistics summarized in Table S2). At 5 Å resolution MAINMAST produced low quality backbone traces (**Fig. 3B**). Remarkably, even these low quality C*α* traces, were enough for MELD-ReMDFF to successfully produced models comparable to our high-resolution refinements. After two MELD-ReMDFF iterations, the best structure obtained was within 2.29 Å RMSD from the SFX model. The MELD-only predictions modelled the *β*-sheets accurately, they failed to accurately converge on all helices (SI Fig. S1). For example, a 4-turn helix was underestimated to contain only 2-3 turns. However, guidance by the density map in CryoFold recovered these turns in both the high and low resolution cases (Tables S2 and S3). Thus, the Flpp3 test shows that the CryoFold trio of methods gives accurate structures for longer chains than is otherwise possible with either one of these methods.

Here, we are also able to test an important aspect of physics-based structure determination, namely whether we can generate proper conformational ensembles, not just single average structures. The quality of the CryoFold ensembles is accessed against a set of 20 NMR models of Flpp3 (34) by looking at the conformation of key residues (Y83,K35 and D4) responsible for binding tularemia drugs (Fig. S2A). Upon projecting the ensemble of 50 lowest-energy CryoFold structures onto a space defined by the distance between Y83-K35 & Y83-D4, where closed Flpp3 is represented by (Y83-K35 *<*5.00 Å & Y83-D4 > 10.00 Å), and open Flpp3 implies (Y83-K35 *>*10.00 Å & Y83-D4 *<* 5.00 Å), all the major conformational states seen in the NMR experiments have been recovered (Fig. S2B). Thus, extending beyond the prediction of a single stationary structure, the cluster of low-energy conformations predicted by CryoFold captures both the open and closed conformations, starting only with data from the closed state. The classification of structural ensembles based on projections onto the distance space requires *a priori* knowledge of the structural features of all the major states in the ensemble. In an alternate scheme that does not require such knowledge, the models were classified based on their Rosetta-energy and RMSD relative to the crystal structure (36). Rosetta is chosen as a benchmark due to its use of energy functions analogous to the CHARMM or AMBER force fields (37, 38) in MELD and MDFF (39). In this energy space, the ensemble of structures derived from Rosetta-EM visited almost all the states of Flpp3 observed in NMR, while CryoFold recovered only a minimum number of these states at 1.8 Å resolution (Fig. S3). In contrast, for the (5.00 Å) regime, CryoFold shows a markedly better performance with predictions overlapping with the majority of NMR intermediates, as well as consistently determining lower energy structures than Rosetta-EM. Thus, extended sampling benefits of CryoFold is apparent in fuzzier data sets. Here, a broader segment of the protein folding funnel is accessed by MELD, recovering models from the poor initial guesses generated by MAINMAST(Fig. S4). Taken together, the ubiquitin and Flpp3 examples establish CryoFold as an enhanced sampling tool for resolving multiple metastable states of proteins with > 100 residues, guided only by a single experimental data set at 3-5 Å

### C. Test on soluble domains of a membrane protein with heterogeneous-resolution data

We look at the cytoplasmic domain of a large trans-membrane protein, TRPV1, a heat-sensing ion channel (592 amino acids long). The point of this test is that the data is highly heterogeneous, with experimental electron densities ranging between 3.8 to 6.0 Å (40, 41), as determined by Resmap (42). Furthermore, TRPV1 has two apo-structures deposited in the RCSB database, one with moderately resolved transmembrane helices and cytoplasmic domains(41) (pdb id:3J5P, EMDataBank: EMD-5778), and another with highly-resolved transmembrane helices (pdb id:5IRZ, EMDataBank: EMD-8118) but with the cytoplasmic regions, particularly the *β*-sheets, less resolved than in 3J5P.

CryoFold was employed to regenerate these unresolved segments of the cytoplasmic domain from the heterogeneous lower-resolution data of 5IRZ. We compare the CryoFold model to the reported 3J5P structure **(Fig. 4)**, where these domains are much better resolved showing clear patterns of *β*-strands. The final model was observed to be at an RMSD of 3.41 Å with a CC of 0.74 relative to 5IRZ. The same model with some loops removed for consistency with the EMD-5778 density produced an RMSD of 2.49 Å and CC of 0.73 with respect to 3J5P. Taken together, models derived from the CryoFold refinement of 5IRZ capture in atomistic details the highly resolved features of this density, yet without compromising with the mid-resolution cytoplasmic areas where it performs as well as the 3J5P model (Table S4).

**Fig. 4.**
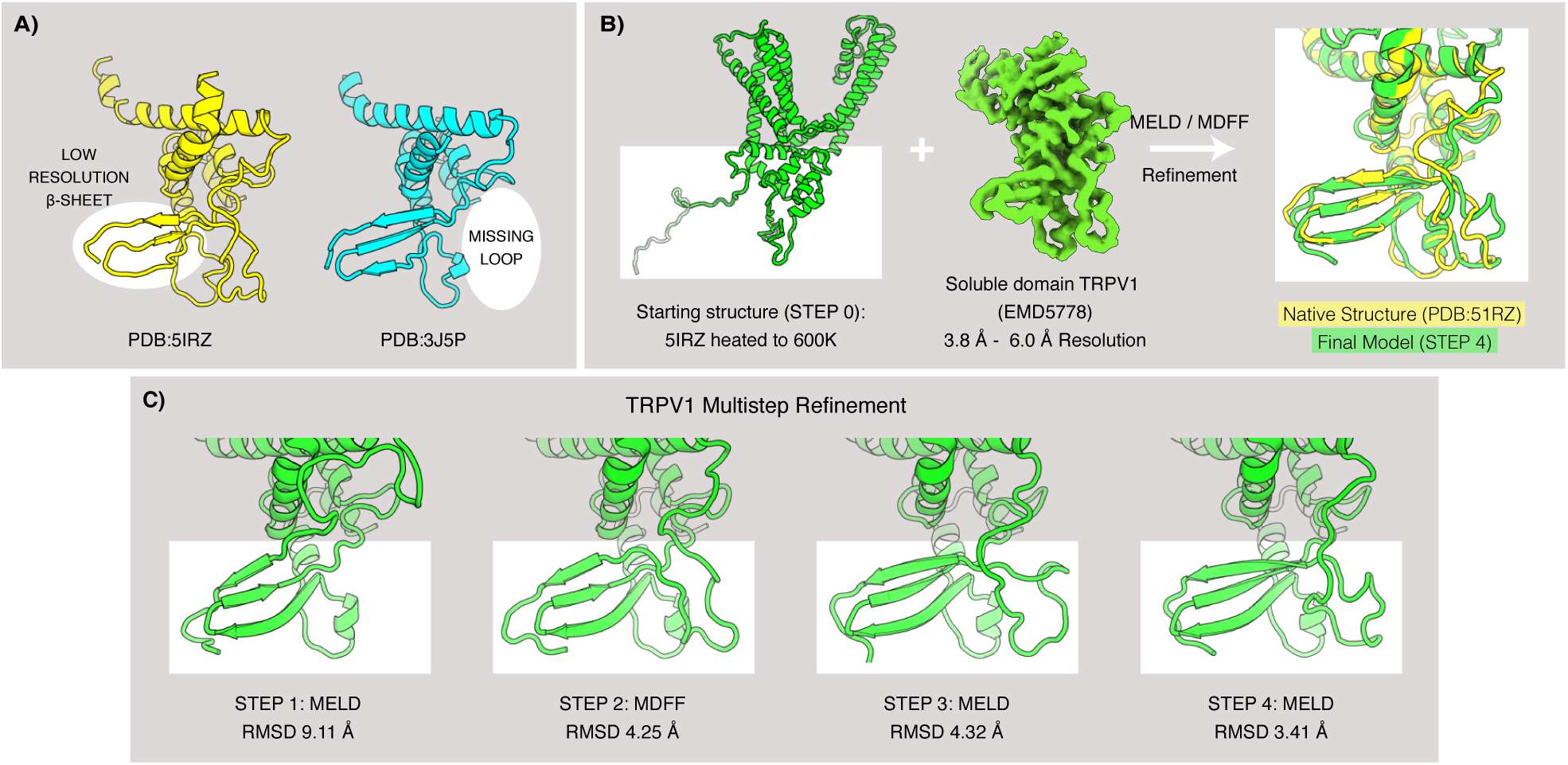
Modeling of the soluble domain of TRPV1. **(A)** TRPV1 structures deposited in 2016 (pdb 5IRZ in yellow) and in 2013 (pdb 3J5P in cyan in cartoon representation, showing the latter has a more resolved *β*-sheet while the former possess an additional extended loop. **(B)** The 5IRZ model was heated at 600 K using brute-force MD, while constraining the *α* helices. After 10 ns of simulation, this treatment resulted in a search model with the loop regions significantly deviated and the *β* sheets completely denatured. The search model was subjected to MELD-ReMDFF refinement. A single round of MELD regenerated most of the *β*-sheet from this random chain, however the 5- to 15-residue long interconnecting loops still occupied non-native positions. Subsequent ReMDFF refinement with the 5IRZ density resurrected the loop positions. One more round of the MELD and ReMDFF resulted in the further refinement of the model. The final refined model agrees well with 5IRZ **(C)** Progress of the refinement in each step of CryoFold. MELD step 1 shows the *β* sheets modeled correctly, while the loops recovered in MDFF step 2, and refinement was complete by step 4.

TRPV1 was part of the 2016 Cryo-EM modeling challenge where only ReMDFF was used(43). Presented in Table S5, our updated CryoFold model of TRPV1 (model no. 4), represents the the top - 20% of the submissions with *>* 90% Ramachandran favored statistics, and an EMRinger score of 2.54. This model is vastly refined over the originally reported structure with a score of 1.75, and our previous submission at 2.25. The improvement is attributed solely to the higher-quality *β*-sheet models that is now derived from the enhanced sampling obtained by running MELD and ReMDFF in tandem. Starting with a random coil as search model (Fig. 4B), the recovery of these *β*-sheets is highly improbable with the limited conformational space that MDFF visits. Addressing this issue, MELD invokes a multi-replica temperature exchange scheme, wherein at high replica indices it samples many distinct structures that have short lifetimes (44). At the lower-temperature replica a stronger coupling with the data is achieved, and these structures are folded into a smaller number of long-lived clusters, each with varying degrees of native contacts and secondary structure (Fig.S5). Thus, unlike MDFF, MELD allows for a search of structural motifs constrained by features in the data. When these methods are combined within CryoFold, both the backbone and sidechain geometries are refined to capture rare secondary structural changes, enabling the determination of TRPV1’s labile *β*-sheets.

An analysis of the CryoFold ensembles reveal partial unfolding of the *beta*-sheets in the soluble domains of TRPV1 with around 3-4% of the structures presenting incomplete *beta*-sheets, akin to the model originally submitted with 3J5P (Fig.S5C). Partial unfolding of these regions have not been been attributed to any functional implications in TRPV1, though some peripheral evidence of functional advantages from unfolding exist in TRPV3 channels (45). The *β*-sheets and loops from the soluble domains form the inteprotomer interface within the tertrameric channel. Secondary structural changes at these interfaces, triggers coupling between cytoplasmic and transmembrane domains, priming the channel for opening. Such changes, though rare, are indeed apparent in our MELD assignments. Therefore, the ensemble of structures and not merely a single model that CryoFold offers, opens the door to analyzing a number of distinct folded and unfolded conformations, all of which contribute to the same density map (46–48). Also evident from the TRPV1 case study, we can generate such atomistic ensembles with data of low local-resolution, yet with accuracy commensurate to structures derived from higher resolution density maps.

### D. Tests on apoferritin at three different resolutions from the 2019 EMDB modeling challenge

The EMDB competition is a community-wide effort to assess the limits of structure prediction using cryo-EM data. Here we were tasked to determine the structure of an apoferritin monomer using data at 1.8, 2.3 and 3.1 Å resolution. Following an initial tracing by MAINMAST on the monomeric map, it took two iterations for CryoFold to arrive at the final model for the first two resolutions, and three iterations for the third map. In total 17 teams participated in the 2019 competition that focused primarily on ab-initio structure determination, and all the results are reported on the EMDB website (49). CryoFold (team 73) models were independently assessed to be high accuracy (Fig. S6 (scale labeled in green)), specifically for three different categories of scores: Reference-free, EM-map and target-structure scores. The results were robust over the narrow range of resolutions tested, earning us the top rank for multiple entries (48). Comparability with respect to the target structures is almost always very high, as also reflected in commensurately high Fourier Shell Coefficient (FSC = 0.5) and cross correlations with the experimental map. Another noticeable strength is the strong EMRinger scores of the MD-based refinement, very similar to MDFF’s performance in the 2016 competition (43). A relatively new measure to evaluate mainchain geometry and to identify areas of probable secondary structure based on C-Alpha geometry, called CaBLAM (50) also found the CryoFold models to be favorable. One limitation however, is the increased number of Ramachandran outliers observed in the CryoFold and MDFF determined structures, which implicates the assumptions of classical CHARMM-type force fields(43). Our recently developed neural network potentials have already been useful to circumvent this issue (43, 46).

### E. Test on a large multi-chain protein complex with mid-resolution data

A grand challenge for cryo-EM is to determine structures of multi-chain complexes. Symmetry is used wherever possible, e.g., in viruses or homo-oligomeric membrane proteins (45, 51). However, most protein-protein or protein-nucleic acid complexes are asymmetric. Our test here is whether CryoFold could obtain the structure in an asymmetric complex. We focused on ATP synthase. It contains 31 chains. Recently Murphy et al. reported 30 distinct conformations of this motor at 2.7-4.3 Å resolution (52). Similar to the Flpp3 and TRPV1 cases, here the ensemble computed by CryoFold correctly captured the low-lying states of the multi-chain system in addition to the target 6RET conformation. For simplicity, we have removed the transmembrane *c*-ring of this system; the transmembrane challenge will be addressed in the next section.

Seven of the reported thirty models by Murphy et al. included overall deformations of the system without rotation of the *c*-ring. Using RMSD matrices (Fig. S7A), these structures were clustered in 4 distinct states (States I: 6RET; II: 6RDQ, 6RDR; III: 6RDK, 6RDL; and IV: 6RDW, 6RDX). Remarkably, all these four states are identifiable in an RMSD matrix of 220 MELD structures within CryoFold (Fig. 5B). States II, III and IV from MELD are initially at RMSD 7.6, 12.0 and 8.4 Å from 6RET respectively (Fig: S7B). After MDFF refinements, structures are consistent with experimental models from Murphy et al. listed for states II, III and IV were refined to RMSD values of 2.1, 2.8, and 1.8 Å relative to the target models (Fig. 5C, S7C and S8C). Beyond sampling the rare secondary structural changes, seen in the first four examples, here MELD visits states separated by variations in tertiary structure at the protein-protein interfaces (Fig. S9). Therefore, starting with an ensemble of structures generated to resolve 6RET, the inter-state hoping promoted by MELD’s enhanced sampling of the interface contacts (53), and refinement by ReMDFF allowed for the resolution of three more conformations of ATP synthase consistent with 6RDQ, 6RDK and 6RDW (Tables: S6 and S7).

**Fig. 5.**
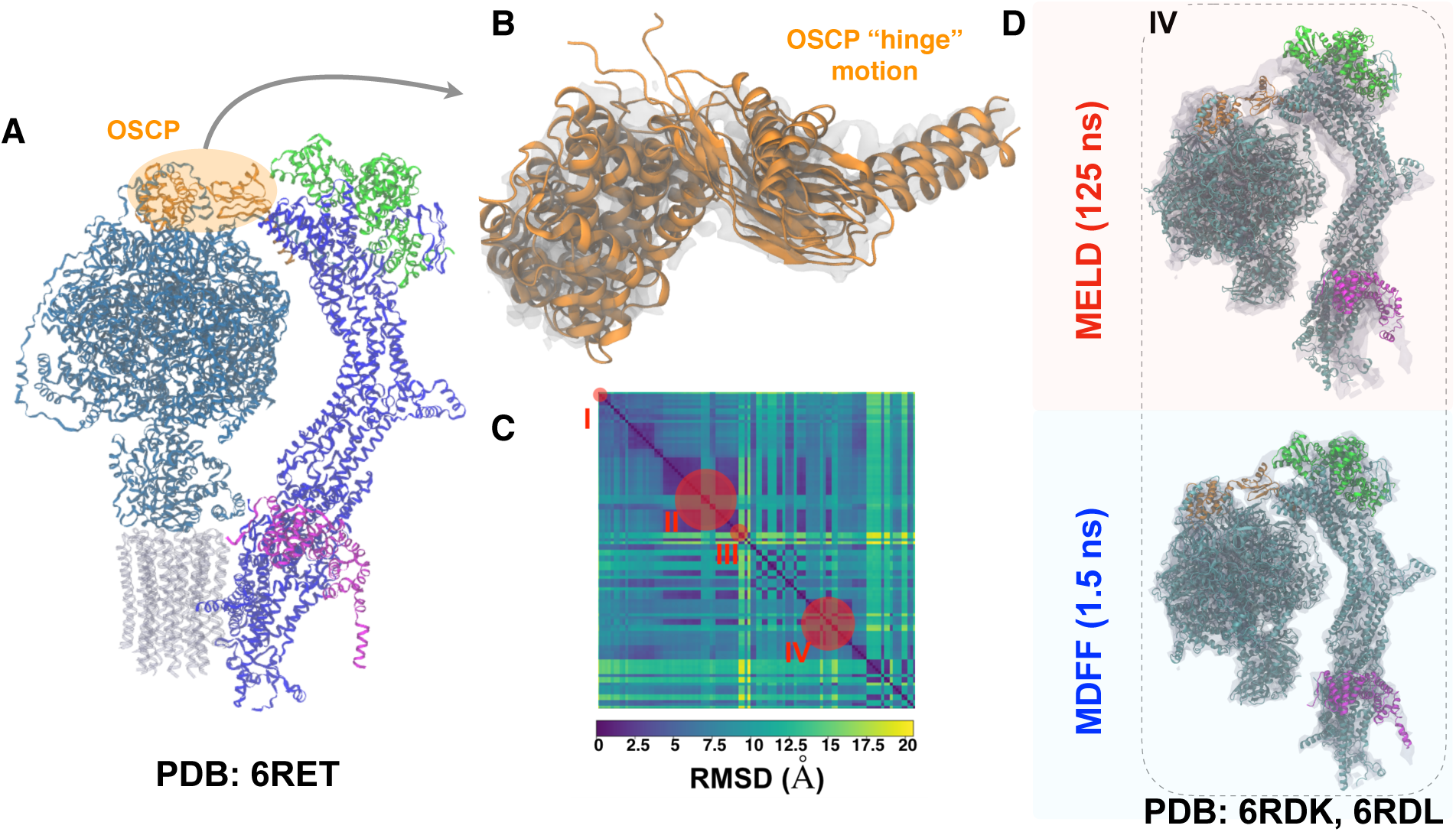
CryoFold samples several biologically relevant states of the soluble domain of mitochondrial F_1_ - F_0_ ATP-synthase. We modeled mitochondrial F_1_ - F_0_ ATPsynthase starting from pdb 6RET (state I) and excluding the grey region embedded in the membrane from refinement. CryoFold samples different conformations through a hinge motion in the OSCP region (orange) connecting the arm (blue) with the rotary domains (cyan). Clustering and 2D-RMSD analysis shows Cryofold samples conformations of additional ATPsynthase states represented by pdb codes 6RDK, 6RDL (state IV). Ohter states represented by pdb codes 6RDQ, 6RDR (state II) and 6RDW, 6RDX (state III) are included in SI.

A key biophysical outcome that we make from the CryoFold ensembles of ATP synthase is the flexibility of this motor’s peripheral stalk domains. Specifically, the OSCP hinge (chain P) assumes a number of distinct open and closed conformations with an RMSD of 3.3-6.4 Å (Fig. 5D) relative to the hinge from 6RET. The elastic coupling in ATP synthase has remained a topic of contention in the bioenergy community with crystallographers claiming minimum flexibility of the stalk regions (54), in sharp contrast to single-molecule observations of “power-strokes” that originate from deformations of the stalk (55). Within the CryoFold ensembles incorporating all the states I-IV, we see that the central stalk is in fact less flexible than the peripheral stalk with an RMSD ranging between 2.4-3.8 Å relative to 6RET. So, our results show that most of the elastic coupling in *polytomella* ATP synthase comes from the peripheral stalk, rather than the central stalk.

### F. Tests on soluble and membrane domains of a large ion channel with mid-resolution data

A second major challenge in *de novo* structure determination arises from the modeling of complete transmembrane protein systems, including structure of both the soluble and TM domains. The refinement becomes particularly daunting for CryoFold, as MELD simulations fail to capture structural changes from explicit protein-membrane interactions (44). Consequently, the accuracy of the model will depend on the structural information available from the map, and less on the fidelity of the physical interactions that underscore MELD.

Addressing this challenge, CryoFold was employed to model a monomer from the pentameric Magnesium channel CorA, containing 349 residues, at 3.80 Å resolution(56) (pdb id: 3JCF, EMDataBank: EMD-6551) **(Figs. 6)** and S10. An initial topological prediction of the channel was obtained by flexibly fitting of a linear polypeptide onto the C*α* trace obtained from the cryo-EM density using MAINMAST. These traces were already within 6.0 Å of the target C_*α*_ conformation in 3JCF, providing high-confidence coarse-grained information for MELD to operate. Leveraging the MAINMAST trace, MELD was used to perform local conformational sampling, regenerating most of the secondary structures. The model with the highest cross-correlation to the map was then refined using ReMDFF, finally resulting in models which were at 2.90 Å RMSD to the native state. Even though this model possessed high secondary structure content of 76%, substantial unstructured regions remained both in the cytoplasmic and the transmembrane regions, warranting a further round of refinement. In the subsequent MELD-ReMDFF iteration, the resulting models were 2.60 Å to the native state and agreed well with the map with a CC of 0.84. Moreover, the CryoFold models were comparable in geometry to that deposited in the database (**Fig. 6**).

**Fig. 6.**
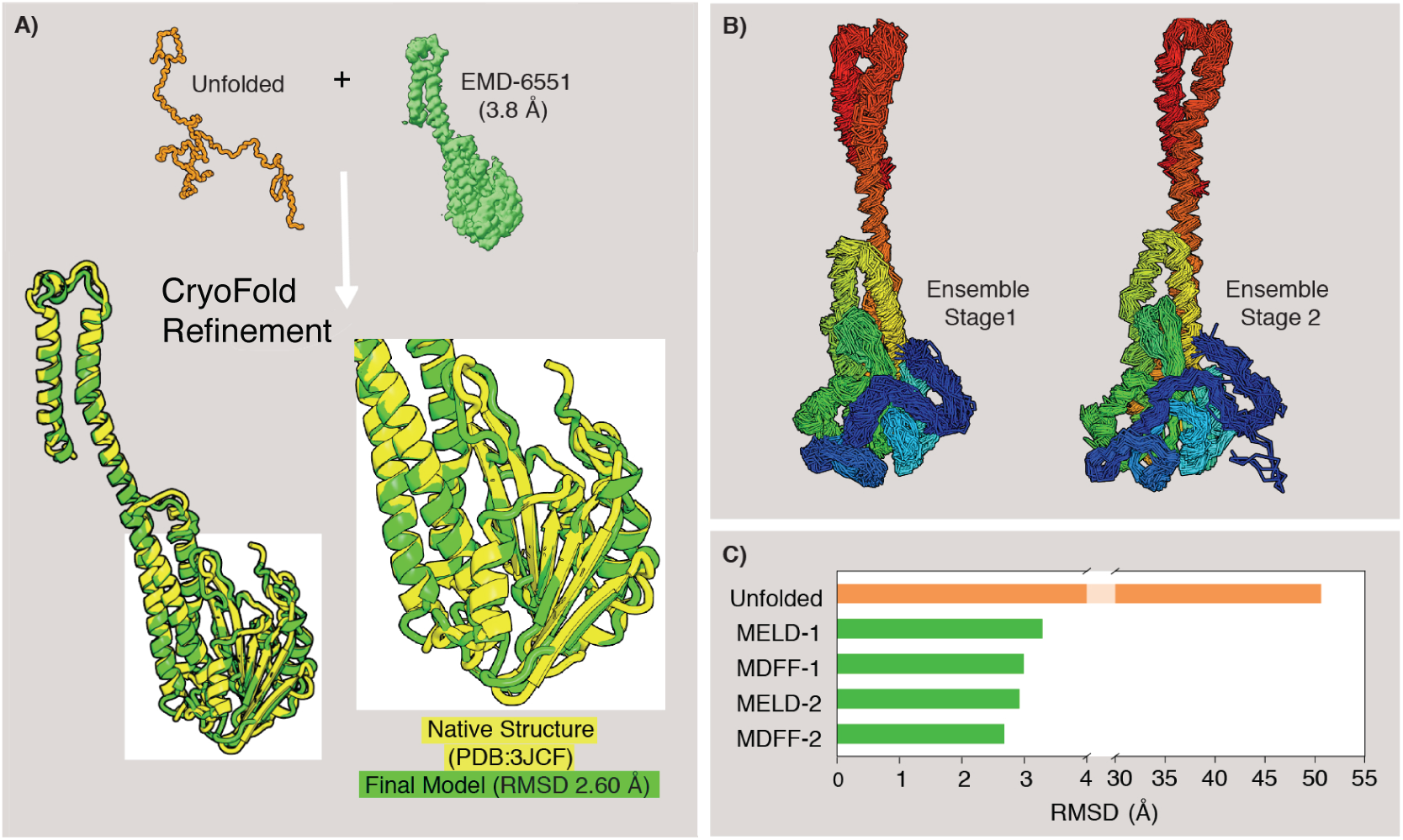
Modeling transmembrane Magnesium-channel CorA. (A) The CryoFold protocol on CorA. A starts from an C*α* trace based Cryo-EM density map using MAINMAST and refined through different cycles of MELD and MDFF produces a structure that agrees extremely well with the native structure (yellow), featuring accurate beta structures. (B) CryoFold produces narrower, more constraint ensembles as we iterate through MELD/MDFF. (C) The evolution of the RMSD of CryoFold models with each MELD-ReMDFF refinement. The end-model is 2.60 Å RMSD from the native structure.

## Discussion

The systems presented here have been chosen as challenging problems to the methods that constitute CryoFold. We have not over-optimized any aspect of the protocol to fit one problem, rather complemented the uncertainties and weakness of one method with the strengths of another. This approach is akin to the consensus methods that are known to improve performance over single methods in blind prediction challenges (57). A selected combination of methods within CryoFold’s plug-and-play protocol will enable the prediction of completely unseen data sets (Fig. S11), where the individual methods will potentially fail.

While CryoFold appears promising for obtaining biomolecular structures from cryo-EM, we are aware of some limitations. First, its success depends upon the correctness of the initial trace generated by MAINMAST. It is not clear when and whether the MD tools can recover from a wrong chain trace, particularly for resolving the transmembrane systems. Unlike Flpp3, repeating the CorA refinement with a poor-quality MAINMAST trace resulted in unreliable models. We do not have a good implicit membrane model to use in the MELD simulations and the use of explicit solvent would require many replicas, seeking more resources than currently available. Thus, by relying solely on the information coming from the density map we impose positional restraints and focus sampling on the transmembrane domains. Second, as with any MD simulation of biomolecules, the force fields are still not perfect and larger structures will be a challenge for the searching and sampling, even with an accelerator such as MELD. Finally, in our current approach, MELD is the most computationally limiting, requiring between one and ten days of sampling with 30 GPUs for the systems studied. This computational expense is not prohibitive using supercomputing resources available to academic researchers.

Despite the aforementioned limitations, CryoFold has been compared to the popular Rosetta protocols for TRPV1, ATP synthase and CorA. While for TRPV1 and CorA, Rosetta converged to models with unphysical overlap between the *β*-sheets (Fig. S5 and S12), a multi-protein refinement for ATP synthase could not be reproduced in ROSETTA-ES using standard resources, though individual chain refinements were achieved and are reported in Fig. S13. Thus, barring the Flpp3 case at 1.8 Å, CryoFold was always found to offer higher quality models, but more importantly a diverse range of structures consistent with the expected biophysics.

A key benefit of this work is the ability to capture ensembles rather than single structures. Consequently, we identify conformations that are close to the native structure but also some alternative meta-stable states that are favored by the combination of force field and data. An important question follows – are these structures really relevant or just spurious? To this end, we have now validated using NMR and cryo-EM experiments that in addition to the narrow set of models consistent with one electron density map, there exists orthogonal states that are observed both in the experiments in CryoFold refinements. These orthogonal structures sampled by MELD are indeed leveraged in biological functions, as we shown by the open → close transition in Flpp3 or flexibility of the peripheral stalks in elastic coupling of the ATP synthase example, yet behooves resolution by the limited sampling capacity of brute-force MD or MC sampling used in stationary structure determination.

Finally, evident from the 2016 and 2019 EMDB competition results, heterogeneous map resolutions affect the completeness of all the ensuing models. While a significant number of modelers prefer to truncate the more dynamic regions, MDFF offers a way to quantify uncertainty of the dynamic regions with root mean square deviations from an average model (27), and to correlate the inherent flexibility of proteins with the local resolution of density maps. Now, inside CryoFold, the fluid-like regions are even more thoroughly sampled by MELD offering the possibility of seeking hidden states in these fuzzy regions. Altogether, we present the first MD based methodology for data-guided protein folding and ensemble refinement, bridging the strengths from two distinct areas of Biophysics. The implementation is semi-automated, and manual fitting is completely avoided. However, the user will require to control the I/O between the three methods, and optimize the default parameters as required. We have provided a GUI to facilitate this stage.

## Conclusions

Structures, dynamics and function are interlinked. We often concentrate on a set of tools to determine structures from data and then use alternate computational techniques to determine dynamics between these metastable structures to ultimately elucidate biological functions. By leveraging the parallel algorithms with techniques such as CryoEM that capture multiple states (but an unknown number of them) tools that can go beyond single structures to establish molecular dynamics directly from data. CryoFold is a first step in that direction.

## Methods

The data-guided fold and fitting paradigm presented herein combines three real-space refinement methodologies, namely MELD, MAINMAST and ReMDFF. In what follows, these three formulations are articulated individually and the readers are referred to the original publications for details. Then, we outline the hybridization of the methods to provide a molecular dynamics-based *de novo* structure determination tool, CryoFold. Details of the setup for each individual system is outlined in Supplementary Information to showcase the different contexts in which CryoFold can operate.

### MELD

Modeling Employing Limited Data (MELD) employs a Bayesian inference approach (eq. Eq. (1)) to incorporate empirical data into MD simulations(15, 28). The bayesian prior 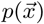 comes from an atomistic force field (ff14SB sidechain, ff99SB backbone) and an implicit solvent model (Generalized born with neck correction, gb-neck2) (37, 38). The likelihood 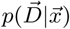, representing a bias towards known information, determines how well do the sampled conformations agree with known data, *D*. 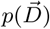 refers to the likelihood of the data, which we take as a normalization term that can typically be ignored. Taken together,

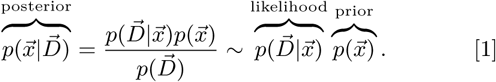

MELD is designed to handle data with one or more of these features: sparsity, noise and ambiguity. Brute-force use of such data leads to incorrect models(58) as not all the data is compatible with the native state. MELD addresses the refinement of low-resolution data by enforcing only a fraction (*x*%) of this data at every step of the MD simulation. Although *x* is kept fixed, the subset of data chosen to bias the simulation keeps changing with the simulation steps in a deterministic way. For a give nstructure all the data is evaluated, sorted according to their energy penalty and the *x*% with lowest energy guide the simulation until the next step. The data is incorporated as flat-bottom harmonic restraints *E*(*r*_*ij*_) for evaluating the likelihood 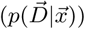.

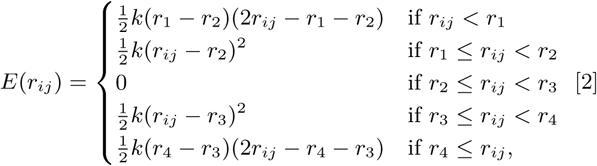

When these restraints are satisfied they do not contribute to the energy or forces, contributing for flat bottom region of eq. 2 and (Fig. S12). When the restraints are not satisfied they add energy penalties and force biases to the system – guiding it to regions that satisfy a subset of the data, or conformational envelopes. Details of MELD implementation are provided in **Supplementary methods: Description of MELD**.

### MAINMAST

MAINchain Model trAcing from Spanning Tree (MAINMAST) is a *de novo* modeling program that directly builds protein main-chain structures from an EM map of around 4-5 Å or better resolutions(26). MAINMAST automatically recognized main-chain positions in a map as dense regions and does not use any known structures or structural fragments.The procedure of MAINMAST consists of mainly four steps (Fig. S14). In the first step, MAINMAST identifies local dense points (LDPs) in an EM map by mean shifting algorithm. All grid points in the map are iteratively shifted by a gaussian kernel function and then merged to the clusters. The representative points in the clusters are called LDPs. In the second step, all the LDPs are connected by constructing a minimum spanning tree (MST). It is found that the most edges in the MST covers the main-chain of the protein structure in EM map(26). In the third step, the initial tree structure (MST) is refined iteratively by the so-called tabu search algorithm. This algorithm attempts to explore a large search space by using a list of moves that are recently considered and then forbidden. In the final step, the longest path of the refined tree is aligned with the amino acid sequence of the target protein. This process assigns optimal C*α* positions of the target protein on the path and evaluates the fit of the amino acid sequence to the longest path in a tree. Details of MAINMAST implementation are provided in **Supplementary methods: Description of MAINMAST**.

### Traditional MDFF

The protocol for molecular dynamics flexible fitting (MDFF) has been described in detail(6). Briefly, a potential map V_EM_ is generated from the cryo-EM density map, given by

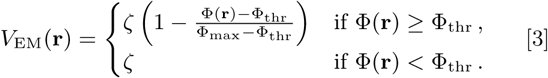

where Φ(**r**) is the biasing potential of the EM map at a point **r**, *ζ* is a scaling factor that controls the strength of the coupling of atoms to the MDFF potential, Φ_thr_ is a threshold for disregarding noise, and Φ_max_ = max(Φ(**r**)).

A search model is refined employing MD, where the traditional potential energy surface is modified by V_EM_. The density-weighted MD potential conforms the model to the EM map, while simultaneously following constraints from the traditional force fields. The output structure offers a real-space solution, resolving the density with atomistically detailed structures.

### ReMDFF

While traditional MDFF works well with low-resolution density maps, recent high-resolution EM maps have proven to be more challenging. This is because high-resolution maps run the risk of trapping the search model in a local minimum of the density features. To overcome this unphysical entrapment, resolution exchange MDFF (ReMDFF) employs a series of MD simulations. Starting with *i* = 1, the *i*th map in the series is obtained by applying a Gaussian blur of width *σ*_*i*_ to the original density map. Each successive map in the sequence *i* = 1, 2, … *L* has a lower *σ*_*i*_ (higher resolution), where *L* is the total number of maps in the series (*σ*_*L*_ = 0 Å). The fitting protocol assumes a replica-exchange approach described in details(27) and illustrated in Fig. S15. At regular simulation intervals, replicas *i* and *j*, of coordinates **x**_*i*_ and **x**_*j*_ and fitting maps of blur widths *σ*_*i*_ and *σ*_*j*_, are compared energetically and exchanged with Metropolis acceptance probability

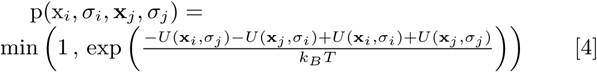

where *k*_*B*_ is the Boltzmann constant, *U* (**x**, *σ*) is the instantaneous total energy of the configuration **x** within a fitting potential map of blur width *σ*. Thus, ReMDFF fits the search model to an initially large and ergodic conformational space that is shrinking over the course of the simulation towards the highly corrugated space described by the original MDFF potential map. Details of ReMDFF implementation are provided in **Supplementary methods: Description of Resolution exchange MDFF**.

### CryoFold (MELD-MAINMAST-ReMDFF) protocol

Illustrated in **Fig. 1**, the CryoFold protocol begins with MELD computations, which guided by backbone traces from MAINMAST yields folded models. These models are flexibly fitted into the EM density by ReMDFF to generate refined atomistic structures.

1. First, information for the construction of Bayesian likelihood is derived from secondary structure predictions (PSIPRED), which were enforced with a 70% confidence. This percentage of confidence offers an optimal condition for MELD to recover from the uncertainties in secondary structure predictions(29). For membrane proteins, this number can be increased to 80% when the transmembrane motifs are well-defined helices. MELD extracts additional prior information from the MD force field and the implicit solvent model (see eq.1).
2. In the second step, any region determined with high accuracy will be kept in place with cartesian restraints imposed on the C*α* during the MELD simulations. This way, the already resolved residues can fluctuate about their initial position.
3. In the third step, distance restraints (e.g. from the C*α* traces of MAINMAST) are derived. The application of MAINMAST allows construction of pairwise interactions as MELD-restraints directly from the EM density features. Together with the cartesian restraints of step 2, these MAINMAST-guided distance restraints are enforced via flat-bottom harmonic potentials (see eq. 2) to guide the sampling of a search model; notably, the search model is either a random coil or manifests some topological features when created by fitting the coil to the C*α* trace with targeted MD. Depending upon the stage of CryoFold refinements, only a percent of the cartesian and distance restraints need be satisfied. The cartesian restraints are often localized on the structured regions, while the distance restraints typically involve regions that are more uncertain (e.g loop residues).
4. Fourth, a Temperature and Hamiltonian replica exchange protocol (H,T-REMD) is employed to accelerate the sampling of low-energy conformations in MELD(15, 28), refining the secondary-structure content of the model. The Hamiltonian is changed by changing the force constant applied to the restraints. Simulations at higher replica indexes have higher temperatures and lower (vanishing) force constants so sampling is improved. At low replica index, temperatures are low and the force constants are enforced at their maximum value (but only a certain per cent of the restraints, the ones with lower energy, are enforced). See SI for details for individual applications.
5. Fifth, cross-correlation of the H,T-REMD-generated structures with the EM-density is employed as a metric to select the best model for subsequent refinement by ReMDFF (Fig. S16). Resolution exchange across 5 to 11 maps with successively increasing Gaussian blur of 0.5 Å (*σ* in eq. 4) sufficed to improve the cross-correlation and structural statistics. The model with the highest EMringer score forms the starting point of the next round of MELD simulations. Thereafter, another round ReMDFF is initiated, and this iterative MELD-ReMDFF protocol continues until the *δ* CC between two consecutive iterations is *<*0.1.

Throughout different rounds of iterative refinement, the strucures from ReMDFF are used as seeds in new MELD simulaions. At the same time, distance restraints from the ReMDFF model are updated and the pairs of residues present in those nteractions are enforced at different accuracy levels. As expected, the more rounds of refinement we do, the higher the accuracy levels for the contacts is achieved in CryoFold. In going through this procedure, the ensembles produced get progressively narrower as we increase the amount of restraints nforced. A video tutorial and the description of this implemenation is provided in **Supplementary methods: Graphical User Interface**.

## Supporting information

Supplemental GUI

## ACKNOWLEDGMENTS

AS and CG acknowledge start-up funds from the SMS and CASD at Arizona State University, CAREER award by NSF-MCB 1942763 and the resources of the OLCF at the Oak Ridge National Laboratory, which is supported by the Office of Science at DOE under Contract No. DE-AC05-00OR22725, made available via the INCITE program. ET laboratory is supported by NIH (P41GM104601); ET, AS and MS acknowledge NIH (R01GM098243-02). This research is part of the Blue Waters sustained-petascale computing project, which is supported by the National Science Foundation (Awards OCI-0725070 and ACI-1238993) and the state of Illinois. KD and AP appreciate support from a PRAC computer allocation supported by NSF Award ACI1514873, support from NIH Grant GM125813 and the Laufer Center. AP appreciates start-up support from the University of Florida. DK acknowledges support from the National Institutes of Health (R01GM123055), the National Science Foundation (DMS1614777, CMMI1825941), and the Purdue Institute of Drug Discovery. WVH acknowledges NIH (R01GM112077).

