## Supplemental GUI for "CryoFold: determining protein structures and ensembles from cryo-EM data": gui.py

```
#!/usr/bin/env python

from kivy.core.window import Window

import sys
sys.path.insert(0, 'MAINMAST_GUI')
sys.path.insert(0, 'MELD_GUI')
sys.path.insert(0, 'TaMD_GUI')
sys.path.insert(0, 'MDFF_GUI')
sys.path.insert(0, 'Utils')

from kivy.app import App
from MainmastScreen import MainmastScreen
from MeldScreen import MeldScreen
from TaMDScreen import TaMDScreen
from TaMDScreen import Simulation_params
from MDFFScreen import MDFFScreen

from kivy.uix.gridlayout import GridLayout
from kivy.uix.boxlayout import BoxLayout

from kivy.uix.screenmanager import ScreenManager, Screen
from kivy.lang import Builder

class MenuScreen(Screen):
    pass


sm = Builder.load_string("""
<MenuScreen>:
    BoxLayout:
        padding: Window.height * 0.2
        GridLayout:
            rows: 2
            GridLayout:
                cols: 2
                BoxLayout:
                    padding: Window.height * 0.066
                    Button:
                        text: 'MAINMAST'
                        on_release:
                            root.manager.transition.direction = 'right'
                            root.manager.current = 'mainmast'
                BoxLayout:
                    padding: Window.height * 0.066
                    Button:
                        text: 'Targeted Molecular Dynamics'
                        on_release:
                            root.manager.transition.direction = 'down'
                            root.manager.current = 'tamd'
            GridLayout:
                cols: 2
                BoxLayout:
                    padding: Window.height * 0.066
                    Button:
                        text: 'MDFF'
                        on_release:
                            root.manager.transition.direction = 'up'
                            root.manager.current = 'mdff'
                BoxLayout:
                    padding: Window.height * 0.066
                    Button:
                        text: 'MELD'
                        on_release:
                            root.manager.transition.direction = 'left'
                            root.manager.current = 'meld'

ScreenManager:
    id: screen_manager
    MenuScreen:
        name: 'menu'
        manager: screen_manager
    MainmastPart:
        name: 'mainmast'
        manager: screen_manager
    MeldScreen:
        name: 'meld'
        manager: screen_manager
    TaMDPart:
        name: 'tamd'
        manager: screen_manager
    MDFFScreen:
        name: 'mdff'
        manager: screen_manager

""")


class CryoFold(App):
    def build(self):
        return sm


if __name__ == '__main__':
    CryoFold().run()
```
