## Supplemental GUI for "CryoFold: determining protein structures and ensembles from cryo-EM data": install_linux.sh

```
#!/bin/bash

echo "Compilation of Mainmast and ThreadCA"
gfortran ./MAINMAST_GUI/MainmastThreadCA/MAINMAST.f -w -O3 -fbounds-check -o ./MAINMAST_GUI/MainmastThreadCA/MAINMAST -mcmodel=medium
gfortran ./MAINMAST_GUI/MainmastThreadCA/ThreadCA.f -w -O3 -fbounds-check -o ./MAINMAST_GUI/MainmastThreadCA/ThreadCA -mcmodel=medium
echo "Compilation done"
conda config --add channels maccallum_lab
conda config --add channels omnia
conda config --add channels conda-forge
yes | conda create -n cryofold python=3.5 kivy numpy mdtraj meld-cuda75
echo "Anaconda virtual environment created"
chmod +x ./gui.py
chmod +x start.sh
echo "Installation done"
```
