## Supplemental GUI for "CryoFold: determining protein structures and ensembles from cryo-EM data": MainmastScreen.py

```
from kivy.uix.boxlayout import BoxLayout
from kivy.uix.gridlayout import GridLayout
from kivy.event import EventDispatcher
from kivy.properties import ObjectProperty, StringProperty
from kivy.uix.popup import Popup
from kivy.lang import Builder
from LoadDialog import LoadDialog
from AlertDialog import AlertDialog
from kivy.uix.screenmanager import ScreenManager, Screen, SlideTransition
from kivy.uix.button import Button
from kivy.uix.textinput import TextInput
from kivy.uix.label import Label
from kivy.uix.checkbox import CheckBox

import os
import subprocess


Builder.load_file('./MAINMAST_GUI/MainmastScreenManager.kv')

Builder.load_file('./MAINMAST_GUI/mainmastscreen.kv')
Builder.load_file('./MAINMAST_GUI/threadca.kv')


class MainmastPart(Screen):
    pass


class GeneralScreen(Screen):

    def dismiss_popup(self):
        self._popup.dismiss()

    def show_load(self, fileinput):
        content = LoadDialog(load=self.load, cancel=self.dismiss_popup, fileinput=fileinput)
        self._popup = Popup(title="Load file", content=content,
                            size_hint=(0.9, 0.9))
        self._popup.open()

    def show_alert(self, message):
        content = AlertDialog(close=self.dismiss_popup, message=message)
        self._popup = Popup(title="Alert", content=content,
                            size_hint=(0.5, 0.2))
        self._popup.open()

    def load(self, path, filename, fileinput):
        with open(os.path.join(path, filename[0])) as stream:
            getattr(self, fileinput).text = stream.name
        self.dismiss_popup()

    def close():
        self.dismiss_popup()

class MainmastScreen(GeneralScreen):

    def check_file(self):
        format = self.file_name.text.split('.')[-1];
        if format != 'situs':
            self.show_alert('Wrong Format')
        else:
            self.manager.transition = SlideTransition(direction='left')
            self.manager.current = 'threadca'

    def on_leave(self, *largs):
        if self.gw.text == '':
            self.gw.text = '2.0'
        if self.d_keep.text == '':
            self.d_keep.text = '0.6'
        if self.t.text == '':
            self.t.text = '0'
        if self.allow.text == '':
            self.allow.text = '10.0'
        if self.filter.text == '':
            self.filter.text = '0.1'
        if self.merge.text == '':
            self.merge.text = '0.6'
        if self.n_round.text == '':
            self.n_round.text = '5000'
        if self.n_nb.text == '':
            self.n_nb.text = '30'
        if self.n_tb.text == '':
            self.n_tb.text = '100'
        if self.r_local.text == '':
            self.r_local.text = '10'
        if self.const.text == '':
            self.const.text = '1.01'
        setattr(self.manager.statedata, 'file_name', self.file_name.text)
        setattr(self.manager.statedata, 'gw', self.gw.text)
        setattr(self.manager.statedata, 'd_keep', self.d_keep.text)
        setattr(self.manager.statedata, 't', self.t.text)
        setattr(self.manager.statedata, 'allow', self.allow.text)
        setattr(self.manager.statedata, 'filter', self.filter.text)
        setattr(self.manager.statedata, 'merge', self.merge.text)
        setattr(self.manager.statedata, 'n_round', self.n_round.text)
        setattr(self.manager.statedata, 'n_nb', self.n_nb.text)
        setattr(self.manager.statedata, 'n_tb', self.n_tb.text)
        setattr(self.manager.statedata, 'r_local', self.r_local.text)
        setattr(self.manager.statedata, 'const', self.const.text)

class ThreadCAScreen(GeneralScreen):

    def change_button_run_name(self, widget, message):
        widget.text = message

    def run_command(self, widget):
        if (
            self.manager.statedata.file_name.split('.')[-1] != 'situs' or
            self.file_name_spider2.text.split('.')[-1] != 'spd3'
            ):
            self.show_alert('Wrong Format')
        else:
            if self.reverse.active:
                ca_file = "./outputs/CA_r.pdb"
                reverse = '-r'
            else:
                ca_file = "./outputs/CA.pdb"
                reverse = ''
            try:
                mainmast = open("./outputs/tmp/path.pdb","wb")
                threadca = open(ca_file,"wb")
                subprocess.check_call([
                                './MAINMAST_GUI/MainmastThreadCA/MAINMAST',
                                '-m',
                                self.manager.statedata.file_name,
                                '-gw',
                                self.manager.statedata.gw,
                                '-Dkeep',
                                self.manager.statedata.d_keep,
                                '-t',
                                self.manager.statedata.t,
                                '-allow',
                                self.manager.statedata.allow,
                                '-filter',
                                self.manager.statedata.filter,
                                '-Nround',
                                self.manager.statedata.n_round,
                                '-Ntb',
                                self.manager.statedata.n_tb,
                                '-Rlocal',
                                self.manager.statedata.r_local
                                ], stdout=mainmast)
                subprocess.check_call([
                                './MAINMAST_GUI/MainmastThreadCA/ThreadCA',
                                '-i',
                                './outputs/tmp/path.pdb',
                                '-a',
                                '20AA.param',
                                '-spd',
                                self.file_name_spider2.text,
                                reverse
                                ], stdout=threadca)
                mainmast.close()
                threadca.close()
                self.parent.parent.manager.transition = SlideTransition(direction='right')
                self.manager.transition = SlideTransition(direction='right')
                self.manager.current = 'mainmast'
                self.parent.parent.manager.current = 'menu'
            except:
		
                self.show_alert('Error!')
        widget.text = 'Run'


class MainmastState(EventDispatcher):
        file_name = ''
        gw = ''
        d_keep = ''
        t = ''
        allow = ''
        filter = ''
        merge = ''
        n_round = ''
        n_nb = ''
        n_tb = ''
        r_local = ''
        const = ''

class StateManager(ScreenManager):
    statedata = ObjectProperty(MainmastState())
```
