## Supplemental GUI for "CryoFold: determining protein structures and ensembles from cryo-EM data": MAINMAST.f

```
C       ***************************************************************
c
c	MAINMAST: Mainchain Model Tracing From Spanning Tree
c
C       ***************************************************************
c
c	INPUT: situs format file
c
c
c	#COPYRIGHT
c	(c) 2017 Genki Terashi, Daisuke Kihara and Purdue University
c
c	MAINMAST is a free software for academic and non-commercial users. It is released under the terms of the GNU General Public License Ver.3 (https://www.gnu.org/licenses/gpl-3.0.en.html).
c	Commercial users please contact for alternate licensing.
c 	Citation of the following reference should be included in any publication that uses data or results generated by MAINMAST program.
c	(a) Genki Terashi1, and Daisuke Kihara. De novo main-chain modeling for EM maps using MAINMAST. (in submission)
c


	IMPLICIT INTEGER(I-Z)
        character*5 struct(5)
	character*3 aa(-1:20), NAME

	PARAMETER(NDIM=800)
	PARAMETER(NREPS=50)
	PARAMETER(MSIZE=300)
	PARAMETER(MAXB=5)

	dimension ftmp_xyz(3),xyz_t(3,ndim),xyz_u(3,ndim),xyz_o(3,ndim)
	DIMENSION reps_xyz(3,ndim,NREPS)
	DIMENSION TMP_XYZ(3),AS_XYZ(3),AM_XYZ(3),A_ROT(3,3),I_MAT(3,3)
	COMMON /MAP_DATA/  gstep,gbase(3),NumOfGrid(3),dreso,
     *          dmap(MSIZE,MSIZE,MSIZE),amap(MSIZE,MSIZE,MSIZE),
     *          cmap(MSIZE,MSIZE,MSIZE), !raw data
     *          obox(3),pbox(3),list(NDIM),Nlist,lbase(3),lbase1(3)
	DIMENSION buff(10),add_f(0:23),best_f(0:23)
	DIMENSION add_w(0:23),best_w(0:23)
	DIMENSION gbase1(3),NumOfGrid1(3), gbase2(3),NumOfGrid2(3)
	!For Mutual Information
	COMMON /MI/	bins(21),Nvox,Npos,Nmap(21),Nmod(21)
	!DIMENSION Emap(MSIZE,MSIZE,MSIZE)
	REAL fin(MSIZE*MSIZE*MSIZE),fori(MSIZE*MSIZE*MSIZE)
        INTEGER order(MSIZE*MSIZE*MSIZE),posi(3,MSIZE*MSIZE*MSIZE)
	DIMENSION After(3,MSIZE*MSIZE*10) !After shift
	DIMENSION Before(3,MSIZE*MSIZE*10) !After shift
	DIMENSION bins1(21),bins2(101),path(MSIZE*MSIZE)
	DIMENSION Nx(21),Ny(101),Nxy(21,101)

	COMMON /GRAPH/ Nbranch(MSIZE*MSIZE*10),IDb(MSIZE*MSIZE*10,MAXB*2),
     *	edged(MSIZE*MSIZE*10,MAXB*2),
     *		nextp(MSIZE*MSIZE*10),
     *		cost(MSIZE*MSIZE*10),
     *  id_edge(MSIZE*MSIZE*10,MAXB*2)
	DIMENSION member(MSIZE*MSIZE*10)
	!TMalign rmsd.f
	double precision fitrms,
     *	fx(3,ndim),fy(3,ndim),fw(ndim),fu(3,3),ft(3)
	COMMON /KRU/ Nround,Nlocal,Ntabu,allow_rate_max,dlocal,dside,mode
	!for command
	character*500 arg(20),fname_pdb,fname_map,buf;
	INTEGER vpath(MSIZE*MSIZE*10),best_path(10,MSIZE*MSIZE*10),
     *  best_Nv(10)

C
c	Command line ver
c
c
!Default value
	Nres=327;
	mode=0;
	dreso=2.0
	fminD=0.5
	control=0.0;
	allow_shift=10.0;
	af_mg_cut=0.1
	af_mg_r=0.5;
	Nround=5000;
	Nlocal=30;
	Ntabu=100;
	dlocal=10.0;
	allow_rate_max=1.01;
	!get command line options
	if(iargc().ge.2) then
	 do i=1,iargc()
	  call getarg(i,arg(i))
	  !read(arg(i),*),buf;
	  buf=arg(i);
	  !write(*,*),buf;
	  if(buf(1:5).eq."-Tree")then
	   write(*,*)"#Show MST mode";
	   mode=1;
           cycle;
          endif
	  if(buf(1:6).eq."-Graph")then
	   write(*,*)"#Show Fully Connected Graph mode";
	   mode=2;
           cycle;
          endif
	  if(buf(1:2).eq."-m ")then
	   call getarg((i+1),arg(i+1))
	   fname_map=arg(i+1);
	   write(*,*),"#MAP=",trim(fname_map)
	   cycle;
	  endif
	  if(buf(1:3).eq."-gw")then
           call getarg(i+1,arg(i+1))
           read(arg(i+1),*),dreso;
           cycle;
          endif
	  if(buf(1:6).eq."-Dkeep")then
           call getarg(i+1,arg(i+1))
           read(arg(i+1),*),fminD;
           cycle;
          endif
	  if(buf(1:2).eq."-t")then
           call getarg(i+1,arg(i+1))
           read(arg(i+1),*),control;
           cycle;
          endif
	  if(buf(1:6).eq."-allow")then
           call getarg(i+1,arg(i+1))
           read(arg(i+1),*),allow_shift;
           cycle;
          endif
	  if(buf(1:7).eq."-filter")then
           call getarg(i+1,arg(i+1))
           read(arg(i+1),*),af_mg_cut;
           cycle;
          endif
	  if(buf(1:6).eq."-merge")then
           call getarg(i+1,arg(i+1))
           read(arg(i+1),*),af_mg_r;
           cycle;
          endif
	  if(buf(1:7).eq."-Nround")then
           call getarg(i+1,arg(i+1))
           read(arg(i+1),*),Nround;
           cycle;
          endif
	  if(buf(1:4).eq."-Nnb")then
           call getarg(i+1,arg(i+1))
           read(arg(i+1),*),Nlocal;
           cycle;
          endif
	  if(buf(1:4).eq."-Ntb")then
           call getarg(i+1,arg(i+1))
           read(arg(i+1),*),Ntabu;
           cycle;
          endif
	  if(buf(1:7).eq."-Rlocal")then
           call getarg(i+1,arg(i+1))
           read(arg(i+1),*),dlocal;
           cycle;
          endif
	  if(buf(1:6).eq."-Const")then
           call getarg(i+1,arg(i+1))
           read(arg(i+1),*),allow_rate_max;
           cycle;
          endif
	 enddo
	else
	 write(*,*),"Usage: MAINMAST "
     *   ,"-m [MAP file (situs format)]"
     *   ," (option)"
	 write(*,*),"Option ver2.0:"
	 write(*,*)," -Tree   : Show MSTree mode"
	 write(*,*)," -Graph  : Graph mode"
	 write(*,*),"---Parameters in MeanShift----"
	 write(*,*)," -gw    [f] : bandwidth of the gaussian filter"
	 write(*,*),"               def=2.0, sigma = 0.5*[float]"
	 write(*,*)," -Dkeep [f] : Keep edge where distance < [f] def=0.5"
	 write(*,*)," -t     [f] : Threshold of density values. def=0.0"
	 write(*,*)," -allow [f] : Max shift distance < [f] def=10.0"
         write(*,*)," -filter [f]: Filter of representative points"
	 write(*,*),"               def=0.1"
         write(*,*)," -merge [f]: After MeanShifting, merge d<[f]"
	 write(*,*),"               def=0.5"
	 write(*,*),"---Parameters in Tabu-search----"
         write(*,*)," -Nround [i]: Number of Iterations"
	 write(*,*),"               def=5000"
	 write(*,*)," -Nnb   [i]: Number of Neighborss"
	 write(*,*),"               def=30"
	 write(*,*)," -Ntb   [i]: Size of tabu-list"
	 write(*,*),"               def=100"
	 write(*,*)," -Rlocal [f]: Radius of Locat MST"
	 write(*,*),"               def=10"
	 write(*,*)," -Const  [f]: Constraint of total length of edge"
	 write(*,*),"               def=1.01,Total(Tree) <[f]*Total(MST)"
	 stop
	endif


	write(*,*),"#Gaussian Width",dreso
	write(*,*),"#Dkeep",fminD;
	write(*,*),"#Threshold of Density value",control;
	write(*,*),"#Allow shift",allow_shift;
	write(*,*),"#Nround",Nround;
	write(*,*),"#N neighbors",Nlocal;
	write(*,*),"#Ntabu",Ntabu;
	write(*,*),"#MG_R=",af_mg_r
	write(*,*),"#MG_CUT=",af_mg_cut
	write(*,*),"#R_LOCAL=",dlocal
	write(*,*),"#Const",allow_rate_max;


	!Reading MAP data
	!OPEN(UNIT=44,FILE='MAP',STATUS='OLD') !New!!!
	OPEN(UNIT=44,FILE=fname_map,STATUS='OLD') !New!!!
c	Reading MAP...............
        read(44,*) gstep,(gbase(k),k=1,3),(NumOfGrid(k),k=1,3)
        write(*,*) '#ReadingMap..'
	write(*,*),"#GridSize=",gstep
	write(*,*),"#BaseCoordinates=",gbase(:)
	write(*,*),"#Number of Grids=",NumOfGrid(:)

        !check size
        if(maxval(NumOfGrid(:)).gt.MSIZE)then
         write(*,*) 'ERROR !! Size of MAP > MSIZE !!'
         stop
        endif

        MaxLines=NumOfGrid(1)*NumOfGrid(2)*NumOfGrid(3)/10
        write(*,*) '#MaxLines=',MaxLines
	read(44,*) ! for space
        i=1;j=1;k=1;SpaseN=0;
        do l=1,MaxLines
         read(44,*) (buff(m),m=1,10)
         do m=1,10
	  if(buff(m).gt.0)then !Ignore negatve value
           amap(i,j,k)=buff(m)
	  endif
          if(buff(m).gt.control) then
	   SpaseN=SpaseN+1
	   posi(1:3,SpaseN)=(/i,j,k/)
	   fin(SpaseN)=buff(m)
	  endif
          i=i+1
          if(i.gt.NumOfGrid(1)) then
           i=1
           j=j+1
          endif
          if(j.gt.NumOfGrid(2)) then
           j=1
           k=k+1
          endif
         enddo
        enddo
        !for remain lines......
        remain=NumOfGrid(1)*NumOfGrid(2)*NumOfGrid(3)-MaxLines*10
        if(remain.gt.0) then
        read(44,*) (buff(m),m=1,remain)
        do m=1,remain
	 if(buff(m).gt.0)then !Ignore negatve value
          amap(i,j,k)=buff(m)
	 endif
	 if(buff(m).gt.control) then
	   SpaseN=SpaseN+1
	   posi(1:3,SpaseN)=(/i,j,k/)
	   fin(SpaseN)=buff(m)
	  endif

         i=i+1
         if(i.gt.NumOfGrid(1)) then
          i=1
          j=j+1
         endif
         if(j.gt.NumOfGrid(2)) then
          j=1
          k=k+1
         endif
        enddo
        endif
        close(44)

	write(*,*) "#GridPoints= ",SpaseN
	!Upsampling
	if(gstep.gt.1.0)then
	 write(*,*),"#Grid Step",gstep," >1.0";
	 call conv_amap_double();
	 !gstep=1.0;
	 amap=dmap;
	 SpaseN=0
	 do i=1,NumOfGrid(1)
	  do j=1,NumOfGrid(2)
	   do k=1,NumOfGrid(3)
	    if(amap(i,j,k).gt.control) then
	     SpaseN=SpaseN+1
	     posi(1:3,SpaseN)=(/i,j,k/)
	     fin(SpaseN)=amap(i,j,k)
	    endif
	   enddo
	  enddo
	 enddo
	write(*,*) "#GridPoints(Upsampling)= ",SpaseN
	endif

	Nbf=SpaseN
	fori(1:SpaseN)=fin(1:SpaseN);

	fin=0;
	do i=1,SpaseN
	 !Mean Shift
	 call MeanShift(float(posi(1:3,i)),after(1:3,i),fin(i),
     *   allow_shift/gstep)
	enddo
	!Merge points

	if(.false.) then
	fmax=maxval(fin)
	Natm=1
	write(*,'("MODEL    2")')
	do kk=1,SpaseN
	 ftmp_xyz=(posi(:,kk)-1)*gstep+gbase;
	 write(*,'("HETATM",I5,"  O   HOH",I6,"    ",3f8.3,f6.2,f6.2)'),
     *   Natm,Natm,ftmp_xyz,1.00,fin(kk)/fmax
	 Natm=Natm+1
	enddo
	write(*,'("ENDMDL")')
	stop
	endif
	Before(1:3,1:SpaseN)=after(1:3,1:SpaseN);
	call MergePoints(after(1:3,1:SpaseN),SpaseN,fin,
     *  af_mg_r/gstep,af_mg_cut,member)


	vflag=0
	if(mode.eq.1.or.mode.eq.2)then
	fmax=maxval(fin)
	Natm=1
	write(*,'("MODEL    3")')
	do kk=1,SpaseN
	 ftmp_xyz=(after(:,kk)-1)*gstep+gbase;
	 write(*,'("ATOM  ",I5,"  CA  ALA A",I4,"    ",3f8.3,f6.2,f6.2)'),
     *   Natm,Natm,ftmp_xyz,1.00,fin(kk)/fmax
	 Natm=Natm+1
	enddo
	!stop
	endif

	!Set up graph
	!Generate Tree
	dlocal=dlocal/gstep
	call Kruskal(after(1:3,1:SpaseN),SpaseN,fin,fminD/gstep,
     *  posi(1:3,1:Nbf),Nbf,member,path,Np,best_path,best_Nv)


	if (Np.gt.0) then
c	Put CA
	Natm=1;
	fmax=maxval(fin(1:SpaseN));
	!Convert real space
	do ii=1,SpaseN
	 after(1:3,ii)=(after(1:3,ii)-1.00)*gstep+gbase;
	enddo
	do ii=1,Nbf
	 Before(1:3,ii)=(Before(1:3,ii)-1.00)*gstep+gbase;
	enddo


	do ppp=1,10
	 if(best_Nv(ppp).gt.1) then
	 call ca_trace_kmean(after(1:3,1:SpaseN),SpaseN,
     *  Before(1:3,1:Nbf),Nbf,fori,fin,
     *  best_path(ppp,1:best_Nv(ppp)),best_Nv(ppp),Nres)
	 endif
	enddo
	endif
	END

	SUBROUTINE CA_TRACE_KMEAN(af,n,bf,Nbf,fori,fin,path,Np,Nres)
	IMPLICIT INTEGER(I-Z)
	PARAMETER(MSIZE=300)
	PARAMETER(MAXB=5)
	DIMENSION af(3,MSIZE*MSIZE*10),fin(MSIZE*MSIZE*MSIZE)
	DIMENSION path(MSIZE*MSIZE*10),ftmp(3)
	DIMENSION bf(3,MSIZE*MSIZE*10)
	REAL fori(Nbf),cog(3,Nres),sumcd(3,Nres),sumw(0:Np+1)
	INTEGER gp(Np),Ngp(Np);
	LOGICAL used(Np)
	DIMENSION path_use(Nres);
	REAL smtx(0:Np,0:Nres),sumpath(Np)
	INTEGER direc(0:Np,0:Nres),Pali(Nres);

	gp=0;
	Ngp=0;

	!Show main path
	fmax=maxval(fin(1:n))

	sumw=0;
	ftotal=sum(fori(1:Nbf));
	AveSum=ftotal/float(Nres)
	do ii=1,Nbf
	 fmin=10000000;
	 minid=0;
	 do kk=1,Np
	  ftmp=bf(1:3,ii)-af(1:3,path(kk))
	  d=dot_product(ftmp,ftmp)
	  if(d.lt.fmin)then
	   fmin=d;
	   minid=kk;
	  endif
	 enddo
	 sumw(minid)=sumw(minid)+fori(ii);
	enddo

	!Split per residue
	rcnt=1;
	tmpsum=0;
	do i=1,Np
	 tmpsum=tmpsum+sumw(i);
	 gp(i)=rcnt;
	 if(tmpsum.gt.rcnt*AveSum)then
	  rcnt=rcnt+1;
	 endif
	enddo

	!Show
	Natm=1
	fmax=maxval(sumw(1:Np))
	do ii=1,Np
	 kk=path(ii);
	 ftmp=af(:,kk);
	 write(*,'("ATOM  ",I5,"  CA  ALA ",I5,"    ",3f8.3,f6.2,f6.2)'),
     *   Natm,Natm,ftmp,1.00,sumw(ii)/fmax
	 Natm=Natm+1
	enddo
	write(*,'("ENDMDL")')
	END

	SUBROUTINE KRUSKAL(af,n,fin,cut,bf,Nbf,member,Plgst,Nlgst,
     *  best_path,best_Nv)
	IMPLICIT INTEGER(I-Z)
	PARAMETER(MSIZE=300)
	PARAMETER(MAXB=5)
	PARAMETER(MAX_TABU=100)
	DIMENSION af(3,MSIZE*MSIZE*10),fin(MSIZE*MSIZE*MSIZE),
     *   member(MSIZE*MSIZE*10)
	INTEGER bf(3,MSIZE*MSIZE*10)
	COMMON /GRAPH/ Nbranch(MSIZE*MSIZE*10),IDb(MSIZE*MSIZE*10,MAXB*2),
     *	edged(MSIZE*MSIZE*10,MAXB*2),
     *		nextp(MSIZE*MSIZE*10),
     *		cost(MSIZE*MSIZE*10),
     *  id_edge(MSIZE*MSIZE*10,MAXB*2)
	DIMENSION Nbranch_tmp(MSIZE*MSIZE*10),
     *  IDb_tmp(MSIZE*MSIZE*10,MAXB*2),
     *  edged_tmp(MSIZE*MSIZE*10,MAXB*2),
     *  id_edge_tmp(MSIZE*MSIZE*10,MAXB*2)
	COMMON /FLAGS/ vflag
	DIMENSION ftmp(3),cd_st(3),cd_ed(3),itmp(3),fvec(3)
	LOGICAL activeg(MSIZE*MSIZE*10),activeg_tmp(MSIZE*MSIZE*10),
     *          activeg_ori(MSIZE*MSIZE*10)
	DIMENSION list_act(MSIZE*MSIZE) !active list
	DIMENSION tmp_list(MSIZE*MSIZE)
	DIMENSION path(MSIZE*MSIZE),Plgst(MSIZE*MSIZE);
	LOGICAL flag,flag1,flag2,flag3
	INTEGER cid(n),Nc(n),con_cnt(n,n)
	DIMENSION edge(n*n),ide(2,n*n),order(n*n),list_bad(n*n),
     *  Tree(n*n),edge_tmp(n*n),edge_best(n*n),edge_dens(n*n),
     *  TreeKth(n*n,100),NtKth(100),FtotalKth(100),
     *  PathKth(n*n,100),NpKth(100),tmp_IDb(MSIZE*MSIZE*10,MAXB*2),
     *  TreeNow(n*n),TreeBest(n*n),TreeRoundBest(n*n),
     *  path_bad(n*n),id2order(n*n),tmp_tree(n*n),
     *  id2orderNow(n*n),id2orderTmp(n*n),id2orderRbest(n*n)
	real sco(100),best_sco,round_best_sco,now_sco
	DIMENSION ope(4),TabuList(2,MSIZE),TabuListTmp(2,MSIZE)
	COMMON /KRU/ Nround,Nlocal,Ntb,allow_rate_max,dlocal,dside,mode
	DIMENSION EdgeTable(MSIZE*MSIZE)
	LOGICAL mstv(n),local_edge(MSIZE*MSIZE*10),used_v(n);
	INTEGER vpath(n),best_path(10,MSIZE*MSIZE*10),best_Nv(10)
	DIMENSION best_trj_sco(10),TmpPath(MSIZE*MSIZE*10);
	DIMENSION PathNow(MSIZE*MSIZE*10),PathRoundBest(MSIZE*MSIZE*10);
	!check area contact
	con_cnt=0
	do ii=1,Nbf
	 if(member(ii).eq.0) cycle
	 do jj=ii+1,Nbf
	  if(member(jj).eq.0) cycle
	  if(member(ii).eq.member(jj))cycle
	  itmp=abs(bf(1:3,ii)-bf(1:3,jj))
	  if(maxval(itmp(1:3)).eq.1)then
	   con_cnt(member(ii),member(jj))=
     *     con_cnt(member(ii),member(jj))+1
	   con_cnt(member(jj),member(ii))=
     *     con_cnt(member(jj),member(ii))+1
	  endif
	 enddo
	enddo
	!write(*,*),"CON","163-246",con_cnt(163,246)
	if(Ntb.gt.MAX_TABU)Ntb=MAX_TABU

	!init cluster id
	!sort edge
	Ne=0
	dl2=dl*dl
	du2=du*du
	do ii=1,n
	 cid(ii)=ii
	 do jj=ii+1,n
	  if(con_cnt(ii,jj).eq.0) cycle
	  ftmp=af(:,ii)-af(:,jj)
	  d=sqrt(dot_product(ftmp,ftmp))
	  Ne=Ne+1
	  edge(Ne)=d
	  ide(1,Ne)=ii;
	  ide(2,Ne)=jj;
	 enddo
	enddo
	write(*,*),"#Ne=",Ne
	edge_tmp=edge
	call quick_sort(edge,order,Ne)

	!Find MST using all edges
	 !init cid
	 do ii=1,n;cid(ii)=ii;enddo
	Nt=0
	 do ii=1,Ne
	  !minimum edge
	  Me=order(ii)
	  if(Me.eq.rmid) cycle
	  v1=ide(1,Me)
	  v2=ide(2,Me)
	  if(cid(v1).ne.cid(v2)) then
	   !Add edge ii to Tree
	   Nt=Nt+1
	   Tree(Nt)=Me
	   !Merge cluster
	   tmpid=cid(v2)
	   do jj=1,n
	    if(cid(jj).eq.tmpid) cid(jj)=cid(v1)
	   enddo
	  endif
	 enddo

	!cleaning remove island data
	Nc=0
	do ii=1,n
	 Nc(cid(ii))=Nc(cid(ii))+1
	enddo
	Maxc=maxval(Nc)
	do ii=1,n
	 if(Nc(ii).eq.Maxc)then
	  UseChain=ii
	  exit
	 endif
	enddo
	tmp_nt=Nt
	Nt=0
	do ii=1,tmp_nt
	 if(UseChain.eq.cid(ide(1,Tree(ii))))then
	 Nt=Nt+1
	 Tree(Nt)=Tree(ii) !shift
	 endif
	enddo
	mstv(1:n)=.false.
	!sort edge again-----------------
	Ne=0
	do ii=1,n
	 if(cid(ii).ne.UseChain) cycle
	 do jj=ii+1,n
	  !if(.not.con_flag(ii,jj)) cycle
	  if(con_cnt(ii,jj).eq.0) cycle
	  if(cid(jj).ne.UseChain) cycle
	  ftmp=af(:,ii)-af(:,jj)
	  d=sqrt(dot_product(ftmp,ftmp))
	   Ne=Ne+1
	   edge(Ne)=d !distance
	   !!!!!!!!!!
	   edge_dens(Ne)=(fin(ii)+fin(jj))*0.5; !Average Density
	   !!!!!!!!!!
	   ide(1,Ne)=ii;
	   ide(2,Ne)=jj;
	   mstv(ii)=.true.
	   mstv(jj)=.true.
	 enddo
	enddo
	do i=1,Ne
	 ii=ide(1,i);
	 jj=ide(2,i);
	 fvec=af(:,ii)-af(:,jj);!Vec jj->ii
	 dens_min=999999999.00;
	 a=0;
	 do j=1,9
	  ftmp=af(:,jj)+fvec*float(j)*0.1;
	  call MeanShift_Pos(ftmp,dens_data);
	  if(dens_data.lt.dens_min) dens_min=dens_data;
	 enddo
	 edge_dens(i)=dens_min*edge(i); !minimum density*length
	enddo

	!===================================
	edge_tmp=edge
	edge_best=edge
	call quick_sort(edge,order,Ne)
	write(*,*),"#Maxc=", Maxc," UseCid=",UseChain," Ne=",Ne
	write(*,*),"#sum_edgedens=", sum(edge_dens(1:Ne))
	write(*,*),"#ave_edgedens=", sum(edge_dens(1:Ne))/Ne
	!--------------------------------
	!Find MST using selected edge again!!
	 !init cid
	 do ii=1,n;cid(ii)=ii;enddo
	Nt=0
	activeg=.false.
	id2order(1:Ne)=-1
	 do ii=1,Ne
	  !minimum edge
	  Me=order(ii)
	  !if(Me.eq.rmid) cycle
	  v1=ide(1,Me)
	  v2=ide(2,Me)
	  if(cid(v1).ne.cid(v2)) then
	   !Add edge ii to Tree
	   Nt=Nt+1
	   Tree(Nt)=Me
	   activeg(Me)=.true. !using edge in initial MST
	   id2order(Me)=Nt
	   !Merge cluster
	   tmpid=cid(v2)
	   do jj=1,n
	    if(cid(jj).eq.tmpid) cid(jj)=cid(v1)
	   enddo
	  endif
	 enddo
	activeg_ori(1:Ne)=activeg(1:Ne)

	!Local Space MST
	local_edge(1:Ne)=.false.
	d2=dlocal*dlocal
	do pos=1,n
	 if(.not.mstv(pos)) cycle;

	 !init chain id
	 do ii=1,n;cid(ii)=ii;enddo

	 !write(*,*),"Local",pos
	 do ii=1,Ne
	  !minimum edge
	  Me=order(ii)
	  v1=ide(1,Me)
	  v2=ide(2,Me)
	  if(cid(v1).eq.cid(v2)) cycle
	  ftmp=af(1:3,pos)-af(1:3,v1);
	  if(dot_product(ftmp,ftmp).gt.d2) cycle;
	  ftmp=af(1:3,pos)-af(1:3,v2);
	  if(dot_product(ftmp,ftmp).gt.d2) cycle;
	  local_edge(Me)=.true.
	  !Merge cluster
	  tmpid=cid(v2)
	  do jj=1,n
	   if(cid(jj).eq.tmpid) cid(jj)=cid(v1)
	  enddo
	 enddo
	enddo
	!END Local Space MST
	Nel=0;
	do ii=1,Ne
	 !Add Initial MST edge
	 if(activeg_ori(ii)) local_edge(ii)=.true.
	 if(local_edge(ii)) Nel=Nel+1;
	enddo

	write(*,*),"#Ne=",Ne,"localNe=",Nel

	if(mode.eq.1)then
	 do ii=1,Nt
	  !write(*,'("bond resi ",I5," and tmp, resi "I5, " and tmp")'),
	  write(*,'("BOND ",2I6)'),
     *    ide(1:2,Tree(ii))
	 enddo
	 return
	else if(mode.eq.2) then
	 do ii=1,Ne
	  if(local_edge(ii)) then
	  !write(*,'("bond resi ",I5," and tmp, resi "I5, " and tmp")'),
	  write(*,'("BOND ",2I6)'),
     *    ide(1:2,ii)
	  endif
	 enddo
	 return
	endif

	ftotal=0
	do ii=1,Nt
	 ftotal=ftotal+edge_tmp(Tree(ii))
	enddo
	BestMST=ftotal
	call setup_tree(n,ide,Tree,Nt);
	call quality_tree(n,edge_tmp,edge_dens,
     *       ide,Tree,Nt,Nst,Ned,flen,fin,used_v,vpath,Nv)
	!stop
	PreNt=Nt;
	NtKth(1)=Nt
	NpKth(1)=Np
	TreeKth(1:Nt,1)=Tree(1:Nt)
	sco(1)=flen
	FscoMst=flen
	FtotalKth(1)=ftotal
	FtotalMst=ftotal;
	tmp_IDb=IDb
	write(*,*),"#Kth=",1,0,sco(1),Nt,Nst,Ned,flen,BestMST
	!----------------------------------
	if(.false.)then
	 do ii=1,NpKth(1)
	  write(*,'("bond resi ",I5," and tmp",I0,
     *    ", resi ",I5," and tmp",I0)'),
     *    ide(1,PathKth(ii,1)),1,ide(2,PathKth(ii,1)),1
	 enddo
	 stop
	endif

	!!SEARCH!!
	!----------------------------------
	allow_rate=allow_rate_max
	AllowMST=allow_rate*BestMST
	best_sco=best_len;
	TreeBest(1:Nt)=Tree(1:Nt);
	Best_Nv(1:10)=0;
	DO TRJ=1,10 !10 trajectories

	best_trj=0;
	Ntabu=0;
	TreeNow(1:Nt)=Tree(1:Nt);
	id2orderNow(1:Ne)=id2order(1:Ne)
	flag1=.false.
	activeg(1:Ne)=activeg_ori(1:Ne)
	FtotalNow=FtotalMst;
	!write(*,*),FtotalNow;
	!stop;
	!control stack
	!MaxTry=Nt*(Nel-Nt);
	MaxTry=Nel;
	MaxTry2=1;
	!write(*,*),"#MaxTry=",MaxTry,MaxTry2
	do ro=2,Nround
	 round_best_sco=0;
	 !Generate Local Candidates
	 TreeRoundBest(1:Nt)=TreeNow(1:Nt)

	 call setup_tree(n,ide,TreeNow(1:Nt),Nt);

	 Nbranch_tmp(1:n)=Nbranch(1:n)
	 do jj=1,n
          IDb_tmp(jj,1:Nbranch(jj))=IDb(jj,1:Nbranch(jj))
          id_edge_tmp(jj,1:Nbranch(jj))=id_edge(jj,1:Nbranch(jj))
	 enddo

	 allow_rate=allow_rate_max
	 !Two-edge swap
	 TryCnt2=0;
	 do ii=1,Nlocal


	  !call setup_tree(n,ide,TreeNow(1:Nt),Nt);
	  Nbranch(1:n)=Nbranch_tmp(1:n)
	  do jj=1,n
           IDb(jj,1:Nbranch(jj))=IDb_tmp(jj,1:Nbranch(jj))
           id_edge(jj,1:Nbranch(jj))=id_edge_tmp(jj,1:Nbranch(jj))
	  enddo
	  TryCnt=0;

 3001	  call random_number(a)

	  TryCnt=TryCnt+1;
	  if(TryCnt.gt.MaxTry) then
	   write(*,*),"#NumOfTry",TryCnt,MaxTry;
	   Ntabu=0;
	   TryCnt=0;
	   TryCnt2=TryCnt2+1;
	   !Avoid Too many loops
	   if(TryCnt2.gt.MaxTry2)then
	    TryCnt2=0;
	    goto 3003;
	    !cycle; ! end loop
	   endif
	  endif

	  OrderID=int(a*Nt)+1;
 	  p11=TreeNow(OrderID)
	  !New Kepp short path in the MST
	  if(activeg_ori(p11).and.edge_tmp(p11).lt.cut) goto 3001
          !Split chain
	  !write(*,*),"setup tree -1"
          call rmpath_tree(n,ide,p11);
          call split_chain(cid,n,ide,Nt,p11);

	!----set list-----
	  Nee=0
	  do jj=1,Ne
	   if(.not.local_edge(jj))then
	    cycle;
	   endif
	   !used in the tree
	   if(activeg(jj)) cycle;
	   v1=ide(1,jj); v2=ide(2,jj)
           if(cid(v1).eq.cid(v2)) cycle
           if(cid(v1)*cid(v2).eq.0) cycle
	   if(FtotalNow+edge_tmp(jj)-edge_tmp(p11).
     *     gt.AllowMST) cycle

	   Nee=Nee+1
	   EdgeTable(Nee)=jj;
	  enddo
	  if(Nee.eq.0)then
	   call addpath_tree(n,ide,p11);
	   goto 3001;
	  endif
	  activeg_tmp(1:Ne)=activeg(1:Ne)
          !Swap bad bonds
 	  flag=.false.

 	  do jj=1,Nee !try
 	   call random_number(a)
	   p12=EdgeTable(int(a*Nee)+1)
	   ftotal_tmp=FtotalNow+edge_tmp(p12)-edge_tmp(p11)
	   !if(ftotal_tmp.gt.AllowMST) cycle

	   !tabu check
	   do kk=1,Ntabu
	    if(TabuList(1,kk).eq.p11.and.TabuList(2,kk).eq.p12)
     *      goto 3002
	    if(TabuList(1,kk).eq.p12.and.TabuList(2,kk).eq.p11)
     *      goto 3002
	   enddo
	   tmp_tree(1:Nt)=TreeNow(1:Nt)
           tmp_tree(OrderID)=p12
	   activeg_tmp(p11)=.false.
	   activeg_tmp(p12)=.true.

	   flag=.true.
	   exit
 3002	  enddo
	  !write(*,*) "Nee1-2",Nee
	  if(.not.flag)then
	   call addpath_tree(n,ide,p11);
           goto 3001
	  endif
	  call addpath_tree(n,ide,p12);

	  !p21=p11;p22=p12;
	  ftotal=ftotal_tmp;
	  !write(*,*) "Nee1-3",Nee

	  call quality_tree(n,edge_tmp,edge_dens,ide,tmp_Tree,Nt,Nst,
     *          Ned,flen,fin,used_v,vpath,Nv)
	 !used_v=v of longest path
	  if(used_v(ide(1,p11)).and.used_v(ide(2,p11))
     *       .and.activeg_ori(p11)) then
	   call addpath_tree(n,ide,p11);
	   call rmpath_tree(n,ide,p12);
	   goto 3001;
	  endif

	  if(now_sco.eq.flen.and.ftotal.gt.FtotalNow)then
	   call addpath_tree(n,ide,p11);
	   call rmpath_tree(n,ide,p12);
	   goto 3001
	  endif
          if(round_best_sco.le.flen)then
	   if(ftotal.ge.ft_rb.and.round_best_sco.eq.flen)then
	    call addpath_tree(n,ide,p11);
	    call rmpath_tree(n,ide,p12);
	    goto 3001;
	   endif
	   !check Kendall tau
	   if(trj.gt.1)then
	    TmpPath(1:NowNp)=vpath(1:NowNp)-PathNow(1:NowNp)
	    if(maxval(TmpPath(1:NowNp)).ne.0.and.
     *      minval(TmpPath(1:NowNp)).ne.0) then
	    do ppp=1,trj-1
	     !if(best_trj_sco(ppp).ge.flen)then
	     fken=fkendall(vpath(1:Nv),Nv,
     *       best_path(ppp,:),best_Nv(ppp),n)
	     if(abs(fken).gt.0.95)then
	      !write(*,*),"Kendall=",fken,Nv,best_Nv(ppp)
	      call addpath_tree(n,ide,p11);
	      call rmpath_tree(n,ide,p12);
	      goto 3001;
	     endif
	    !endif
	    enddo
	   endif
	   endif

	   !END Kendall
           round_best_sco=flen
	   ft_rb=ftotal
           TreeRoundBest(1:Nt)=tmp_tree(1:Nt)
	   PathRoundBest(1:Nv)=vpath(1:Nv);
	   RoundBestNp=Nv
	   !id2orderRbest(1:Ne)=id2orderTmp(1:Ne)
	   ope(1)=p11; ope(2)=p12;
	   if(best_trj.lt.flen)then
	    best_trj=flen;
	    best_path(trj,1:Nv)=vpath(1:Nv);
	    best_Nv(trj)=Nv;
	    best_trj_sco(trj)=flen;
	    !write(*,*),"trjbest=",Nv;
	   endif
          endif
	  !FIN Local Search
    	 enddo !End Nlocal

	 write(*,*),trj,ro,round_best_sco,best_sco,ft_rb

	 !Update Now state
	 TreeNow(1:Nt)=TreeRoundBest(1:Nt)
	 PathNow(1:RoundBestNp)=PathRoundBest(1:RoundBestNp);
	 NowNp=RoundBestNp;
	 FtotalNow=ft_rb
	 FtotalNow=0;
	 do iii=1,Nt
	  FtotalNow=FtotalNow+edge_tmp(TreeNow(iii))
	 enddo
	 now_sco=round_best_sco

	!Update Best state
	 IF(best_sco.lt.round_best_sco)then
	  best_sco=round_best_sco
	  TreeBest(1:Nt)=TreeRoundBest(1:Nt)
	  TabuList(1,1)=ope(2)
	  TabuList(2,1)=ope(1)
	  Ntabu=1;
	 ELSE
	  Ntabu=Ntabu+1
	  if(Ntabu.gt.Ntb) Ntabu=Ntb
	  TabuListTmp(1:2,1:Ntabu)=TabuList(1:2,1:Ntabu)
	  TabuList(1,1)=ope(2)
	  TabuList(2,1)=ope(1)
	  TabuList(1:2,2:Ntabu)=TabuListTmp(1:2,1:Ntabu-1);
	 ENDIF

	 activeg(1:Ne)=.false.

	 do ii=1,Nt
	  activeg(TreeNow(ii))=.true.
	 enddo
   	enddo
	ENDDO
	!=====================================================
	 call longestpath_tree(n,edge_tmp,ide,TreeBest,Nt,Nst,
     *          Ned,flen,path,Np)
 	Nlgst=0
	  do ii=1,Np-1
	   v1=ide(1,Path(ii));
	   v2=ide(2,Path(ii));
	   v21=ide(1,Path(ii+1));
	   v22=ide(2,Path(ii+1));
	   if(v1.eq.v21.or.v1.eq.v22)then
	    Nlgst=Nlgst+1
	    Plgst(Nlgst)=v2;
	    Plgst(Nlgst+1)=v1;
	   else
	    Nlgst=Nlgst+1
	    Plgst(Nlgst)=v1;
	    Plgst(Nlgst+1)=v2;
	   endif
	  enddo
	  ii=Np;
	  v1=ide(1,Path(ii));
	  v2=ide(2,Path(ii));
	  if(Plgst(Nlgst+1).eq.v1)then
	   Nlgst=Nlgst+1
	   Plgst(Nlgst)=v1;
	   Nlgst=Nlgst+1
	   Plgst(Nlgst)=v2;
	  else
	   Nlgst=Nlgst+1
	   Plgst(Nlgst)=v2;
	   Nlgst=Nlgst+1
	   Plgst(Nlgst)=v1;
	  endif
 3003	END


	SUBROUTINE CHECK_TREE(n,edge,ide,Tree,Nt,
     *                 list_bad,Nbad,bad_score)
	IMPLICIT INTEGER(I-Z)
	PARAMETER(MSIZE=300)
	PARAMETER(MAXB=5)
	DIMENSION af(3,MSIZE*MSIZE*10),fin(MSIZE*MSIZE*MSIZE)
	COMMON /GRAPH/ Nbranch(MSIZE*MSIZE*10),IDb(MSIZE*MSIZE*10,MAXB*2),
     *	edged(MSIZE*MSIZE*10,MAXB*2),
     *		nextp(MSIZE*MSIZE*10),
     *		cost(MSIZE*MSIZE*10),
     *  id_edge(MSIZE*MSIZE*10,MAXB*2)
	DIMENSION ftmp(3),cd_st(3),cd_ed(3)
	LOGICAL discovered(MSIZE*MSIZE*10)
	DIMENSION list_act(MSIZE*MSIZE) !active list
	DIMENSION tmp_list(MSIZE*MSIZE)
	DIMENSION path(MSIZE*MSIZE)
	LOGICAL flag,used(MSIZE*MSIZE*10)
	INTEGER cid(n)
	DIMENSION edge(n*n),ide(2,n*n),order(n*n),list_bad(n*n),
     *  Tree(n*n),edge_tmp(n*n),TreeKth(100,n*n),NtKth(100)
	real sco(100),path_branch(n,MAXB*2),edge_sum(n*n)

	!MAke tree
	!init
	Nbranch=0
	IDb=0
	do ii=1,Nt
	 it=Tree(ii)
	 id1=ide(1,it)
	 id2=ide(2,it)
	 !input
	 Nbranch(id1)=Nbranch(id1)+1
	 IDb(id1,Nbranch(id1))=id2
	 edged(id1,Nbranch(id1))=edge(it)
	 path_branch(id1,Nbranch(id1))=it

	 Nbranch(id2)=Nbranch(id2)+1
	 IDb(id2,Nbranch(id2))=id1
	 edged(id2,Nbranch(id2))=edge(it)
	 path_branch(id2,Nbranch(id2))=it
	enddo


	Nbad=0
	!check balance of branch
	used=.false.
	ftotal=0
	do ii=1,Nt
	 ftotal=ftotal+1.00
	 do jj=1,2
	  !vv=ide(jj,path(ii))
	  vv=ide(jj,Tree(ii))
	  if(used(vv)) cycle
	  if(Nbranch(vv).lt.3) cycle
	  !write(*,*),"vv=",vv,Nbranch(vv)
	  used(vv)=.true.

	  !check the balance of branches
	  edge_sum=0
	  Nb=Nbranch(vv)
	  do kk=1,Nbranch(vv)
	   root=IDb(vv,kk)
	   !DFS
	   Na=1
	   list_act(Na)=root
	   discovered=.false.
	   discovered(vv)=.true.
	   edge_sum(kk)=edged(vv,kk)
	   do p=1,Nt
	    v=list_act(1) !pop
	    list_act(1:Na-1)=list_act(2:Na)
	    Na=Na-1
	    if(.not.discovered(v))then
	     discovered(v)=.true.
	     do iii=1,Nbranch(v)
	      w=IDb(v,iii)
	      if(discovered(w)) cycle
	      tmp_list(1:Na)=list_act(1:Na)
	      list_act(1)=w
	      list_act(2:Na+1)=tmp_list(1:Na)
	      Na=Na+1
	      edge_sum(kk)=edge_sum(kk)+sqrt(edged(v,iii))
	     enddo
	    endif
	    if(Na.eq.0) exit
	   enddo
	   !write(*,*),kk,"EdgeSum=",edge_sum(kk),root
	  enddo
	  call quick_sort(edge_sum,order,Nbranch(vv));
	  frate=(edge_sum(Nb)+edge_sum(Nb-1))/sum(edge_sum(1:Nb))
	  if(frate.lt.0.99)then
	   write(*,*),"Warrning!! ",vv,Nbranch(vv),frate
	   Nbad=Nbad+1
	   list_bad(Nbad)=vv
	  endif
	  ftotal=ftotal+sum(edge_sum(1:Nb-3))
	 enddo
	enddo
	write(*,*),"lost edge =",ftotal
	bad_score=ftotal
	END

	!Setup Nbranch and id_edge
	SUBROUTINE SETUP_TREE(n,ide,Tree,Nt)
	IMPLICIT INTEGER(I-Z)
	PARAMETER(MSIZE=300)
	PARAMETER(MAXB=5)
	!DIMENSION af(3,MSIZE*MSIZE*10),fin(MSIZE*MSIZE*MSIZE)
	COMMON /GRAPH/ Nbranch(MSIZE*MSIZE*10),IDb(MSIZE*MSIZE*10,MAXB*2),
     *	edged(MSIZE*MSIZE*10,MAXB*2),
     *	nextp(MSIZE*MSIZE*10),
     *	cost(MSIZE*MSIZE*10),
     *  id_edge(MSIZE*MSIZE*10,MAXB*2)
	DIMENSION ide(2,n*n),
     *  Tree(n*n)

	!init
	Nbranch=0
	IDb=0

	do ii=1,Nt
	 it=Tree(ii)
	 id1=ide(1,it)
	 id2=ide(2,it)
	 !input
	 Nbranch(id1)=Nbranch(id1)+1
	 IDb(id1,Nbranch(id1))=id2
	 id_edge(id1,Nbranch(id1))=it;
	 Nbranch(id2)=Nbranch(id2)+1
	 IDb(id2,Nbranch(id2))=id1;
	 id_edge(id2,Nbranch(id2))=it;
	enddo

	END

	SUBROUTINE ADDPATH_TREE(n,ide,iadd)
	IMPLICIT INTEGER(I-Z)
	PARAMETER(MSIZE=300)
	PARAMETER(MAXB=5)
	COMMON /GRAPH/ Nbranch(MSIZE*MSIZE*10),IDb(MSIZE*MSIZE*10,MAXB*2),
     *	edged(MSIZE*MSIZE*10,MAXB*2),
     *		nextp(MSIZE*MSIZE*10),
     *		cost(MSIZE*MSIZE*10),
     *  id_edge(MSIZE*MSIZE*10,MAXB*2)
	DIMENSION ide(2,n*n)

	id1=ide(1,iadd)
	id2=ide(2,iadd)

	!input
	Nbranch(id1)=Nbranch(id1)+1
	IDb(id1,Nbranch(id1))=id2
	id_edge(id1,Nbranch(id1))=iadd
	Nbranch(id2)=Nbranch(id2)+1
	IDb(id2,Nbranch(id2))=id1
	id_edge(id2,Nbranch(id2))=iadd

	END

	SUBROUTINE RMPATH_TREE(n,ide,irm)
	IMPLICIT INTEGER(I-Z)
	PARAMETER(MSIZE=300)
	PARAMETER(MAXB=5)
	COMMON /GRAPH/ Nbranch(MSIZE*MSIZE*10),IDb(MSIZE*MSIZE*10,MAXB*2),
     *	edged(MSIZE*MSIZE*10,MAXB*2),
     *		nextp(MSIZE*MSIZE*10),
     *		cost(MSIZE*MSIZE*10),
     *  id_edge(MSIZE*MSIZE*10,MAXB*2)
	DIMENSION ide(2,n*n)

	id1=ide(1,irm)
	id2=ide(2,irm)

	!remove
	do i=1,Nbranch(id1)
	 if(IDb(id1,i).eq.id2)then
	  !shift
	  IDb(id1,i:Nbranch(id1)-1)=IDb(id1,i+1:Nbranch(id1))
	  id_edge(id1,i:Nbranch(id1)-1)=id_edge(id1,i+1:Nbranch(id1))
	  Nbranch(id1)=Nbranch(id1)-1
	  exit;
	 endif
	enddo
	do i=1,Nbranch(id2)
	 if(IDb(id2,i).eq.id1)then
	  !shift
	  IDb(id2,i:Nbranch(id2)-1)=IDb(id2,i+1:Nbranch(id2))
	  id_edge(id2,i:Nbranch(id2)-1)=id_edge(id2,i+1:Nbranch(id2))
	  Nbranch(id2)=Nbranch(id2)-1
	  exit;
	 endif
	enddo

	END


	SUBROUTINE SPLIT_CHAIN(cid,n,ide,Nt,igp)
	IMPLICIT INTEGER(I-Z)
	PARAMETER(MSIZE=300)
	PARAMETER(MAXB=5)
	COMMON /GRAPH/ Nbranch(MSIZE*MSIZE*10),IDb(MSIZE*MSIZE*10,MAXB*2),
     *	edged(MSIZE*MSIZE*10,MAXB*2),
     *		nextp(MSIZE*MSIZE*10),
     *		cost(MSIZE*MSIZE*10),
     *  id_edge(MSIZE*MSIZE*10,MAXB*2)
	LOGICAL discovered(MSIZE*MSIZE*10)
	DIMENSION list_act(MSIZE*MSIZE) !active list
	LOGICAL flag
	INTEGER cid(n)
	DIMENSION ide(2,n*n)

	!Depth First Search From id1->???
	!chain1 and 2
	cid(1:n)=0
	do idchain=1,2
	root=ide(idchain,igp)
	cid(root)=idchain;
	!push
	Na=1
	list_act(Na)=root
	discovered(1:n)=.false.
	do p=1,Nt
	 v=list_act(Na) !pop
	 Na=Na-1
	 if(.not.discovered(v))then
	  discovered(v)=.true.
	  do ii=1,Nbranch(v)
	   w=IDb(v,ii)
	   if(discovered(w)) cycle
	   Na=Na+1
	   list_act(Na)=w
	   cid(w)=idchain;
	  enddo
	 endif
	 if(Na.eq.0) exit
	enddo
	enddo

	END

	SUBROUTINE LONGESTPATH_TREE(n,edge,ide,Tree,Nt,Nst,Ned,flen,
     *                              path,Np)
	IMPLICIT INTEGER(I-Z)
	PARAMETER(MSIZE=300)
	PARAMETER(MAXB=5)
	DIMENSION af(3,MSIZE*MSIZE*10),fin(MSIZE*MSIZE*MSIZE)
	COMMON /GRAPH/ Nbranch(MSIZE*MSIZE*10),IDb(MSIZE*MSIZE*10,MAXB*2),
     *	edged(MSIZE*MSIZE*10,MAXB*2),
     *		nextp(MSIZE*MSIZE*10),
     *		cost(MSIZE*MSIZE*10),
     *  id_edge(MSIZE*MSIZE*10,MAXB*2)
	DIMENSION ftmp(3),cd_st(3),cd_ed(3)
	LOGICAL discovered(MSIZE*MSIZE*10)
	DIMENSION list_act(MSIZE*MSIZE) !active list
	DIMENSION tmp_list(MSIZE*MSIZE)
	DIMENSION path(MSIZE*MSIZE)
	LOGICAL flag
	INTEGER cid(n)
	DIMENSION edge(n*n),ide(2,n*n),order(n*n),
     *  Tree(n*n),edge_tmp(n*n),TreeKth(100,n*n),NtKth(100)
	real sco(100),path_branch(n,MAXB*2)

	!init
	Nbranch(1:n)=0
	IDb=0

	do ii=1,Nt
	 it=Tree(ii)
	 id1=ide(1,it)
	 id2=ide(2,it)
	 !input
	 Nbranch(id1)=Nbranch(id1)+1
	 IDb(id1,Nbranch(id1))=id2
	 edged(id1,Nbranch(id1))=edge(it)
	 path_branch(id1,Nbranch(id1))=it

	 Nbranch(id2)=Nbranch(id2)+1
	 IDb(id2,Nbranch(id2))=id1
	 edged(id2,Nbranch(id2))=edge(it)
	 path_branch(id2,Nbranch(id2))=it

	enddo

	!Depth First Search From id1->???
	root=id1
	pre_root=-1
	do icnt=1,100
	 cost(1:n)=0
	 !push
	 Na=1
	 list_act(Na)=root
	 discovered=.false.
	 fmaxc=0
	 maxi=-1
	 do p=1,Nt
	  v=list_act(Na) !pop
	  Na=Na-1
	  if(.not.discovered(v))then
	   discovered(v)=.true.
	   do ii=1,Nbranch(v)
	    w=IDb(v,ii)
	    if(discovered(w)) cycle
	    Na=Na+1
	    list_act(Na)=w;
	    cost(w)=cost(v)+edged(v,ii)
	    nextp(w)=path_branch(v,ii)
	    if(cost(w).ge.fmaxc)then
	     fmaxc=cost(w)
	     maxi=w
	    endif
	   enddo
	  endif
	  if(Na.eq.0) exit
	 enddo

	 if(maxi.eq.pre_root) exit
	 pre_root=root
	 root=maxi
	enddo
	Nst=root;Ned=maxi;flen=fmaxc;
	!trace back
	now=maxi
	Np=0
	do ii=1,Nt
	 Np=Np+1
	 path(Np)=nextp(now)
	 next_v=ide(1,nextp(now))
	 if(next_v.eq.now) next_v=ide(2,nextp(now))
	 if(next_v.eq.root) exit
	 now=next_v
	enddo

	END


	SUBROUTINE QUALITY_TREE(n,edge,edge_dens,ide,Tree,Nt,Nst,Ned,flen,
     *                              fin,used_v,vpath,Nv)
	IMPLICIT INTEGER(I-Z)
	PARAMETER(MSIZE=300)
	PARAMETER(MAXB=5)
	DIMENSION af(3,MSIZE*MSIZE*10),fin(MSIZE*MSIZE*MSIZE)
	COMMON /GRAPH/ Nbranch(MSIZE*MSIZE*10),IDb(MSIZE*MSIZE*10,MAXB*2),
     *	edged(MSIZE*MSIZE*10,MAXB*2),
     *	nextp(MSIZE*MSIZE*10),
     *	cost(MSIZE*MSIZE*10),
     *  id_edge(MSIZE*MSIZE*10,MAXB*2)
	LOGICAL discovered(MSIZE*MSIZE*10),used(n),
     *	used_v(n)
	DIMENSION list_act(MSIZE*MSIZE) !active list
	!DIMENSION tmp_list(MSIZE*MSIZE)
	LOGICAL flag
	!INTEGER cid(n)
	DIMENSION edge(n*n),ide(2,n*n),edge_dens(n*n),
     *  Tree(n*n),edge_tmp(n*n)
	DIMENSION vpath(n),nextv(n);
	INTEGER best_v;
	COMMON /KRU/ Nround,Nlocal,Ntabu,allow_rate_max,dlocal,dside,mode

	!Depth First Search From id1->???
	!root=id1
	root=ide(1,Tree(Nt))
	pre_root=-1
	do icnt=1,100
	 cost(1:n)=0
	 !push
	 Na=1
	 list_act(Na)=root
	 discovered(1:n)=.false.
	 fmaxc=0
	 maxi=-1
	 do p=1,Nt
	  v=list_act(Na) !pop
	  Na=Na-1
	  if(.not.discovered(v))then
	   discovered(v)=.true.
	   do ii=1,Nbranch(v)
	    w=IDb(v,ii)
	    if(discovered(w)) cycle
	    Na=Na+1
	    list_act(Na)=w;
	    cost(w)=cost(v)+edge_dens(id_edge(v,ii))
	    nextv(w)=v;
	    if(cost(w).ge.fmaxc)then
	     fmaxc=cost(w)
	     maxi=w
	    endif
	   enddo
	  endif
	  if(Na.eq.0) exit
	 enddo
	 if(maxi.eq.pre_root) exit
	 pre_root=root
	 root=maxi
	enddo
	!write(*,*),"maxci=",fmaxc,maxi," root=",root
	Nst=root;Ned=maxi;flen=fmaxc;
	flen=fmaxc*fmaxc;!1st longest path
	!write(*,*),"#longest:",root,fmaxc
	!trace back
	now=maxi
	Np=0
	Nv=1;vpath(Nv)=now;
	used(1:n)=.false.

	!1st path
	do ii=1,Nt
	 !Np=Np+1
	 next_v=nextv(now)
	 Nv=Nv+1; vpath(Nv)=next_v;
	 used(next_v)=.true.
	 if(next_v.eq.root) exit
	 now=next_v
	enddo
	used_v(1:n)=used(1:n);
	!used_v(maxi)=.false. !Ignore terminal <-New
	used_v(root)=.false. !Ignore terminal

	!if(.false.) then

	!write(*,*),vpath(1:Nv);
	!Find long branches 2nd 3rd,....100th,
	do ro=2,100
	 best_len=0;
	 best_v=-1;
	 do ii=1,n
	  if(used(ii)) cycle;
	  if(Nbranch(ii).ne.1) cycle;
	  !traca back
	  now=ii;
	  do jj=1,n
	   next_v=nextv(now)
	   if(used(next_v)) exit;
	   now=next_v
	  enddo
	 !write(*,*),"**",ii,next_v,cost(ii)-cost(next_v);
	  diff_cost=cost(ii)-cost(next_v);
	  if(best_len.lt.diff_cost)then
	   best_len=diff_cost;
	   best_v=ii;
	  endif
	 enddo
	 if(best_v.eq.-1) exit;
	 flen=flen+best_len*best_len;
	 !trace back
	  now=best_v;
	  used(now)=.true.
	 do jj=1,n
	  next_v=nextv(now)
	  if(used(next_v)) exit;
	  used(next_v)=.true.
	  now=next_v
	 enddo
	enddo
	!endif

	END

	SUBROUTINE SETUP_GRAPH(af,n,fin,dl,du)
	IMPLICIT INTEGER(I-Z)
	PARAMETER(MSIZE=300)
	PARAMETER(MAXB=5)
	DIMENSION af(3,MSIZE*MSIZE*10),fin(MSIZE*MSIZE*MSIZE)
	COMMON /GRAPH/ Nbranch(MSIZE*MSIZE*10),IDb(MSIZE*MSIZE*10,MAXB*2),
     *	edged(MSIZE*MSIZE*10,MAXB*2),nextp(MSIZE*MSIZE*10),
     *  cost(MSIZE*MSIZE*10),
     *  id_edge(MSIZE*MSIZE*10,MAXB*2)
	DIMENSION ftmp(3),fmid(3)
	logical flag
	d2=d*d


	fmax=maxval(fin(1:n))
	write(*,*),fmax
	!init
	Nbranch(1:n)=0;
	!write(*,*) ,"d1=",d1,"d2=",d2
	do ii=1,n-1
	 do jj=ii+1,n
	  ftmp=af(1:3,ii)-af(1:3,jj)

	  !check=dot_product(ftmp,ftmp)
	  check=sqrt(dot_product(ftmp,ftmp))
	  !write(*,*),ii,jj,check
	  if(check.lt.du.and.check.gt.dl)then
	   fmid=(af(1:3,ii)+af(1:3,jj))*0.5
	   check=check
	   if(Nbranch(ii).lt.MAXB)then
	    Nbranch(ii)=Nbranch(ii)+1
	    IDb(ii,Nbranch(ii))=jj
	    edged(ii,Nbranch(ii))=check
	   else
	    dmax=0;mid=0
	    do kk=1,MAXB
	     if(dmax.lt.edged(ii,kk))then
	      dmax=edged(ii,kk)
	      mid=kk
	     endif
	    enddo
	    if(dmax.gt.check)then
	     IDb(ii,mid)=jj
	     edged(ii,mid)=check
	    endif
	   endif
	   if(Nbranch(jj).lt.MAXB)then
	    Nbranch(jj)=Nbranch(jj)+1
	    IDb(jj,Nbranch(jj))=ii
	    edged(jj,Nbranch(jj))=check
	   else
	    dmax=0;mid=0
	    do kk=1,MAXB
	     if(dmax.lt.edged(jj,kk))then
	      dmax=edged(jj,kk)
	      mid=kk
	     endif
	    enddo
	    if(dmax.gt.check)then
	     IDb(jj,mid)=ii
	     edged(jj,mid)=check
	    endif
	   endif
	  endif
	 enddo
	 !write(*,*) ii,Nbranch(ii),IDb(ii,1:Nbranch(ii))
	 !write(*,'(10f6.2)') edged(ii,1:Nbranch(ii))
	enddo
	!do ii=1,n
	! write(*,*) ii,Nbranch(ii),IDb(ii,1:Nbranch(ii))
	! write(*,'(10f6.2)') edged(ii,1:Nbranch(ii))
	!enddo
	!Merge
	do ii=1,n
	 do jj=1,Nbranch(ii)
	  kk=IDb(ii,jj)
	  flag=.false.
	  do ll=1,Nbranch(kk)
	   if(IDb(kk,ll).eq.ii) then
	    flag=.true.
	    exit
	   endif
	  enddo
	 if(.not.flag)then
	  if(Nbranch(kk).lt.MAXB*2)then
	   Nbranch(kk)=Nbranch(kk)+1
	   IDb(kk,Nbranch(kk))=ii
	   edged(kk,Nbranch(kk))=edged(ii,jj)
	  !else

	  endif
	 endif
	 enddo
	enddo
	END

	SUBROUTINE MERGEPOINTS(af,n,fin,d,cut,member)
	IMPLICIT INTEGER(I-Z)
	PARAMETER(MSIZE=300)
	DIMENSION af(3,MSIZE*MSIZE*10),fin(MSIZE*MSIZE*MSIZE)
	LOGICAL stock(MSIZE*MSIZE*10)
	DIMENSION ftmp(3),member(MSIZE*MSIZE*10),
     *  tmp_member(MSIZE*MSIZE*10)
	write(*,*),"#N=",n
	stock(1:n)=.true.;
	Ncnt=1;
	d2=d*d

	fmax=maxval(fin(1:n))
	fmin=minval(fin(1:n))
	frange=fmax-fmin

	!init member data
	do ii=1,n
	 member(ii)=ii;
	enddo

	do ii=1,n-1
	 if((fin(ii)-fmin)/frange.lt.cut) stock(ii)=.false.
   	 if(.not.stock(ii)) cycle
	 do jj=ii+1,n
	  if(.not.stock(jj)) cycle
	  if((fin(jj)-fmin)/frange.lt.cut) stock(jj)=.false.
	  if(.not.stock(jj)) cycle
	  !check dist
	  ftmp=af(1:3,ii)-af(1:3,jj)
	  check=dot_product(ftmp,ftmp)
	  !write(*,*),ii,jj,check,ftmp,af(1:3,ii),af(1:3,jj)
	  if(check.lt.d2)then
	   if(fin(ii).gt.fin(jj))then
	    !keep ii
	    stock(jj)=.false.
	    member(jj)=ii
	   else !keep jj
	    stock(ii)=.false.
	    member(ii)=jj
	    exit
	   endif
	  endif
	 enddo
	enddo

	do ii=1,n
	 now=member(ii)
	 do jj=1,n
	  if(now.eq.member(now)) exit;
	  now=member(now)
	 enddo
	 member(ii)=now
	enddo

	!Copy
	Ncnt=1;
	do ii=1,n
	 if(stock(ii))then
	  af(:,Ncnt)=af(:,ii)
	  fin(Ncnt)=fin(ii)
	  tmp_member(ii)=Ncnt
	  Ncnt=Ncnt+1
	 endif
	enddo
	do ii=1,n
	 member(ii)=tmp_member(member(ii))
	 !write(*,*),"mb=",ii,member(ii)
	enddo
	n=Ncnt-1
	write(*,*),"#After merge N=",n
	END

	FUNCTION GFILTER_VALUE(cd)
	IMPLICIT INTEGER(I-Z)
        PARAMETER(NDIM=800)
        PARAMETER(MSIZE=300)
        DIMENSION p_xyz(3),tmp(3)
	DIMENSION af(3),bf(3),ftmp(3),cd(3)
	INTEGER stp(3),edp(3)
	COMMON /MAP_DATA/  gstep,gbase(3),NumOfGrid(3),dreso,
     *          dmap(MSIZE,MSIZE,MSIZE),amap(MSIZE,MSIZE,MSIZE),
     *          cmap(MSIZE,MSIZE,MSIZE), !raw data
     *          obox(3),pbox(3),list(NDIM),Nlist,lbase(3),lbase1(3)

	fs=(dreso/gstep)*0.5
	fs=fs*fs
	fmaxd=(dreso/gstep)*2.0

	ftmp=cd
	stp=nint(ftmp-fmaxd)
	edp=nint(ftmp+fmaxd)
	do ii=1,3
	 if(stp(ii).lt.1) stp(ii)=1
	 if(edp(ii).gt.NumOfGrid(ii)) edp(ii)=NumOfGrid(ii)
	enddo

	ftotal=0
	do kk=stp(3),edp(3)
	 fr3=ftmp(3)-float(kk); fr3=fr3*fr3
	do jj=stp(2),edp(2)
	 fr2=ftmp(2)-float(jj); fr2=fr2*fr2
	do ii=stp(1),edp(1)
	 fr1=ftmp(1)-float(ii); fr1=fr1*fr1
	 fr=fr1+fr2+fr3
	 d=exp(-1.50*fr/fs)*amap(ii,jj,kk)
	 ftotal=ftotal+d
	enddo
	enddo
	enddo
	gfilter_value=ftotal

	END

	SUBROUTINE MEANSHIFT(bf,af,fout,as)
	IMPLICIT INTEGER(I-Z)
        PARAMETER(NDIM=800)
        PARAMETER(MSIZE=300)
        DIMENSION p_xyz(3),tmp(3)
	DIMENSION af(3),bf(3),ftmp(3)
	INTEGER stp(3),edp(3)
	COMMON /MAP_DATA/  gstep,gbase(3),NumOfGrid(3),dreso,
     *          dmap(MSIZE,MSIZE,MSIZE),amap(MSIZE,MSIZE,MSIZE),
     *          cmap(MSIZE,MSIZE,MSIZE), !raw data
     *          obox(3),pbox(3),list(NDIM),Nlist,lbase(3),lbase1(3)
	stack=0
	!Gaussian kernel dreso=window size
	fs=(dreso/gstep)*0.5
	fs=fs*fs
	fsiv=1.000/fs
	fmaxd=(dreso/gstep)*2.0
	!write(*,'("bf=",4f8.2)'),bf,fs
	d2=as*as;

	ftmp=bf
	do
	if(stack.gt.10000) exit
	stp=nint(ftmp-fmaxd)
	edp=nint(ftmp+fmaxd)
	do ii=1,3
	 if(stp(ii).lt.1) stp(ii)=1
	 if(edp(ii).gt.NumOfGrid(ii)) edp(ii)=NumOfGrid(ii)
	enddo

	!write(*,'("bf=",3I4,3I4)'),stp,edp
	ftotal=0
	af=0
	do kk=stp(3),edp(3)
	 fr3=ftmp(3)-float(kk); fr3=fr3*fr3
	do jj=stp(2),edp(2)
	 fr2=ftmp(2)-float(jj); fr2=fr2*fr2
	do ii=stp(1),edp(1)
	 fr1=ftmp(1)-float(ii); fr1=fr1*fr1
	 fr=fr1+fr2+fr3
	 d=exp(-1.50*fr*fsiv)*amap(ii,jj,kk)
	 af=af+d*(/ii,jj,kk/)
	 ftotal=ftotal+d
	enddo
	enddo
	enddo
	if(ftotal.eq.0.000) exit
	 af=af/ftotal
	ftmp=ftmp-af
	if(dot_product(ftmp,ftmp).lt.0.001) exit
	ftmp=af-bf;
	if(dot_product(ftmp,ftmp).gt.d2) exit

c	write(*,'("#af=",3f8.3,"total=",f10.4,f10.4)'),af,ftotal,
c     *  dot_product(ftmp,ftmp)
	ftmp=af
	stack=stack+1
	enddo
	fout=ftotal
	!write(*,'("##af=",4f8.3)'),af,ftotal
	END

	SUBROUTINE MEANSHIFT_POS(bf,fout)
	IMPLICIT INTEGER(I-Z)
        PARAMETER(NDIM=800)
        PARAMETER(MSIZE=300)
        DIMENSION p_xyz(3),tmp(3)
	DIMENSION af(3),bf(3),ftmp(3)
	INTEGER stp(3),edp(3)
	COMMON /MAP_DATA/  gstep,gbase(3),NumOfGrid(3),dreso,
     *          dmap(MSIZE,MSIZE,MSIZE),amap(MSIZE,MSIZE,MSIZE),
     *          cmap(MSIZE,MSIZE,MSIZE), !raw data
     *          obox(3),pbox(3),list(NDIM),Nlist,lbase(3),lbase1(3)
	stack=0
	!Gaussian kernel dreso=window size
	fs=(dreso/gstep)*0.5
	fs=fs*fs
	fsiv=1.000/fs
	fmaxd=(dreso/gstep)*2.0

	ftmp=bf

	stp=nint(ftmp-fmaxd)
	edp=nint(ftmp+fmaxd)
	do ii=1,3
	 if(stp(ii).lt.1) stp(ii)=1
	 if(edp(ii).gt.NumOfGrid(ii)) edp(ii)=NumOfGrid(ii)
	enddo

	ftotal=0
	af=0
	do kk=stp(3),edp(3)
	 fr3=ftmp(3)-float(kk); fr3=fr3*fr3
	do jj=stp(2),edp(2)
	 fr2=ftmp(2)-float(jj); fr2=fr2*fr2
	do ii=stp(1),edp(1)
	 fr1=ftmp(1)-float(ii); fr1=fr1*fr1
	 fr=fr1+fr2+fr3
	 d=exp(-1.50*fr*fsiv)*amap(ii,jj,kk)
	 ftotal=ftotal+d
	enddo
	enddo
	enddo

	fout=ftotal
	!write(*,'("##af=",4f8.3)'),af,ftotal
	END


	SUBROUTINE POWER(a,n,x,xmax,itr)
	implicit real*8 (a-h,o-z)
	dimension a(n,n),x(n),y(n)
	m=0
	xmax=0
	c=dot_product(x(1:n),x(1:n))
	x=x/dsqrt(c)
	do while (m.lt.iter)
	 m=m+1
	 y=matmul(a,x)
	 s=dot_product(y(1:n),y(1:n))
	 c= s/dot_procuct(y(1:n),x(1:n))
	 if(dabs(xmax-c).lt.1.d-6) exit
	 xmax=c
	 x=y/dsqrt(s)
	enddo
	iter=m
	return

	END


	FUNCTION ROULETTE(fin,Np)
        PARAMETER(NDIM=800)
        IMPLICIT INTEGER(I-Z)
        PARAMETER(MSIZE=300)
	DIMENSION fin(MSIZE*MSIZE*MSIZE)
        !Find 3 to lenf2
        ftotal=0
        do ii=1,Np
         ftotal=ftotal+fin(ii) !select bad regions
        enddo

        call random_number(a)

	a=a*ftotal

	ftotal=0
        do ii=1,Np
         ftotal=ftotal+fin(ii)
         if (ftotal.gt.a) then
          ROULETTE=ii
          exit
         endif
        enddo
        END

	SUBROUTINE SET_CCENTER(posi,fin,SpaseN,Np,fmind,fmaxd)

        IMPLICIT INTEGER(I-Z)
        PARAMETER(NDIM=800)
        PARAMETER(MSIZE=300)
        DIMENSION p_xyz(3),tmp(3)
	DIMENSION posi(3,msize*msize*msize),fin(msize*msize*msize)
	DIMENSION ftable(msize*msize*msize)
	COMMON /CLUSTER/ Ccenter(3,ndim),Nc(ndim),
     *         p2c(MSIZE*MSIZE*MSIZE),
     *         ic2p(ndim,MSIZE*MSIZE),fsum(ndim),ainfo(ndim)

	r2=fmind*fmind

	!ii=roulette(fin,SpaseN)
	!Ccenter(:,1)=posi(:,ii)
	!write(*,*) "r2=",r2,ii
	icnt=0;
	do i=1,Np
	 icnt=0
	 if(Nc(i).gt.0) cycle
 9991	 ii=roulette(fin,SpaseN)
	 icnt=icnt+1
	 if(icnt.gt.SpaseN*2)then
	  write(*,*) "Cannot find intial position..."
	  stop
	 endif
	!check mindist
	 do k=1,Np
	  if(Nc(k).eq.0.or.k.eq.i) cycle
	  tmp(:)=posi(:,ii)-Ccenter(:,k)
	  ir=dot_product(tmp,tmp)
	  if(ir.lt.r2)then
	   goto 9991
	  endif
	 enddo
	 Ccenter(:,i)=posi(:,ii)
	 Nc(i)=1
	 !write(*,*) i,ii
	enddo

	!Assign initial cluster member
	Nc=0
	do i=1,SpaseN
	 mind=0
	 do j=1,Np
	  tmp(:)=posi(:,i)-Ccenter(:,j)
	  ir=dot_product(tmp,tmp)
	  if(j.eq.1.or.ir.lt.mind)then
	   !write(*,*) i,"->",j,ir
	   mind=ir
	   id=j
	   !write(*,*) i,"->",id
	  endif
	 enddo
	 Nc(id)=Nc(id)+1
	 ic2p(id,Nc(id))=i
	 p2c(i)=id
	 !write(*,*) i,"->",id,Nc(id)
	enddo

	END

        SUBROUTINE CONV_AMAP_DOUBLE()

        IMPLICIT INTEGER(I-Z)
        PARAMETER(NDIM=800)
        PARAMETER(MSIZE=300)
        DIMENSION p_xyz(3),tmp(3)
        COMMON /MAP_DATA/  gstep,gbase(3),NumOfGrid(3),dreso,
     *          dmap(MSIZE,MSIZE,MSIZE),amap(MSIZE,MSIZE,MSIZE),
     *          cmap(MSIZE,MSIZE,MSIZE), !raw data
     *          obox(3),pbox(3),list(NDIM),Nlist,lbase(3),lbase1(3)
        DIMENSION ftmp(3),atmp(3),btmp(3),min_vox(3),max_vox(3)
        DIMENSION minp(3),maxp(3)

        !###############
        step=1 !=0.61*2

        maxp=NumOfGrid*2

        write(*,"('Maxp=' I4, I4, I4)"),maxp

	dmap=0;

        do ii=1,maxp(1)

	 atmp(1)=(ii-1)*0.5*gstep;
         tmp(1)=int(atmp(1)/gstep)+1

	 if(tmp(1).lt.1) cycle
	 fx=(atmp(1)/gstep+1-tmp(1))
         do jj=1,maxp(2)
	  atmp(2)=(jj-1)*0.5*gstep
          tmp(2)=int((atmp(2))/gstep)+1
	  fy=(atmp(2)/gstep+1-tmp(2))
	  if(tmp(2).lt.1) cycle
          do kk=1,maxp(3)
	   atmp(3)=(kk-1)*0.5*gstep
           tmp(3)=int(atmp(3)/gstep)+1
	   fz=(atmp(3)/gstep+1-tmp(3))
	   if(tmp(3).lt.1) cycle
           !Approximate Density map

	   apx1=amap(tmp(1),tmp(2),tmp(3))
     *     +(amap(tmp(1)+1,tmp(2),tmp(3))
     *     -amap(tmp(1),tmp(2),tmp(3)))*fx
	   apx2=amap(tmp(1),tmp(2)+1,tmp(3))
     *     +(amap(tmp(1)+1,tmp(2)+1,tmp(3))
     *     -amap(tmp(1),tmp(2)+1,tmp(3)))*fx

	   apy1=apx1+(apx2-apx1)*fy

	   apx1=amap(tmp(1),tmp(2),tmp(3)+1)
     *     +(amap(tmp(1)+1,tmp(2),tmp(3)+1)
     *     -amap(tmp(1),tmp(2),tmp(3)+1))*fx
           apx2=amap(tmp(1),tmp(2)+1,tmp(3)+1)
     *     +(amap(tmp(1)+1,tmp(2)+1,tmp(3)+1)
     *     -amap(tmp(1),tmp(2)+1,tmp(3)+1))*fx

	   apy2=apx1+(apx2-apx1)*fy

           !dmap(ii,jj,kk)=apy1+(apy2-apy1)*fz
           dmap(ii,jj,kk)=(apy1+(apy2-apy1)*fz)

          enddo
         enddo
        enddo

	NumOfGrid=maxp
	gstep=gstep*0.5


	END

c	From Others
	RECURSIVE SUBROUTINE quick_sort(list, order,N)

	! Quick sort routine from:
	! Brainerd, W.S., Goldberg, C.H. & Adams, J.C. (1990) "Programmer's Guide to
	! Fortran 90", McGraw-Hill  ISBN 0-07-000248-7, pages 149-150.
	! Modified by Alan Miller to include an associated integer array which gives
	! the positions of the elements in the original order.

	IMPLICIT NONE
	INTEGER, PARAMETER :: MSIZE=300
	!REAL, DIMENSION (:), INTENT(IN OUT)  :: list
	REAL  list(MSIZE*MSIZE*MSIZE)
	!INTEGER, DIMENSION (:), INTENT(OUT)  :: order
	INTEGER order(MSIZE*MSIZE*MSIZE)

	! Local variable
	INTEGER :: i
	INTEGER :: N

	!DO i = 1, SIZE(list)
	DO i = 1, N
	  order(i) = i
	END DO

	!CALL quick_sort_1(1, SIZE(list))
	CALL quick_sort_1(1, N)

	CONTAINS

	RECURSIVE SUBROUTINE quick_sort_1(left_end, right_end)

	INTEGER, INTENT(IN) :: left_end, right_end

	!     Local variables
	INTEGER             :: i, j, itemp
	REAL                :: reference, temp
	INTEGER, PARAMETER  :: max_simple_sort_size = 6

	IF (right_end < left_end + max_simple_sort_size) THEN
	  ! Use interchange sort for small lists
	  CALL interchange_sort(left_end, right_end)

	ELSE
	  ! Use partition ("quick") sort
	  reference = list((left_end + right_end)/2)
	  i = left_end - 1; j = right_end + 1

	  DO
	    ! Scan list from left end until element >= reference is found
	    DO
	      i = i + 1
	      IF (list(i) >= reference) EXIT
	    END DO
	    ! Scan list from right end until element <= reference is found
	    DO
	      j = j - 1
	      IF (list(j) <= reference) EXIT
	    END DO


	    IF (i < j) THEN
	      ! Swap two out-of-order elements
	      temp = list(i); list(i) = list(j); list(j) = temp
	      itemp = order(i); order(i) = order(j); order(j) = itemp
	    ELSE IF (i == j) THEN
	      i = i + 1
	      EXIT
	    ELSE
	      EXIT
	    END IF
	  END DO

	  IF (left_end < j) CALL quick_sort_1(left_end, j)
	  IF (i < right_end) CALL quick_sort_1(i, right_end)
	END IF

	END SUBROUTINE quick_sort_1


	SUBROUTINE interchange_sort(left_end, right_end)

	INTEGER, INTENT(IN) :: left_end, right_end

	!     Local variables
	INTEGER             :: i, j, itemp
	REAL                :: temp

	DO i = left_end, right_end - 1
	  DO j = i+1, right_end
	    IF (list(i) > list(j)) THEN
	      temp = list(i); list(i) = list(j); list(j) = temp
	      itemp = order(i); order(i) = order(j); order(j) = itemp
	    END IF
	  END DO
	END DO

	END SUBROUTINE interchange_sort

	END SUBROUTINE quick_sort

	FUNCTION fKENDALL(p1,N1,p2,N2,n)
	IMPLICIT INTEGER(I-Z)
        PARAMETER(NDIM=800)
        PARAMETER(MSIZE=300)
	DIMENSION p1(MSIZE*MSIZE*10),p2(MSIZE*MSIZE*10),
     *  indx(MSIZE*MSIZE*10);
	DIMENSION posi1(n,n) ,posi2(n,n);
	LOGICAL ex1(n),ex2(n)

	!init
	ex1(1:n)=.false.
	ex2(1:n)=.false.
	indx(1:maxval(p2(1:N2)))=0;
	do i=1,N2
	 ex2(p2(i))=.true.
	 indx(p2(i))=i;!index of path2
	enddo

	Ntrue=0;
	Nfalse=0;
	do i=1,N1,3
	 o11=i;!order
	 i11=p1(i);!id11
	 if(.not.ex2(i11)) continue;
	 o21=indx(i11);
	 do j=i+1,N1,3
	  o12=j;
	  i12=p1(j);
	  if(.not.ex2(i12)) continue;
	  o22=indx(i12);
	  !o11 < o12 so...
	  if(o21.lt.o22) then
           Ntrue=Ntrue+1;
	  else
           Nfalse=Nfalse+1;
	  endif
	 enddo
	enddo

	fKENDALL=float(2*Ntrue)/float(Ntrue+Nfalse)-1.00

	END
```
