## Supplemental GUI for "CryoFold: determining protein structures and ensembles from cryo-EM data": ThreadCA.f

```
C	***************************************************************
c	Thread sequence on Mainchain models
c
c       #COPYRIGHT
c       (c) 2017 Genki Terashi, Daisuke Kihara and Purdue University
c
c       MAINMAST is a free software for academic and non-commercial users. It is released under the terms of the GNU General Public License Ver.3 (https://www.gnu.org/licenses/gpl-3.0.en.html).
c       Commercial users please contact for alternate licensing.
c       Citation of the following reference should be included in any publication that uses data or results generated by MAINMAST program.
c       (a) Genki Terashi1, and Daisuke Kihara. De novo main-chain modeling for EM maps using MAINMAST. (in submission)
c


	IMPLICIT INTEGER(I-Z)
        character*5 struct(5)
	character*3 aa(-1:20), NAME

	character*500 arg(20),fname_pdb,fname_in,
     *  lbuf,
     *  fname_20aa,fname_spd;

	PARAMETER(NDIM=800)
	PARAMETER(NREPS=50)
	PARAMETER(MSIZE=300)
	PARAMETER(MAXB=5)
	
	COMMON /RES/ EREST, MRES(ndim),KRES(ndim,50),awrca(ndim,420)

	!Genki
	COMMON /CHAINS_XYZ/  XYZ(3,NDIM),GSIDE(3,NDIM),BETA(3,NDIM)
	COMMON /FLAGS/ vflag
	dimension a_xyz(3),b_xyz(3),c_xyz(3),d_xyz(3)
	dimension i_xyz(3),j_xyz(3),p_xyz(3),k_xyz(3)
	dimension l_xyz(3),m_xyz(3),w_xyz(3),n_xyz(3)
	dimension ftmp_xyz(3),xyz_t(3,ndim),xyz_u(3,ndim),xyz_o(3,ndim)
	DIMENSION reps_xyz(3,ndim,NREPS)
	DIMENSION TMP_XYZ(3),AS_XYZ(3),AM_XYZ(3),A_ROT(3,3),I_MAT(3,3)
	COMMON /PEAK/ fw_peak(800,800),peak_xyz(3,800,800)
	DIMENSION buff(10),add_f(0:23),best_f(0:23)
	DIMENSION add_w(0:23),best_w(0:23)
	COMMON /GRIDS_SEQ/    gridseq(ndim)
	DIMENSION ftmp_peak(-7:7,-7:7,-7:7)
	DIMENSION gbase1(3),NumOfGrid1(3), gbase2(3),NumOfGrid2(3)
	!For Mutual Information
	COMMON /MI/	bins(21),Nvox,Npos,Nmap(21),Nmod(21)
	!DIMENSION Emap(MSIZE,MSIZE,MSIZE)
	REAL fin(MSIZE*MSIZE*MSIZE),fori(MSIZE*MSIZE*MSIZE)
        INTEGER order(MSIZE*MSIZE*MSIZE),posi(3,MSIZE*MSIZE*MSIZE)
	DIMENSION After(3,MSIZE*MSIZE*10) !After shift
	DIMENSION Before(3,MSIZE*MSIZE*10) !After shift
	DIMENSION bins1(21),bins2(101),path(MSIZE*MSIZE)
	DIMENSION Nx(21),Ny(101),Nxy(21,101)

	COMMON /GRAPH/ Nbranch(MSIZE*MSIZE*10),IDb(MSIZE*MSIZE*10,MAXB*2),
     *	edged(MSIZE*MSIZE*10,MAXB*2),
     *		nextp(MSIZE*MSIZE*10),
     *		cost(MSIZE*MSIZE*10),
     *  id_edge(MSIZE*MSIZE*10,MAXB*2)
	DIMENSION member(MSIZE*MSIZE*10)
	!TMalign rmsd.f
	double precision fitrms,
     *	fx(3,ndim),fy(3,ndim),fw(ndim),fu(3,3),ft(3)
	COMMON /KRU/ Nround,Nlocal,Ntabu,allow_rate_max,dlocal,dside

	!Command
	character*1 SingleAA(-1:20);
	!PDB
	real  nat_xyz(3,10000),mod_xyz(3,10000),dens(10000)
	real  Waa(-1:20),GapP;
	real  mod_all(3,100000),dens_all(100000)
	DIMENSION ModRg(10);
	integer seqaa(10000),seqss(10000)
	DIMENSION tbl_nat(10000),tbl_mod(10000)
	allocatable dmtx(:,:)
	real,allocatable :: pmtx(:,:)
	!DP
	integer, allocatable :: direc(:,:),prev(:,:)
	real, allocatable :: smtx(:,:)

	!character*3 aa(-1:20), NAME
	data struct /' coil','helix',' turn',' beta','    ?'/
	data aa/ 'BCK','GLY','ALA','SER','CYS','VAL','THR','ILE',
     &		     'PRO','MET','ASP','ASN','LEU',
     &		     'LYS','GLU','GLN','ARG',
     &		     'HIS','PHE','TYR','TRP','CYX'/
	data SingleAA/ 'U','G','A','S','C','V','T','I',
     &		     'P','M','D','N','L',
     &		     'K','E','Q','R',
     &		     'H','F','Y','W','X'/

	real FilterWidth,AverageBond,WeightBond,Wss(1:3)
	real Rsum,AveR,StdR,RefZ(10000),RefD(10000);
	real Tsum,AveT,StdT,TagZ(10000),TagD(10000);
	real up,left,dia
	LOGICAL rmode;
	integer Ali(10000),Pali(10000),BestAli(10000);
	DIMENSION BestWss(3);
	integer StRg,EndRg;
	!for command

	BestSco=-1000000000;
	BestWss=1;
	rmode=.false.

	!Set default
	FilterWidth=1.0;
	BondAvg=3.5;
	BondWeight=0.9;
	rmode=.false.
	
	if(iargc().ge.2) then
	 do i=1,iargc()
	  call getarg(i,arg(i))
	  lbuf=arg(i);
	  if(lbuf(1:2).eq."-i")then
	   call getarg((i+1),arg(i+1))
	   fname_in=arg(i+1);
	   write(*,*),"#INPUT=",trim(fname_in)
	   cycle;
	  endif
	  if(lbuf(1:2).eq."-a")then
	   call getarg((i+1),arg(i+1))
	   fname_20aa=arg(i+1);
	   write(*,*),"#20AA file=",trim(fname_20aa)
	   cycle;
	  endif
	  if(lbuf(1:4).eq."-spd")then
	   call getarg((i+1),arg(i+1))
	   fname_spd=arg(i+1);
	   write(*,*),"#SPD3=",trim(fname_spd)
	   cycle;
	  endif
	  if(lbuf(1:3).eq."-fw")then
           call getarg(i+1,arg(i+1))
           read(arg(i+1),*),FilterWidth;
           cycle;
          endif
	  if(lbuf(1:3).eq."-Ab")then
           call getarg(i+1,arg(i+1))
           read(arg(i+1),*),BondAvg;
           cycle;
          endif
	  if(lbuf(1:3).eq."-Wb")then
           call getarg(i+1,arg(i+1))
           read(arg(i+1),*),BondWeight;
           cycle;
          endif
	  if(lbuf(1:3).eq."-r")then
	   rmode=.true.
           cycle;
          endif
	 enddo
	else
	 write(*,*),"Usage: ThreadCA "
     *   ,"-i   [OUT file from MAINMAST] "
     *   ,"-a   [20AA.param] "
     *   ,"-spd [*.spd3] "
     *   ," (option)"
	 write(*,*),"Option ver1.0:"
	 write(*,*)," -i   [file] : Result file of MAINMAST "
	 write(*,*)," -a   [file] : 20AA.param"
	 write(*,*)," -spd [file] : Result of SPIDER2"
	 write(*,*)," -fw [f]   : Filter width def=1.0"
	 write(*,*)," -Ab [f]   : Average length of CA-CA Bond def=3.5"
	 write(*,*)," -Wb [f]   : Weight of Bond score def=0.9"
	 write(*,*)," -r : Reverse mode, reverse mainchain order"
	 stop
	endif
	
	
	
	GapP=-10000;
	call read_mainchain_model(fname_in,mod_all,NRmod,dens_all,.true.,
     *  ModRg,Nmdl);!main chain only
	call read_spd3(fname_spd,singleaa,SeqAA,SeqSS,Nseq);
	call read_20aaparam(fname_20aa,aa,Waa);

	MaxResNum=ModRg(1);
	do i=2,Nmdl
	 if(ModRg(i)-ModRg(i-1).gt.MaxResNum)then
	  MaxResNum=ModRg(i)-ModRg(i-1);
	 endif
	enddo
	write(*,*),"#MaxResNum=",MaxResNum;
	
	allocate(dmtx(0:MaxResNum,0:MaxResNum))
	allocate(pmtx(0:MaxResNum,0:Nseq))
	allocate(smtx(0:MaxResNum,0:Nseq))
	allocate(prev(0:MaxResNum,0:Nseq))
	allocate(direc(0:MaxResNum,0:Nseq))


	!Multi Models!!

	do mdlid=1,Nmdl
	
	 if(mdlid.eq.1)then
	  length=ModRg(mdlid);
	  StRg=1;
	  EndRg=ModRg(mdlid);
	 else
	  length=ModRg(mdlid)-ModRg(mdlid-1);
	  StRg=ModRg(mdlid-1)+1;
	  EndRg=ModRg(mdlid);
	 endif
	
	 mod_xyz(:,1:length)=mod_all(:,StRg:EndRg);
	 dens(1:length)=dens_all(StRg:EndRg);
	 NRmod=length;
	 write(*,*),"#model",mdlid,"Start:End",StRg,EndRg
	 !write(*,*),mod_xyz(:,1)
	 !write(*,*),mod_xyz(:,length)
	
	 if(rmode)then
	  !reverse mod_xyz and dens
	  do i=NRmod,1,-1
	   nat_xyz(:,NRmod-i+1)=mod_xyz(:,i)
	   TagD(NRmod-i+1)=dens(i)
	  enddo
	  mod_xyz(:,1:NRmod)=nat_xyz(:,1:NRmod)
	  dens(1:NRmod)=TagD(1:NRmod)
	 endif
	 Np=NRmod
	 Nres=Nseq

	!Distance Matrix
	dmtx=0;
	do i=1,Np
	 do j=i+1,Np
	  ftmp_xyz=mod_xyz(:,i)-mod_xyz(:,j);
	  dmtx(i,j)=sqrt(dot_product(ftmp_xyz,ftmp_xyz));
	  dmtx(j,i)=dmtx(i,j);
	 enddo
	enddo
	!New check turn loop
	do i=1,Np
	 do j=i+2,Np
	  ftmp=dmtx(i,j)
	  do k=i+1,j-1
	   if(dmtx(i,k).gt.ftmp)ftmp=dmtx(i,k);
	   if(dmtx(j,k).gt.ftmp)ftmp=dmtx(j,k);
	  enddo
	  if(ftmp.ne.dmtx(i,j))then
	   !write(*,*),"Re-new",i,j,dmtx(i,j),ftmp;
	  endif
	  dmtx(i,j)=ftmp;
	  dmtx(j,i)=ftmp;
	 enddo
	enddo
	!Filterling
	Rsum=0;
	do i=1,NRmod
	 ftmp=0;
	 do j=i,NRmod
	  d=dmtx(i,j);
	  if(d > 6.0) exit;
  	  FilterG=(1.00/FilterWidth)*exp(-d*d/(2*FilterWidth*FilterWidth));
  	  ftmp=ftmp+dens(j)*FilterG;
	 enddo
	 do j=i-1,1,-1
	  d=dmtx(i,j);
	  if(d > 6.0) exit;
  	  FilterG=(1.00/FilterWidth)*exp(-d*d/(2*FilterWidth*FilterWidth));
  	  ftmp=ftmp+dens(j)*FilterG;
	 enddo
	 RefD(i)=ftmp;
	 Rsum=Rsum+ftmp;
	enddo

	AveR=Rsum/float(NRmod);
	ftmp=0;
	do i=1,NRmod
 	 ftmp=ftmp+(RefD(i)-AveR)**2;
	enddo
	StdR=sqrt(ftmp/float(NRmod));
	do i=1,NRmod
 	 RefZ(i)=(RefD(i)-AveR)/StdR;
	enddo

	!write(*,*),"#Ave=",AveR,"STD=",StdR;

	!Multiple!!!
	!H:1 E:2 C:3
	Wss(3)=1.00;
	do i=0,3
	 Wss(1)=1.000+0.100*i;
	do j=0,3
	 Wss(2)=0.700+0.100*j
	 !write(*,*),Wss(1),Wss(2),Wss(3);

	 !Estimate Sum
	 ftmp=0;
	 do k=1,Nseq
 	  ftmp=ftmp+Waa(seqaa(k))*Wss(seqss(k));
	  TagD(k)=Waa(seqaa(k))*Wss(seqss(k));
	 enddo
	 AveT=ftmp/float(Nseq);
	 ftmp=0;
	 do k=1,Nseq
	  ftmp=ftmp+(TagD(k)-AveT)**2;
	 enddo
	 StdT=sqrt(ftmp/float(Nseq));
	 do k=1,Nseq
	  TagZ(k)=(TagD(k)-AveT)/StdT;
	 enddo

	 !write(*,*),"#Ave=",AveR,"STD=",StdR;
	 !write(*,*),"#Ave=",AveT,"STD=",StdT;


	!SetUp Amtx

	 do ii=1,Np
	 do jj=1,Nres
	  diff=abs(TagZ(jj)-RefZ(ii));
	  pmtx(ii,jj)=1.0-diff;
	  !write(*,*),pmtx(ii,jj);
	 enddo
	 enddo

	!!DP!!
	 direc=0;
	 smtx=0;
	 Prev=0;
	 !Direction 0:stop, 1:up, 2:left, 3:dia
	
	 !Init 0 lines
	 do ii=0,Np
 	  smtx(ii,0)=0;
 	  direc(ii,0)=2;
 	  Prev(ii,0)=0;
	 enddo
	 direc(0,0)=0;
	 smtx(0,0)=0;
	 do ii=1,Nres
 	  smtx(0,ii)=smtx(0,ii-1)+GapP;
 	  direc(0,ii)=1;
 	  Prev(0,ii)=0;
	 enddo

	!Fill====================
	 do p=1,Np
 	  do r=1,Nres
  	   up=smtx(p,r-1)+GapP;
  	   left=smtx(p-1,r);

	   !Aligned!!=dia
  	   nowx=p-1;
  	   nowy=r-1;
  	   nowx=Prev(nowx,nowy);
  	   if(direc(nowx,nowy).eq.0)then
   	    dia=smtx(p-1,r-1)+Pmtx(p,r);
  	   else
   	    d=DMTX(p,nowx);
   	    add=Pmtx(p,r)-(abs(d-BondAvg))*BondWeight;
   	    dia=smtx(p-1,r-1)+add;
  	   endif

  	   if(dia.ge.left.and.dia.ge.up)then
   	    smtx(p,r)=dia;
   	    direc(p,r)=3;
  	    Prev(p,r)=p;
	   else if(up.ge.left) then
   	     smtx(p,r)=up;
 	     direc(p,r)=1;
	     Prev(p,r)=Prev(p,r-1);
	   else
	     smtx(p,r)=left;
	     direc(p,r)=2;
	     Prev(p,r)=Prev(p-1,r);
	   endif
	   !if(r.eq.Nres)then
	   ! write(*,*),p,dia,up,left,smtx(p,r);
	   !endif
	  enddo
	 enddo

	 !TRACE BACK
	 nowx=Np;
	 nowy=Nres;
	 Nali=0;
	 do
 	  if(direc(nowx,nowy).eq.0) exit;
 	  if(direc(nowx,nowy).eq.3)then
  	   Nali=Nali+1;
 	   Pali(Nali)=nowx;
 	   Ali(nowy)=nowx;
 	   nowx=nowx-1;
 	   nowy=nowy-1;
 	  else if(direc(nowx,nowy).eq.1)then
 	   nowy=nowy-1;
 	  else if(direc(nowx,nowy).eq.2)then
 	   nowx=nowx-1;
 	  endif
	 enddo

	 !Record Best
	 write(*,'("model ",I3,2f10.3," Wss=",3f5.1)'),
     *	 mdlid,smtx(Np,Nres),maxval(smtx),Wss
	 !write(*,*),"model   ",mdlid,smtx(Np,Nres),maxval(smtx),"W=",Wss
 	 if(BestSco.lt.smtx(Np,Nres))then
 	 !if(BestSco.lt.maxval(smtx))then
 	  BestAli=Ali;
 	  BestSco=smtx(Np,Nres);
 	  !BestSco=maxval(smtx);
 	  BestWss=Wss;
	  iBestMid=mdlid
 	 endif


	enddo
	enddo

	enddo


	!Show Best Alignment
	Wss=BestWss;
	Ali=BestAli;
	write(*,*),"MODEL",iBestMid,BestSco,
     *  " Wh=",Wss(1)," We=",Wss(2)," Wc=",Wss(3)
	mdlid=iBestMid;

	if(mdlid.eq.1)then
	  length=ModRg(mdlid);
	  StRg=1;
	  EndRg=ModRg(mdlid);
	 else
	  length=ModRg(mdlid)-ModRg(mdlid-1);
	  StRg=ModRg(mdlid-1)+1;
	  EndRg=ModRg(mdlid);
	 endif
	 NRmod=length;
	 mod_xyz(:,1:length)=mod_all(:,StRg:EndRg);
	 dens(1:length)=dens_all(StRg:EndRg);

	 if(rmode)then
	  !reverse mod_xyz and dens
	  do i=NRmod,1,-1
	   nat_xyz(:,NRmod-i+1)=mod_xyz(:,i)
	   TagD(NRmod-i+1)=dens(i)
	  enddo
	  mod_xyz(:,1:NRmod)=nat_xyz(:,1:NRmod)
	  dens(1:NRmod)=TagD(1:NRmod)
	 endif
	 Np=NRmod
	 Nres=Nseq

	Natm=1;
	do ii=1,Nres
 	 n=Ali(ii);
 	 if(n.ne.0)then
	 write(*,'(A4,i7,A6,A3,A2,I4,A4,3f8.3,2f6.2)')
     *   ,"ATOM",Natm,"  CA  ",aa(seqaa(ii))," A"
     *   ,ii,'    ',mod_xyz(1:3,n),1.00,1.00
 	 Natm=Natm+1;
 	 endif
	enddo

	END

	SUBROUTINE READ_SPD3(filename,singleaa,seq,seqss,Nres)
	IMPLICIT INTEGER(I-Z)
	character*500 filename,buf;
	character*1 SingleAA(-1:20);
	character bufaa,bufss
	!PDB
	integer seq(10000),seqss(10000)
	Nres=0;
	write(*,*),"#reading:",trim(filename);
	OPEN(UNIT=22,FILE=filename,STATUS='OLD') !New!!!
	do
	 read(22,'(A80)',END=999) ,buf
	 if(buf(1:1).eq.'#') cycle;
	 read(buf,*),Nres,bufaa,bufss
	 j=-1;
	 do i=0,20
	  if(singleaa(i).eq.bufaa)then
	   j=i;
	   exit;
	  endif
	 enddo
	 if(j.eq.-1) cycle;
	 seq(Nres)=j;
	 if(bufss.eq.'C') seqss(Nres)=3;
	 if(bufss.eq.'H') seqss(Nres)=1;
	 if(bufss.eq.'E') seqss(Nres)=2;
	 !write(*,*),"#SPD3",Nres,seq(Nres),seqss(Nres)
	enddo
  999	close(22)
	END


	SUBROUTINE READ_20AAPARAM(filename,aa,Waa)
	IMPLICIT INTEGER(I-Z)
	character*500 filename,buf;
	character*3 aa(-1:20), NAME
	real cd(3,10000),Waa(-1:20);
	logical flag;!true=main chain only
	Nres=0;
	Waa=0;
	write(*,*),"#reading:",trim(filename)
	OPEN(UNIT=22,FILE=filename,STATUS='OLD') !New!!!
	do
         read(22,'(A80)',END=999) ,buf
	 !write(*,*),buf
	 j=-1;
	 do i=0,20
	  if(aa(i).eq.buf(1:3))then
	   j=i;
	   exit;
	  endif
	 enddo
	 if(j.eq.-1) cycle;
	 read(buf(4:),*),Waa(j)
	 !write(*,*),"#AA",j,Waa(j)
	enddo
  999	close(22)
	END


	SUBROUTINE READ_MAINCHAIN(filename,cd,Nres,dens,flag)
	IMPLICIT INTEGER(I-Z)
	character*500 filename,buf;
	real cd(3,10000),dens(10000);
	logical flag;!true=main chain only
	Nres=0;
	write(*,*),"#reading:",trim(filename)
	OPEN(UNIT=22,FILE=filename,STATUS='OLD') !New!!!
	do
         read(22,'(A80)',END=999) ,buf
	 if(buf(1:4).ne."ATOM") cycle;
	 if(flag.and.buf(14:16).ne."N  ".and.
     *      buf(14:16).ne."CA ".and.
     *      buf(14:16).ne."C  ".and.
     *      buf(14:16).ne."O  "
     *   )then
	 cycle;!Main chain mode
	endif
	 !write(*,*),buf
	 Nres=Nres+1;
	 read(buf(31:38),*),cd(1,Nres)
	 read(buf(39:46),*),cd(2,Nres)
	 read(buf(47:54),*),cd(3,Nres)
	 read(buf(61:66),*),dens(Nres)
	 !write(*,*),cd(:,Nres)
	enddo
  999	close(22)
	write(*,*),"#Nres=",Nres
	END


	SUBROUTINE READ_MAINCHAIN_MODEL(filename,cd,Nres,dens,flag,
     *  MdlRg,Nmdl)
	IMPLICIT INTEGER(I-Z)
	character*500 filename,buf;
	real cd(3,100000),dens(100000);
	DIMENSION MdlRg(10);
	logical flag;!true=main chain only
	Nres=0;
	Nmdl=0;
	write(*,*),"#reading:",trim(filename);
	OPEN(UNIT=22,FILE=filename,STATUS='OLD') !New!!!
	do
         read(22,'(A80)',END=999) ,buf

	 if(buf(1:6).eq."ENDMDL") then
	  Nmdl=Nmdl+1;
	  MdlRg(Nmdl)=Nres;
	  write(*,*),"#Reading MODEL",Nmdl,Nres
	 endif
	 if(buf(1:4).ne."ATOM") cycle;
	 if(flag.and.buf(14:16).ne."N  ".and.
     *      buf(14:16).ne."CA ".and.
     *      buf(14:16).ne."C  ".and.
     *      buf(14:16).ne."O  "
     *   )then
	 cycle;!Main chain mode
	 endif
	 !write(*,*),buf
	 Nres=Nres+1;
	 read(buf(31:38),*),cd(1,Nres)
	 read(buf(39:46),*),cd(2,Nres)
	 read(buf(47:54),*),cd(3,Nres)
	 read(buf(61:66),*),dens(Nres)
	 !write(*,*),cd(:,Nres)
	enddo
  999	close(22)
	write(*,*),"#Nres=",Nres
	END
```
