## Supplemental GUI for "CryoFold: determining protein structures and ensembles from cryo-EM data": MDFF.py

```
import subprocess
import shlex

class MDFF:

    def getSpecificsMDFF(self, psfFile, pdbFile, mapFile, GSCALE, NUMSTEPS):
        if not GSCALE.isdigit():
            GSCALE = 0.3
        if not NUMSTEPS.isdigit():
            NUMSTEPS = 50000
        self.psfFile = psfFile
        self.pdbFile = pdbFile
        self.mapFile = mapFile
        self.GSCALE = GSCALE
        self.NUMSTEPS = NUMSTEPS

    def writeMDFF(self):
        f = open('outputs/tmp/mdff.tcl', 'w')

        f.write("package require mdff\n")
        f.write("package require cispeptide\n")
        f.write("package require chirality\n")
        f.write("package require ssrestraints\n")
        f.write("\n")
        f.write("#Generating Simulated Density\n")
        f.write("\n")
        f.write("#Generating the grid file\n")
        f.write("mdff griddx -i " + self.mapFile + " -o " + "outputs/MAP.dx\n")
        f.write("\n")
        f.write("#Generating the file on which the grid potential will apply to\n")
        f.write('mdff gridpdb -psf ' + str(self.psfFile) + ' -pdb ' + str(self.pdbFile) + ' -seltext "backbone" -o outputs/bb_grid.pdb\n')
        f.write('mdff gridpdb -psf ' + str(self.psfFile) + ' -pdb ' + str(self.pdbFile) + ' -seltext "noh" -o outputs/noh_grid.pdb\n')
        f.write("\n")
        f.write("#Generating the secondary structure restrain files\n")
        f.write("ssrestraints -psf " + str(self.psfFile) + " -pdb " + str(self.pdbFile) + " -o outputs/system_extrabonds.txt -hbonds\n")
        f.write("\n")
        f.write("#Chirality and Cis-peptide restraints\n")
        f.write("mol new " + self.psfFile + "\n")
        f.write("mol addfile " + self.pdbFile + "\n")
        f.write("cispeptide restrain -o outputs/system_cispeptide.txt\n")
        f.write("chirality restrain -o outputs/system_chirality.txt\n")
        f.write("\n")
        f.write("#generating the configuration file\n")
        f.write("mdff setup -o outputs/Mdff -psf " + str(self.psfFile) + " -pdb " + str(self.pdbFile) + " -griddx MAP.dx -gridpdb bb_grid.pdb -extrab {system_extrabonds.txt system_cispeptide.txt system_chirality.txt} -gscale " + str(self.GSCALE) + " -numsteps " + str(self.NUMSTEPS) + " -gbis\n")
        f.write("\n")
        f.write("###generating continution files###\n")
        f.write("touch ../Mdff-step2.namd\n")
        f.write("head -n 17 Mdff-step1.namd >> outputs/Mdff-step2.namd\n")
        f.write('echo "set OUTPUTNAME Mdff-step2" >> outputs/Mdff-step2.namd\n')
        f.write('echo "set INPUTNAME Mdff-step1" >> outputs/Mdff-step2.namd\n')
        f.write("tail -n 23 Mdff-step1.namd >> outputs/Mdff-step2.namd\n")
        f.write("\n")
        f.write('sed -i -e "s/bb_grid.pdb/noh_grid.pdb/g" outputs/Mdff-step2.namd\n')
        f.write("\n")
        f.write("quit\n")

        f.close()

        subprocess.call(shlex.split("""vmd -dispdev text -e outputs/tmp/mdff.tcl"""))
```
