## Supplemental GUI for "CryoFold: determining protein structures and ensembles from cryo-EM data": MDFFScreen.py

```
#!/usr/bin/env python

import os
from kivy.app import App
from kivy.lang import Builder
from kivy.uix.screenmanager import ScreenManager, Screen
from kivy.uix.label import Label
from kivy.uix.textinput import TextInput
from kivy.uix.popup import Popup
from MDFF import MDFF
from LoadDialog import LoadDialog
from AlertDialog import AlertDialog

Builder.load_file('MDFF_GUI/MDFF_Screen.kv')

class MDFFScreen(Screen):

    def dismiss_popup(self):
        self._popup.dismiss()

    def show_load(self, fileinput):
        content = LoadDialog(load=self.load, cancel=self.dismiss_popup, fileinput=fileinput)
        self._popup = Popup(title="Load file", content=content,
                            size_hint=(0.9, 0.9))
        self._popup.open()

    def load(self, path, filename, fileinput):
        with open(os.path.join(path, filename[0])) as stream:
            getattr(self, fileinput).text = stream.name

        self.dismiss_popup()

    def close():
        self.dismiss_popup()

    def saveMDFF(self, GSCALE, NUMSTEPS):
        namd_mdff = MDFF()
        namd_mdff.getSpecificsMDFF(self.psfFile.text, self.pdbFile.text, self.mapFile.text, GSCALE, NUMSTEPS)
        namd_mdff.writeMDFF()
```
