## Supplemental GUI for "CryoFold: determining protein structures and ensembles from cryo-EM data": calculate_contacts_native.py

```
#! /usr/bin/env python

import numpy as np
import mdtraj


def calculate_contacts_native(file_name, distance_max):
    pdb = mdtraj.load_pdb(file_name)
    #distances, contact_pairs = mdtraj.compute_contacts(pdb,contacts='all',scheme='closest-heavy',periodic=False)
    distances, contact_pairs = mdtraj.compute_contacts(pdb,contacts='all',scheme='ca',periodic=False)

    #get distances closer than 8 angstrongs
    indices = distances < float(distance_max)
    (a,b) =  indices.shape
    native_contacts = contact_pairs[indices[0],:]
    with open("./outputs/tmp/contacts.dat", "w") as file:
        for i in native_contacts:
            a,b = i
            if (abs(a-b)) > 6:
                file.write(str(a+1) + ' CA ' + str(b+1) + ' CA ' + str(8.0) + '\n')
```
