## Supplemental GUI for "CryoFold: determining protein structures and ensembles from cryo-EM data": create_template.py

```
def create_template(nb_replicas, nb_steps, block_size, sequence, init_temp, final_temp, file_name, ss_ratio, contacts_ratio):
    output_file = open('./outputs/meld_script.py', 'w')
    output_file.write(
        """#!/usr/bin/env python
# encoding: utf-8

import numpy as np
from meld.remd import ladder, adaptor, master_runner
from meld import comm, vault
from meld import system
from meld import parse
import meld.system.montecarlo as mc
from meld.system.restraints import LinearRamp,ConstantRamp
from collections import namedtuple
import glob


N_REPLICAS = {}
N_STEPS = {}
BLOCK_SIZE = {}


def gen_state(s, index):
    pos = s._coordinates
    pos = pos - np.mean(pos, axis=0)
    vel = np.zeros_like(pos)
    alpha = index / (N_REPLICAS - 1.0)
    energy = 0
    #return system.SystemState(pos, vel, alpha, energy,[0,0,0])
    return system.SystemState(pos, vel, alpha, energy)


def get_dist_restraints(filename, s, scaler):
    dists = []
    rest_group = []
    lines = open(filename).read().splitlines()
    lines = [line.strip() for line in lines]
    for line in lines:
        if not line:
            dists.append(s.restraints.create_restraint_group(rest_group, 1))
            rest_group = []
        else:
            cols = line.split()
            i = int(cols[0])
            name_i = cols[1]
            j = int(cols[2])
            name_j = cols[3]
            dist = float(cols[4]) / 10.

            rest = s.restraints.create_restraint('distance', scaler,LinearRamp(0,100,0,1),
                                                 r1=0.0, r2=0.0, r3=dist, r4=dist+0.2, k=250,
                                                 atom_1_res_index=i, atom_2_res_index=j,
                                                 atom_1_name=name_i, atom_2_name=name_j)
            rest_group.append(rest)
    return dists

def setup_system():
    # load the sequence
    sequence = parse.get_sequence_from_AA1(filename='{}')
    n_res = len(sequence.split())

    # build the system
    p = system.ProteinMoleculeFromSequence(sequence)
    b = system.SystemBuilder(forcefield="ff14sbside")
    s = b.build_system_from_molecules([p])
    s.temperature_scaler = system.GeometricTemperatureScaler(0, 0.6, {}, {})

    #
    # Secondary Structure
    #
    ss_scaler = s.restraints.create_scaler('constant')
    ss_rests = parse.get_secondary_structure_restraints(filename='{}', system=s,ramp=LinearRamp(0,100,0,1), scaler=ss_scaler,
            torsion_force_constant=2.5, distance_force_constant=2.5)
    n_ss_keep = int(len(ss_rests) * {}) #We enforce {}% of restrains
    s.restraints.add_selectively_active_collection(ss_rests, n_ss_keep)

    #
    # Distance Restraints
    #
    dist_scaler = s.restraints.create_scaler('nonlinear', alpha_min=0.4, alpha_max=1.0, factor=4.0)

    #Contacts from initial model
    distances = get_dist_restraints('./tmp/contacts.dat',s,dist_scaler)
    n_distances = int(len(distances)*{})
    print(distances,n_distances)
    s.restraints.add_selectively_active_collection(distances,n_distances)

    # setup mcmc at startup
    movers = []
    n_atoms = s.n_atoms
    for i in range(1, n_res + 1):
        n = s.index_of_atom(i, 'N') - 1
        ca = s.index_of_atom(i, 'CA') - 1
        c = s.index_of_atom(i, 'C') - 1

        mover = mc.DoubleTorsionMover(n, ca, list(range(ca, n_atoms)),
                                      ca, c, list(range(c, n_atoms)))

        movers.append((mover, 1))

    sched = mc.MonteCarloScheduler(movers, n_res * 60)

    # create the options
    options = system.RunOptions()
    options.implicit_solvent_model = 'gbNeck2'
    options.use_big_timestep = False
    options.use_bigger_timestep = True
    options.cutoff = 1.8

    options.use_amap = False
    options.amap_alpha_bias = 1.0
    options.amap_beta_bias = 1.0
    options.timesteps = 11111
    options.minimize_steps = 20000
    #options.min_mc = sched

    # create a store
    store = vault.DataStore(s.n_atoms, N_REPLICAS, s.get_pdb_writer(), block_size=BLOCK_SIZE)
    store.initialize(mode='w')
    store.save_system(s)
    store.save_run_options(options)

    # create and store the remd_runner
    l = ladder.NearestNeighborLadder(n_trials=100)
    policy = adaptor.AdaptationPolicy(2.0, 50, 50)
    a = adaptor.EqualAcceptanceAdaptor(n_replicas=N_REPLICAS, adaptation_policy=policy)

    remd_runner = master_runner.MasterReplicaExchangeRunner(N_REPLICAS, max_steps=N_STEPS, ladder=l, adaptor=a)
    store.save_remd_runner(remd_runner)

    # create and store the communicator
    c = comm.MPICommunicator(s.n_atoms, N_REPLICAS)
    store.save_communicator(c)

    # create and save the initial states
    states = [gen_state(s, i) for i in range(N_REPLICAS)]
    store.save_states(states, 0)

    # save data_store
    store.save_data_store()

    return s.n_atoms


setup_system()
        """.format(nb_replicas, nb_steps, block_size, sequence, init_temp, final_temp, file_name, ss_ratio, float(ss_ratio)*100, contacts_ratio)
        )
    output_file.close()
```
