## Supplemental GUI for "CryoFold: determining protein structures and ensembles from cryo-EM data": MeldScreen.py

```
from kivy.properties import ObjectProperty, StringProperty
from kivy.uix.boxlayout import BoxLayout
from kivy.uix.gridlayout import GridLayout
from kivy.uix.popup import Popup
from kivy.lang import Builder
from LoadDialog import LoadDialog
from AlertDialog import AlertDialog
from calculate_contacts_native import calculate_contacts_native
from kivy.uix.screenmanager import Screen, SlideTransition
from kivy.uix.button import Button
from kivy.uix.textinput import TextInput
from kivy.uix.label import Label

from create_template import create_template

import os
import subprocess

Builder.load_file('./MELD_GUI/meldscreen.kv')

class MeldScreen(Screen):

    def dismiss_popup(self):
        self._popup.dismiss()

    def show_load(self, fileinput):
        content = LoadDialog(load=self.load, cancel=self.dismiss_popup, fileinput=fileinput)
        self._popup = Popup(title="Load file", content=content,
                            size_hint=(0.9, 0.9))
        self._popup.open()

    def show_alert(self, message):
        content = AlertDialog(close=self.dismiss_popup, message=message)
        self._popup = Popup(title="Alert", content=content,
                            size_hint=(0.5, 0.2))
        self._popup.open()

    def load(self, path, filename, fileinput):
        with open(os.path.join(path, filename[0])) as stream:
            getattr(self, fileinput).text = stream.name
        self.dismiss_popup()

    def close():
        self.dismiss_popup()


    def change_button_run_name(self, widget, message):
        widget.text = message

    def run_command(self, widget):
        if self.ss_ratio.text == '':
            self.ss_ratio.text = '0.70'
        if self.distance_max.text == '':
            self.distance_max.text = '0.8'
        if self.contacts_ratio.text == '':
            self.contacts_ratio.text = '0.15'
        if self.nb_replicas.text == '':
            self.nb_replicas.text = '30'
        if self.nb_steps.text == '':
            self.nb_steps.text = '10000'
        if self.block_size.text == '':
            self.block_size.text = '100'
        if self.init_temp.text == '':
            self.init_temp.text = '300'
        if self.final_temp.text == '':
            self.final_temp.text = '450'
        if self.mdff_mainmast_file_name.text.split('.')[-1] != 'pdb':
            self.show_alert('Wrong Format')
        else:
            calculate_contacts_native(self.mdff_mainmast_file_name.text, float(self.distance_max.text))
            create_template(
                self.nb_replicas.text,
                self.nb_steps.text,
                self.block_size.text,
                self.sequence.text,
                self.init_temp.text,
                self.final_temp.text,
                self.ss_prediction_file_name.text,
                self.ss_ratio.text,
                self.contacts_ratio.text,
                )
            self.manager.transition = SlideTransition(direction='right')
            self.manager.current = 'menu'
        widget.text = 'Run'
```
