## Supplemental GUI for "CryoFold: determining protein structures and ensembles from cryo-EM data": README.txt

The following are instructions for installing and starting the CryoFold GUI on a Linux workstartion or mac computer using a bash terminal.

First, make sure you have anaconda installed and it is in your path.  Type conda in command line as shown below (commands that need to be entered in a bash terminal are indicated by the prompt "$" henceforth).

$ conda --version

This should output the version number currently installed.  If the command is not found then the anaconda bin is not in your path.  If this is the case run the following

$ export PATH=/path/to/anaconda3/bin:$PATH

Or place the above line of code in your .bashrc file and restart your terminal.

If you are using anaconda2 make the corresponding chage in the above line of code.  Also you will need to manually change the below line of code in the install.sh bash script

yes | conda create -n cryofold python=3.5 kivy numpy mdtraj meld-cuda75 

to

yes | conda create -n cryofold python=2.7 kivy numpy mdtraj meld-cuda75

Next you will want to make sure the GNU fortran compiler is installed and is in your path.  To do this run the following command in your bash terminal.

$ gfortran --version

If this command is not found visit one of the following links to download and install the the source code for gfortran on your respecitve machine.

https://gcc.gnu.org/wiki/GFortranBinaries

Follow the instructions for installation and make sure it is in your path by reruning the above line of code.  


To install CryoFold on a Linux machine, please run 

$ bash install_linux.sh

To install CryoFold on a mac, please run 

$ bash install_mac.sh


Your CryoFold environment is now setup.

To start CryoFold, please run 

$ ./start.sh

Usage instructions for using the CryoFold GUI are located in the supplementary information of CryoFold: Ab-initio structure determination from electron density maps using molecular dynamics on pages 15 to 23.
