## Supplemental GUI for "CryoFold: determining protein structures and ensembles from cryo-EM data": TaMD.py

```
# IPython log file

class TaMD:

	def getSpecifics(self, jobName, NO, PSF, PDB, TEMP, TIME_IN_NS, MIN, RESTFREQ, DCDFREQ, XSTFREQ, OUTEN, OUTP, TMDFile, TMDk, TMDOutFreq, TMDLastStep, fName):
		if fName == '':
			fName = 'TMD.conf'
		if jobName == '':
			jobName = "TMD"
		if not NO.isdigit():
			NO = 0
		if not TEMP.isdigit():
			TEMP = 310
		if not MIN.isdigit():
			MIN = 1000
		if not TIME_IN_NS.isdigit():
			TIME_IN_NS = 5
		if not RESTFREQ.isdigit():
			RESTFREQ = 5000
		if not DCDFREQ.isdigit():
			DCDFREQ = 5000
		if not XSTFREQ.isdigit():
			XSTFREQ = 5000
		if not OUTEN.isdigit():
			OUTEN = 5000
		if not OUTP.isdigit():
			OUTP = 5000
		if not TMDk:
			TMDk = 200
		if not TMDOutFreq:
			TMDOutFreq = 5000
		if not TMDLastStep:
			TMDLastStep = 50000

		self.jobName = jobName
		self.NO = NO
		self.PSF = PSF
		self.PDB = PDB
		self.TEMP = TEMP
		self.MIN = MIN
		self.TIME_IN_NS = TIME_IN_NS
		self.RESTFREQ = RESTFREQ
		self.DCDFREQ = DCDFREQ
		self.XSTFREQ = XSTFREQ
		self.OUTEN = OUTEN
		self.OUTP = OUTP
		self.TMDFile = TMDFile
		self.TMDk = TMDk
		self.TMDOutFreq = TMDOutFreq
		self.TMDLastStep = TMDLastStep
		self.fName = fName

	def writeNAMD(self):
		#self.getSpecifics()

		f = open('outputs/'  + self.fName, "w")

		f.write("#############################################################\n")
		f.write("## JOB DESCRIPTION                                         ##\n")
		f.write("#############################################################\n")
		f.write("\n")
		f.write("# keeping the heavy atoms of the protein restrained\n")
		f.write("\n")
		f.write("#############################################################\n")
		f.write("## ADJUSTABLE PARAMETERS                                   ##\n")
		f.write("#############################################################\n")
		f.write("\n")
		f.write("set name " + self.jobName + "\n")
		f.write("set NO " + str(self.NO) + "\n")
		f.write("\n")
		f.write("set outputName $name.$NO" + "\n")
		f.write("if {$NO == 0} {\n")
		f.write("    set inputName ''\n")
		f.write("} else {\n")
		f.write("    set inputName $name.[expr $NO - 1]\n")
		f.write("}\n")
		f.write("\n")
		f.write("structure   " + self.PSF + "\n")
		f.write("coordinates " + self.PDB + "\n")
		f.write("outputName  $outputName\n")
		f.write("\n")
		f.write("set temperature		" + str(self.TEMP) + "     ;# K\n")
		f.write("set minimize 		" + str(self.MIN) + "\n")
		f.write("set time	        " + str(self.TIME_IN_NS) + "	;# ns\n")
		f.write("set timestep	        2.0\n")
		f.write("\n")
		f.write("proc get_first_ts { xscfile } {\n")
		f.write("	set fd [open $xscfile r]\n")
		f.write("	gets $fd\n")
		f.write("	gets $fd\n")
		f.write("	gets $fd line\n")
		f.write("	set ts [lindex $line 0]\n")
		f.write("	close $fd\n")
		f.write("	return $ts\n")
		f.write("}\n")
		f.write("\n")
		f.write("if {$NO != 0} {\n")
		f.write("	bincoordinates	$inputName.restart.coor\n")
		f.write("	binvelocities	$inputName.restart.vel\n")
		f.write("	set firsttime	0\n")
		f.write("    extendedSystem  $inputName.restart.xsc\n")
		f.write("} else {\n")
		f.write("	set firsttime	0\n")
		f.write("	temperature	$temperature\n")
		f.write("}\n")
		f.write("\n")
		f.write("firsttimestep $firsttime\n")
		f.write("\n")
		f.write("#############################################################\n")
		f.write("## SIMULATION PARAMETERS                                   ##\n")
		f.write("#############################################################\n")
		f.write("\n")
		f.write("# Input\n")
		f.write("paraTypeCharmm	on\n")
		f.write("parameters          par_all36_prot.prm \n")
		f.write("\n")
		f.write("# Implicit Solvent\n")
		f.write("gbis                on\n")
		f.write("alphaCutoff         14\n")
		f.write("ionConcentration    0.1\n")
		f.write("\n")
		f.write("# Force-Field Parameters\n")
		f.write("exclude             scaled1-4\n")
		f.write("1-4scaling          1.0\n")
		f.write("cutoff              16\n")
		f.write("switching           on\n")
		f.write("vdwForceSwitching   on\n")
		f.write("switchdist          15\n")
		f.write("pairlistdist        17\n")
		f.write("\n")
		f.write("# Integrator Parameters\n")
		f.write("rigidBonds          all  ;# needed for 2fs steps\n")
		f.write("nonbondedFreq       1\n")
		f.write("fullElectFrequency  2\n")
		f.write("stepspercycle       10\n")
		f.write("\n")
		f.write("# Constant Temperature Control\n")
		f.write("langevin            on    ;# do langevin dynamics\n")
		f.write("langevinDamping     1     ;# damping coefficient (gamma) of 1/ps\n")
		f.write("langevinTemp        $temperature\n")
		f.write("langevinHydrogen    off    ;# don't couple langevin bath to hydrogens\n")
		f.write("\n")
		f.write("restartfreq         " + str(self.RESTFREQ) + "\n")
		f.write("dcdfreq             " + str(self.DCDFREQ) + "\n")
		f.write("xstFreq             " + str(self.XSTFREQ) + "\n")
		f.write("outputEnergies      " + str(self.OUTEN) + "\n")
		f.write("outputPressure      " + str(self.OUTP) + "\n")
		f.write("\n")
		f.write("#############################################################\n")
		f.write("## EXTRA PARAMETERS                                        ##\n")
		f.write("#############################################################\n")
		f.write("\n")
		f.write("# Put here any custom parameters that are specific to\n")
		f.write("# this job (e.g., SMD, TclForces, etc...)\n")
		f.write("TMD             on\n")
		f.write("TMDk            " + str(self.TMDk) + "\n")
		f.write("TMDOutputFreq   " + str(self.TMDOutFreq) + "\n")
		f.write("TMDFile         " + self.TMDFile + "\n")
		f.write("TMDLastStep     " + str(self.TMDLastStep) + "\n")
		f.write("\n")
		f.write("#############################################################\n")
		f.write("## EXECUTION SCRIPT                                        ##\n")
		f.write("#############################################################\n")
		f.write("\n")
		f.write("# Minimization\n")
		f.write("\n")
		f.write("minimize	[expr $minimize]\n")
		f.write("reinitvels	$temperature\n")
		f.write("run [expr int($time*1000000/$timestep)]\n")

		f.close()
```
