## Supplemental GUI for "CryoFold: determining protein structures and ensembles from cryo-EM data": TaMDScreen.py

```
#!/usr/bin/env python

from kivy.uix.boxlayout import BoxLayout
from kivy.uix.gridlayout import GridLayout
from kivy.event import EventDispatcher
from kivy.lang import Builder
from kivy.properties import ObjectProperty, StringProperty
from kivy.uix.screenmanager import ScreenManager, Screen
from kivy.uix.label import Label
from kivy.uix.textinput import TextInput
from kivy.uix.popup import Popup
from LoadDialog import LoadDialog
from AlertDialog import AlertDialog
from TaMD import TaMD
#from Colvarsgui import ColvarsObj

import os


# Builder.load_file('TaMD_GUI/BackgroundColor.kv')

Builder.load_file('TaMD_GUI/TaMDScreenManager.kv')

Builder.load_file('TaMD_GUI/TaMD_Screen1.kv')
Builder.load_file('TaMD_GUI/TaMD_Screen2.kv')
#Builder.load_file('TaMD_GUI/Colvars_Screen1.kv')
#Builder.load_file('TaMD_GUI/Colvars_Screen2.kv')

class TaMDPart(Screen):
    pass


class GeneralScreen(Screen):

    def dismiss_popup(self):
        self._popup.dismiss()

    def show_load(self, fileinput):
        content = LoadDialog(load=self.load, cancel=self.dismiss_popup, fileinput=fileinput)
        self._popup = Popup(title="Load file", content=content,
                            size_hint=(0.9, 0.9))
        self._popup.open()

    def show_alert(self, message):
        content = AlertDialog(close=self.dismiss_popup, message=message)
        self._popup = Popup(title="Alert", content=content,
                            size_hint=(0.5, 0.2))
        self._popup.open()

    def load(self, path, filename, fileinput):
        with open(os.path.join(path, filename[0])) as stream:
            getattr(self, fileinput).text = stream.name
            setattr(self.manager.statedata, fileinput, stream.name)
        self.dismiss_popup()

    def close():
        self.dismiss_popup()


class TaMDScreen(GeneralScreen):

    def on_leave(self, *largs):
        setattr(self.manager.statedata, 'psf', self.psf.text)
        setattr(self.manager.statedata, 'pdb', self.pdb.text)
        setattr(self.manager.statedata, 'ref', self.ref.text)
        setattr(self.manager.statedata, 'tmdk', self.tmdk.text)
        setattr(self.manager.statedata, 'tmdoutfreq', self.tmdoutfreq.text)
        setattr(self.manager.statedata, 'tmdlast', self.tmdlast.text)

class Simulation_params(GeneralScreen):

    def save(self, jobName, NO, TEMP, TIME_IN_NS, MIN, RESTARTFREQ, DCDFREQ, XSTFREQ, OUTEN, OUTP, fName):
    	namd = TaMD()
    	namd.getSpecifics(jobName, NO, self.manager.statedata.psf, self.manager.statedata.pdb, TEMP, TIME_IN_NS, MIN, RESTARTFREQ, DCDFREQ, XSTFREQ, OUTEN, OUTP, self.manager.statedata.ref, self.manager.statedata.tmdk, self.manager.statedata.tmdoutfreq, self.manager.statedata.tmdlast, fName)
    	namd.writeNAMD()


#class Colvars(GeneralScreen):

#    def on_leave(self, *largs):
#        setattr(self.manager.statedata, 'atomsFile', self.atomsFile.text)
#        setattr(self.manager.statedata, 'refPosFile', self.refPosFile.text)


#class ColvarsConfig(GeneralScreen):

#    def colvarsSave(self, trajFreq, restFreq, centers, targNumSteps, forceCon, fName):
#        colObj = ColvarsObj()
#        colObj.getSpecs(trajFreq, restFreq, self.manager.statedata.atomsFile, self.manager.statedata.refPosFile, centers, targNumSteps, forceCon, fName)
#        colObj.writeColvars()


class TaMDState(EventDispatcher):
    psf = ''
    pdb = ''
    ref = ''
    tmdk = ''
    tmdoutfreq = ''
    tmdlast = ''
    #protParFile = ''
    #atomsFile = ''
    #refPosFile = ''

class StateManager(ScreenManager):
    statedata = ObjectProperty(TaMDState())
```
