## Supplemental GUI for "CryoFold: determining protein structures and ensembles from cryo-EM data": AlertDialog.py

```
from kivy.factory import Factory
from kivy.lang import Builder
from kivy.uix.boxlayout import BoxLayout
from kivy.uix.button import Button
from kivy.uix.label import Label
from kivy.properties import ObjectProperty, StringProperty

class AlertDialog(BoxLayout):
    close = ObjectProperty(None)
    message = StringProperty('')

root = Builder.load_file('./Utils/alertdialog.kv')
Factory.register('AlertDialog', cls=AlertDialog)
```
