## Supplemental GUI for "CryoFold: determining protein structures and ensembles from cryo-EM data": LoadDialog.py

```
from kivy.factory import Factory
from kivy.lang import Builder
from kivy.uix.boxlayout import BoxLayout
from kivy.uix.button import Button
from kivy.properties import ObjectProperty, StringProperty
from os import getcwd

class LoadDialog(BoxLayout):
    load = ObjectProperty(None)
    cancel = ObjectProperty(None)
    fileinput = StringProperty(None)

    def getPath(self):
        return getcwd()

root = Builder.load_file('./Utils/loaddialog.kv')
Factory.register('LoadDialog', cls=LoadDialog)
```
