## Supplement data for "CryoFold: determining protein structures and ensembles from cryo-EM data"

Tables 28

Figures 36

**Supplementary Methods**

**Description of MELD** Modeling Employing Limited Data (MELD) employs a Bayesian inference approach (eq. (1)) to incorporate empirical data into MD simulations[1,2]. The prior $p(\vec{x})$ comes from an atomistic force field (ff14SB sidechain, ff99SB backbone) and an implicit solvent model (Generalized born with neck correction, gb-neck2) [3,4]. The likelihood $p(\vec{D}|\vec{x})$ determines how well do the sampled conformations agree with known data. $p(\vec{D})$ refers to the likelihood of the data, which we take as a normalization term that can typically be ignored.

$$\overbrace{p(\vec{x}|\vec{D})}^{\text{posterior}} = \frac{p(\vec{D}|\vec{x})p(\vec{x})}{p(\vec{D})} \sim \overbrace{p(\vec{D}|\vec{x})}^{\text{likelihood}}\,\overbrace{p(\vec{x})}^{\text{prior}}\,. \tag{1}$$

The kind of data that MELD is designed to handle has one or more of the following features: sparsity, noise and ambiguity. Brute-force use of such data is deemed inadequate for complete structure determination. A typical MD simulation starts from accurate initial models derived with high-resolution structural data[5]. However, at low resolutions assessing the quality of experimental data is ambiguous, often resulting in the determination of incorrect models. MELD addresses the refinement of low-resolution data by enforcing only a fraction (f%) of this data at every step of the MD simulation. Although the fraction (f) is kept constant during the simulation, the feature of the data used at every step is determined on the flight. For each data point we calculate a penalty term based on flat-bottom harmonic restraints (see eq. 2) which serves as the way of evaluating the likelihood ($p(\vec{D}|\vec{x})$).

$$E(r_{ij}) = \begin{cases} \frac{1}{2}k(r_1 - r_2)(2r_{ij} - r_1 - r_2) & \text{if } r_{ij} < r_1 \\ \frac{1}{2}k(r_{ij} - r_2)^2 & \text{if } r_1 \leq r_{ij} < r_2 \\ 0 & \text{if } r_2 \leq r_{ij} < r_3 \\ \frac{1}{2}k(r_{ij} - r_3)^2 & \text{if } r_3 \leq r_{ij} < r_4 \\ \frac{1}{2}k(r_4 - r_3)(2r_{ij} - r_4 - r_3) & \text{if } r_4 \leq r_{ij}, \end{cases} \quad (2)$$

When these restraints are satisfied they do not contribute to the energy or forces (flat bottom region see eq. 2 and Fig. S16). When the restraints are not satisfied they add energy penalties and the resulting force biases to the system, guiding it to regions that satisfy a subset of the data, or conformational envelopes.

For every structure, the energy restraints are evaluated for all the data provided. These restraints are sorted according to their magnitude, and only those in the lowest f% are chosen to guide the simulation until the next step. Consequently, the forces and energies acting on the system are deterministic: given a structure, the force field terms (priors in eq. 1) are computed, and the biasing forces are added as the subset of restraints that yield the lowest energy for the structure being sampled[1]. A Temperature and Hamiltonian replica exchange molecular dynamics protocol (H,T-REMD) is employed to accelerate the sampling of these low-energy conformations. The Hamiltonian changes by effectively changing the strength of the force constant used for the restraints (product $w(\lambda)k$, where $k$ is the force constant of the restraint and $w(\lambda)$ is the weight assigned to a particular replica (see eq. 2). The parameter $\lambda$ is used to map replica numbers to a value between 0 (lowest replica) and 1 (highest replica).

At the lowest replica index we sample from room temperature (300K) and we enforce the flat-bottom harmonic restraints at their maximum ($w(\lambda)=1$). At the highest replica index, the temperature is at its maximum (450K) and $w(\lambda)=0$ (we sample freely over the energy landscape). We used 30 replicas in all cases, and scaled the Hamiltonian and temperature ladders following previous protocols[2].

The resulting MELD ensembles are processed by analyzing the lowest temperature replicas, which represent the flat region of the energy restraints (the energy will be the same as in the original force field). A standard 2D-RMSD clustering protocol for structural similarity[6] is used to identify the lowest free energy structures.

**Description of MAINMAST** MAINMAST (MAINchin Model trAcing from Spanning Tree) is a *de novo* modeling program that directly builds protein main-chain structures from an EM map of around 4-5 Å or better resolutions[7]. MAINMAST automatically recognized main-chain positions in a map as dense regions and does not use any known structures or structural fragments.The procedure of MAINMAST consists of mainly four steps (Fig. S16). In the first step, MAINMAST identifies local dense points (LDPs) in an EM map by mean shifting algorithm. The mean shifting algorithm is a non-parametric clustering algorithm that was originally developed for image processing. In the MAINMAST algorithm, the assumption is that a density observed in an EM map is the sum of Gaussian density functions that originate from atom positions and the local maxima of dense regions corresponds to the atomic positions of the protein structure. All grid points in the map were iteratively shifted by a gaussian kernel function and then merged to the clusters. The representative points in the clusters are called LDPs. In the second step, all LDPs are connected by constructing a minimum spanning tree (MST). MST is a graph structure that connect all nodes with the minimal total weight of edges by a three-graph structure (i.e. no-cycle graph). We used the Euclidean distance between the two nodes as the weight of the edge. It was found that the most edges in the MST covers the main-chain of the protein structure in EM map. In the third step, the initial tree structure (MST) is refined iteratively by tabu search algorithm[8]. The longest path in the MST usually contains some wrong connection and disconnections. Therefore, the longest path in the MST cannot complete whole main-chain structure. In order to refine the MST, MAINMAST performs a tabu search method. A tabu search attempts to explore a large search space by using a list of moves that were recently considered and then forbidden. In the final step, the longest path of the refined tree is aligned with the amino acid sequence of the target protein. This process assigns optimal C$\alpha$ positions of the target protein on the path and evaluates the fit of the amino acid sequence to the longest path in a tree. For example, amino acids with a large side-chain are mapped to a position on the path with a high density and large volume. All the steps are performed multiple times with various parameter combinations, and then over six thousand C$\alpha$ models were generated. The models are then ranked with the density-volume matching (threading) score. For flpp3 and TRPV1, MAINMAST generated 6,048 and 8,065 C$\alpha$ models, respectively.

**Description of Resolution exchange MDFF** Main computational steps involved in an ReMDFF refinement[9] is described as follows (Fig. S16): First, the reported map is smoothed employing

Gaussian blurs with the half width $\sigma_i$ uniformly spaced at 0.5 Å. As a result a set of at most 11 density maps with varied local resolution were generated. Second, the initial model obtained from the MELD simulation is docked in the EM density using e.g., Situs [10] or Chimera[11,12]. The map with the lowest resolution was chosen for this purpose. Third in order to prevent over-fitting to the density, chirality and cis-peptide restraints are generated from the initial model according to protocols defined in (Schreiner et al., 2011). Finally in accordance to the scheme shown in (Fig. S16), the resolution exchange program is invoked within NAMD, performing ReMDFF. The map-model coupling parameter is empirically determined and is generally set between 0.3-0.6. In the replica-exchange scheme, exchanges were attempted every 20,000 steps. Depending on the system, exchange rates of 30%–60% was achieved. Simulations were performed in implicit solvent environment using CHARMM36m protein force field.

### Supplementary Protocols

**CryoFold for Ubiquitin** The synthetic density for ubiquitin was constructed using phenix.maps with the diffraction data reported for PDB:1UBQ, truncated at 3 Å. A standard MELD-CPI run was performed with 30 replicas[1,13] starting from an extended state and simulated for 25 ns. 50 evenly spaced structures were collected from the highest and lowest temperature replica. Typically, at least 500 ns sampling is required to get native-like topologies for ubiquitin[13]. A faster approach relies on using MAINMAST and a synthetic density map for ubiquitin to generate a series of alpha carbon traces ranging in quality. A conformation with high RMSD to the native structure (since it is known) was chosen from the MELD ensemble (25th frame in this case). Targeted MD was performed for 10 ns with this conformation and the target-RMSD delivered from the C$\alpha$ tracing as restraints. This targeted MD run improved the ubiquitin structure dramatically.

At this point a second MELD run was set up. We calculated C$\alpha$ contacts below 6Å and separated by more than four residues from the model generated in the previous round. We enforced the contacts as flat bottom harmonic restraints at 60% accuracy and secondary structure predictions[14] as in our previous work [1,13],. Simulations were run for C$\alpha$ 100 ns with this data and secondary structure predictions (PSIPRED) enforced. From these trajectories we selected frames based on the best cross-correlation overlap, which were fed to the MDFF machinery. Cross-correlation improved with trajectory length and correlated well with RMSD (see Fig. S).

These structures were further improved with ReMDFF, leading to three new MELD simulations at different MDFF accuracies (3, 4 and 5 Å). The same MELD protocol as in step 2 (Fig. 3) was used. In all cases, running microsecond long simulations resulted in frames of accuracy better than 2.5 Å. Selection of good models can be achieved either on the basis of clustering or by selecting on the basis of higher cross-correlation. These structures are then reintroduced into ReMDFF for a last stage of refinement. Altogether, the combination of MELD and ReMDFF defines new secondary structure elements and orientations and packing them to high accuracy within the density map.

**CryoFold for flpp3** CryoFold was utilized to generate ensembles of flpp3 structures from maps of resolution varying between 1.8 –5 Å. The serial femtosecond crystallography data used for these computations and the high-resolution flpp3 structure determined using traditional methods used here for benchmarking CryoFold are published independently[15].

Starting from a high resolution map of 1.8 Å, a low resolution map of 5 Å was generated by truncating the diffraction data with phenix.maps. Thereafter, initial protein topology to be used in MELD as pairwise distance restraints was obtained by generating C$\alpha$ traces using MAINMAST for both the 1.8 Å and the 5 Å maps. At 5 Å the tracing of the C$\alpha$ by MAINMAST was significantly worse than the other cases, with atoms overlapping (Fig.3A,B). Hence, the 5Å resolution was used as representative of a low resolution (*data-poor*) refinement.

MAINMAST structures were used to construct a contact map between C$\alpha$ atoms, considering all contacts with distances beneath 8 Å and separated by at least 4 residues in the linear chain. These contacts were enforced at 80% accuracy inside MELD simulations – these simulations started from an extended chain as prepared by AMBER's tleap program. At any stage in which MELD structures were refined with MDFF, we used these model as both an initial point and to calculate a contact map. We selected distances below 6 Å from the contact map and enforced them at 60%. This guarantees that we sample structures that are close to the MDFF model, but with enough wiggle room for refinement.

Higher uncertainties in C$\alpha$ positions of the C$\alpha$ atoms in the 5 Å resolution maps results in a concomitant inaccuracy of the predicted distances between these atoms. From the mainmast structure we calculated a contact maps excluding residues closer than 4 residues in the sequence. We ran three protocols with different accuracy and curoff for defining contacts: (1) 8 Å (15% accuracy), (2) 9 Å (20% accuracy) or (3) 10 Å (25% accuracy). At the end of these simulations the best structures from the trajectories (at least 750 ns long) were selected via cross correlation with the 5 Å map. The protocol with 20% of contacts enforced at 9 Å performed the best and hence its structures were used for ReMDFF refinement. Structures from ReMDFF were then used for another stage of MELD refinement. Since we do not know the accuracy of the contact mapd on ReMDFF structures we agan ran 5 protocols using contacts between C$\alpha$ atoms (8 Å cutoff) with an accuracy of either 55, 65, 75, 85 or 95%. The cross correlation was used to select the best protocol to carry forward.

In previous work, we had known accuracies for the input data that went into MELD, and relied on clustering to determine structures. For this case, we do not know a priori the accuracy, but we can always back calculate agreement with the density map which gives us a way to identify good structures – which are further refined wiht MDFF. Protocols that enforce more data that actually present in the native structure will add a biasing potential in the native state, and thus it cannot be guaranteed that this would still be the most populated state in MELD simulations.

**CryoFold for TRPV1** TRPV1 presents a unique case to test capabilities of CryoFold on systems with heterogeneous CryoEM densities. Concentrating on only the soluble domain of TRPV1, the protein was truncated at its membrane interface, while also fixing the interfacial residues with cartesian restrains. This approach enables the application of our methodology to large macromolecules by fragmenting it into several independent smaller domains. Of particular interest in the soluble domain are $\beta$-sheets and a loop (residues 112-122 and 184 to 224) connecting the sheets to the transmembrane domain. These structural elements were heated at 450 K for 10 ns converting them to an unstructured polypeptide chain. Beginning the CryoFold refinement, in the MELD setup 40% of the PSIPRED-predicted secondary structures were enforced for these two regions. A list of all possible contacts between the aforementioned region of interest and additional packing areas within a radius of 8Å (residues 19 to 52 and 81 to 94) was created. At each MELD timestep at most 10 of the total 1610 possible contacts were enforced. This folding procedure resulted in an ensemble of conformations, where the $\beta$-sheets were packed close to the desired density, while the loop amino acids sample non-native regions. In order to filter a structure from the ensemble, cross-correlation to the map was calculated. Structure which had the highest cross-correlation serves as the most representative structure i.e, closet to the density among the MELD ensemble. Subsequently, ReMDFF refinement was initiated from this representative structure using 11 maps of resolution blurred incrementally by steps of 0.5 Å Gaussian half-width. ReMDFF was able to improve on the loop conformation and the structure with the best cross-correlation was fed in the next round on MELD simulations. In a subsequent round, the contacts present in ReMDFF's structure affecting the region 189-224 were identified resulting in 39 contacts. MELD was asked to satisfy 10, 20 or 30 contacts in three different simulations (each with 30 replicas). Cross-correlation score of the ensemble of MELD structures was used to determine the best structures to feed into a final round of ReMDFF refinement.

**CryoFold for Apoferritin** This was a time-senstive target using electron density maps at different resolution provided by the EMDataResource. We used MAINMAST to generate 100 C$\alpha$ models and reconstructed to full atom representation with PULCHRA. We used agreement with the density map to narrow these 100 full atomic models to 30. We minimized those 30 structures with

AMBER[16] and proceeded to collect contacts for each model as $C\alpha$-$C\alpha$ distances below 8Å – only contacts that were more than 6 residues apart in sequence were kept. Rather than using a consensus approach, we input each set of restraints corresponding to an initial MAINMAST prediction, with the constraint that at any point only 80% of the data needs to be satisfied – and only one block of restraints (the one most compatible with the sampled structure) needs to be satisfied during the simulation. Each set had around 100 possible contacts to satisfy. The minimized structures were used to seed the MELD replica exchange ladder and five simulations were run for each resolution, the longest trajectories were 750ns. The best models according to the cross correlation were selected for further refinement with MDFF.

**CryoFold for ATP synthase** For the large ATP synthase assembly we were interested in observing relative motion between domains starting from PDB 6RET. We fixed the arm of the synthase (residues 1 to 2477) with cartesian restraints and set harmonic restraints within each of the other individual domains as contacts (different chains) to keep each domain from unfolding in the replica exchange ladder based on initial structure– and asked that at least 60% be satisfied at low replica indexes(see SI fig ). Finally, the central stalk (yellow in SI fig. ) was free to move. Secondary structure was imposed at 80% accuracy through out the MELD trajectories. We then analyzed the ability of the ensembles to sample other biologically relevant conformations ot the synthase.

**CryoFold for CorA** Pentameric Magnesium channel CorA serves as another transmembrane protein (resolved at 3.8 Å) system that was attempted for modeling. Initial position of the $C\alpha$ atoms were generated using MAINMAST. Thereafter, targeted MD simulation was initiated to fit a linear polypeptide to the MAINMAST-generated C$\alpha$ positions. The structure obtained served as a template for MELD simulations. Given the high confidence on the C$\alpha$ positions for the amino acids and the lack of a good membrane implicit solvent, cartesian restraints were used on all C$\alpha$ positions. Secondary structure prediction was imposed at 80% since membrane protein structure predictions were more accurate than the globular ones. MELD structures recovered a large fraction of secondary structure but introduced small kinks and discontinuities in some places of the transmembrane helix. Structure with high cross-correlation to the cryoEM densities were chosen for ReMDFF refinment. ReMDFF was initiated using 11 densities with highest resolution density to be the experimentally determined one at 3.8 Å, the resolution blurred by Gaussian smoothing in steps of 0.5 Å. Thereafter, the structure with the highest cross-correlation to the map with highest resolution served as the template for another round of MELD simulations. One more round of ReMDFF refinement was initiated from the best structure with maximum cross correlation.

### ROSETTA comparison

**Rosetta *de novo* modeling for flpp3** The flpp3 secondary structure was predicted with Jufo9d[17] and PSIPRED[18,19] servers. Nine- and three-mer fragments were generated using weighted quota protocols such that the PSIPRED and Jufo9d secondary structure predictions contributed to the fragment secondary structures equally. To eliminate bias for benchmarking purposes, the PDB 2MU4, corresponding to the structure of flpp3[20] was omitted during the fragment generation stage. The DUF3568 Family Protein from *Francisella tularensis* virulence determinant sequence (Uniprot: Q5NF33) was used as input for the fragment generation. As per standard prediction protocols, 1000 peptide candidates were considered with the top scoring 200 used as the final fragments for the *de novo* fold.

After generating fragments, *de novo* structure prediction was employed using the automated Rosetta protocol resolution-adapted recombination of structural features (RASREC)[21]. Extensive RASREC details are available in the original publication, however control of model generation can be altered in several ways. For this study, RASREC generated decoys were calculated using default flags, which automates decoy selection through four stages of centroid model generation and two stages of full atom decoy relaxation until convergence is achieved. RASREC has been successfully used in previous studies using sparse NMR restraints and co-evolved position contacts as restraints to generate *de novo* models; however, structural restraints nor evolutionary information were not employed for this study and the *de novo* folds relied solely on fragment inputs.

The RASREC algorithm is written to accept or discard until the pool size is full of decoys that were selected by the work-load manager. The end of the run generates a final pool of decoys, which contain as many decoys as specified by the pool size flag. With flpp3, this pool size was set to 500 decoys, which is the default setting. We clustered these 500 decoys using Calibur with default settings and then analyzed the RMSD of these clusters in PyMol by aligning the C$\alpha$ atoms of $\beta$-sheets (residues 6-8, 11-18, 47-51, 55-61, 69-77, 93-112) and $\alpha$-helices (residues 20-33, 93-112). The flpp3 de novo folding prediction was carried out with only fragments for inputs and achieved converged folds that are similar the NMR determined structure (Fig S1). Relying solely on fragments derived from secondary structure predictions Rosetta predicted flpp3 to an RMSD of

2.3 Å 3.3 Å and 1.8 Å of the 1st, 2nd, and 3rd largest cluster representatives, which was the lowest scoring decoy from each cluster. In Fig S1A, 2MU4 is shown for state 1 of the NMR ensemble next to the superposed decoys from decoys from the clusters. The $\alpha$-helices do not appear to reach convergence, and RASREC predicted a $\beta$-sheet after the first $\alpha$-helix (arrow #1). Fig S1B, shows that the $\beta$-sheets were predicted very well; however, the $\beta$-sheet N-terminal fold is missing in the RASREC predicted decoys (arrow #2).

**Rosetta-ES *de novo* loop modeling for TRPV1** Rosetta ver. 3.9 (rosetta_bin_linux_2018.33.60351 _bundle) was used. We followed the tutorial released on the tutorial website [22]. All the parameters used were as described in the tutorial. First, nine and three residue fragment structures for the query protein were generated by *grower_prep* in the Rosetta package. Then, the *RunRosettaES.py* script [23] built each segment from the generated fragments. The rebuilt regions were assembled by a Monte Carlo Assembly algorithm in the RunRosettaES.py. After the assembly step, the total 100 models were generated and ranked by Rosetta Energy. It should be noted that all generated models have steric clashes. Therefore, we could not have performed further Rosetta refinement.

**Rosetta-EM *de novo* modeling for flpp3 and CorA** First, nine residue fragment structures for the query protein were generated on the Robetta website (http://robetta.bakerlab.org/fragmentqueue.jsp). We excluded fragments from homologues proteins. We performed a local fragment search in an input EM map by using *denovo_density*. This procedure searches the density map for each sequence-predicted backbone fragment generated in the previous step. In the denovo_density command, the number of translations to search (option: -n_to_search) was set to two times the number of residues.

The number of intermediate solutions to keep (option: -n_filtered) was set to ten times the number of residues. The placed fragments were assembled by Monte Carlo sampling, then the consensus was assigned from the Monte Carlo trajectories. The final output file of the consensus assignment was used as input to RosettaCM. RosettaCM was applied to fill gaps where the fragments were not assigned by *denovo_density* to complete a model and to refine the overall model structure. A total of 1,100 full-atom models were generated by RosettaCM. All of the 1,100 full-atom models were ranked by the total score (Rosetta Energy + density score). We selected the best 10% of the ranked models, and then selected the best model based on the density score.

**Graphical User Interface**

**Requirements**

MAINMAST requires 40-200 CPU hrs for the full-automated computation. This time could be reduced if the user manually checks the models of backbone trace by eye.

MELD requires 30 dedicated GPUs and a couple of days depending on the system size. MELD currently performs synchronous REMD, so each GPU is a different replica.

ReMDFF requires 1-2 days on 11 CPUs, depending on system size.

Note that the CryoFold GUI currently supports MDFF, not ReMDFF. MDFF requires less CPU-hours than ReMDFF.

**Installation**

The CryoFold GUI is optimized for LINUX/macOS distributions.

Anaconda and GFortran needs to be installed on your workstation in order to run the CryoFold GUI. CryoFold has four sections, namely, MAINMAST, TMD, MDFF and MELD. The GUI submits MAINMAST and MELD jobs, which requires these softwares to be installed as well. For TMD and MDFF, CryoFold GUI generates NAMD input scripts that the user can submit manually (NAMD need not be installed to run the GUI). However, these two sections require VMD to be installed.

- Anaconda installation: https://www.anaconda.com/distribution/
- GFortran installation: https://gcc.gnu.org/wiki/GFortran
- VMD installation: https://www.ks.uiuc.edu/Research/vmd/
- NAMD installation (not required to run GUI): https://www.ks.uiuc.edu/Research/namd/
- OpenMM installation: https://github.com/pandegroup/openmm
- MELD installation: https://github.com/maccallumlab/meld

**Linux installation:** After downloading and unzipping the folder, open a terminal and go to the `CryoFoldGUI` folder. Run `install_linux.sh` script to install everything needed - GFortran compilation of Mainmast and ThreadCA, Anaconda environment creation, python packages (kivy, numpy, mdtraj and meld). Finally, run `start.sh` to launch the CryoFold GUI.

**MacOSX installation:** After downloading and unzipping the folder, open a terminal and go to the `CryoFoldGUI` folder. Run `install_mac.sh` script to install everything needed - GFortran compilation of Mainmast and ThreadCA, Anaconda environment creation, python packages (kivy, numpy, mdtraj and meld). Finally, run `start.sh` to launch the CryoFold GUI.

**Manual installation:** Alternatively, the packages could be installed manually with the instructions on the following page:

- Numpy: https://anaconda.org/anaconda/numpy
- Mdtraj: https://anaconda.org/omnia/mdtraj
- MELD: https://github.com/maccallumlab/meld
- Kivy: https://anaconda.org/conda-forge/kivy

For gfortran compilation of MAINMAST and ThreadCA, run the following commands from the command line:

```
gfortran ./MAINMAST_GUI/MainmastThreadCA/MAINMAST.f -w -O3 -fbounds-check
-o ./MAINMAST_GUI/MainmastThreadCA/MAINMAST -mcmodel=medium
```

```
gfortran ./MAINMAST_GUI/MainmastThreadCA/ThreadCA.f -w -O3 -fbounds-check
-o ./MAINMAST_GUI/MainmastThreadCA/ThreadCA -mcmodel=medium
```

The CryoFold GUI can then be launched by running `gui.py` by typing ”python gui.py” or by creating an executable.

**Notes:** Manual changes to installation script (`install_linux.sh` or `install_mac.sh`) might be necessary. Line 10 of the script specifies python version 3.5, which the user can change as per their choice. Additionally, MELD is available for CUDA versions 7.5, 8.0, 9.0 and 9.2. The current script will install meld-cuda75 (line 10), which is for CUDA 7.5. The user can manually change this (to meld-cuda80, meld-cuda90 or meld-cuda92) to suit their CUDA compiler.

We recommend using the installation script as it creates a separate python environment called ”CryoFold” where all the packages are installed in. This keeps the default packages untouched.

For more information, refer to the ”README.txt” file distributed with the GUI.

**Usage** The CryoFold GUI has 4 sections, namely, MAINMAST, Targeted Molecular Dynamics, MDFF, and MELD. Note that depending upon the map resolution, you might not need to use all 4 sections. Given below is an explanation of all the parameters that are used as input in CryoFold GUI.

**MAINMAST** :

MAINMAST protocol consists of mainly four steps: (1) Identify local dense points in an

EM map by Mean Shifting clustering algorithm; (2) Connect all Local Dense Points (LDPs) by Minimum Spanning Tree; (3) Refine Tree structure by Tabu Search algorithm; (4) Thread sequence on the longest path. Program MAINMAST will do the (1)-(3) steps. Program ThreadCA threads the amino acid sequence on the longest path in the final step. Here, we explain details of input files and paramaters. Basically, user does not have to change the default values.

- Density file: MAINMAST requires SITUS format file as an input EM map file. MRC format file can be converted to SITUS format by map2map program in SITUS package.
- Bandwidth of the gaussian filter: This paramater control the side of the gaussian filter in Mean Shifting step. Default value is 2.0.
- Threshold of density value: To remove noisy data, user can specify the threshold value. The optimal value depends on the EM map. We recommend (0.25 or 0.5) x Author recommended contour level which is provided by EMDB.
- Filter of the representative point: After Mean Shifting, low dense points are removed by this threshold value. The default value is 0.1.
- Number of iterations: It control the number of iteration in tabu search step. Large EM map needs large number. The default value is 5000. We recommend 500-5000.
- Size of tabu-list: Number of forbidden recent steps in tabu-search. The default value is 100.
- Constraint of total length of edge: MAINMAST considers only the tree graphs whose total length of edge are below [float]x(Total length of MST). The default value is 1.01.

- Keep edge where distance: MAINMAST keeps the edges whose length are shorter this parameter. The default is 1.0.
- Max shift distance d: The parameter for Mean-shifting. During the Mean-shifting, LDPs are not moved than the distance d. The default value is 10.0.
- After MeanShifting, merge d: If the distance between two LDPs is closer than d, these LDPs are merged.
- Number of Neighbors: parameter for tabu-search. It control the size of search space for local.
- Radius of Local MST: It control the number of edges.The default value is 10.0.
- Reverse mode, reverse mainchain order: In the threading part, opposite order of main-chain structure is used.
- parameter file: The parameter file for the threading step. 20AA.param.
- Result of SPIDER2: Result file of Secondary structure prediction.

  This is demonstrated in movie MAINMAST_movie.mov.

**Targeted Molecular Dynamics (TMD)** :

During TMD a random coil structure with the sequence of the target protein is fitted to the backbone trace derived from MAINMAST. This step requires a PDB file, a PSF file, and a reference file. Other parameters can be left at default values.

- PSF file: Protein structure file. This file contains molecular topology information. To generate this file, you need to use the AutoPSF package of VMD. You need the PDB file (see below) and the protein topology file to generate this it.
- PDB file: The protein PDB file.
- TMD reference file: PDB file where occupancy column is nonzero only for atoms to be used in TMD (in this case, the C$\alpha$ atoms fitted by MAINMAST).
- TMD Force constant: This quantity denotes the strength of the force acting on the targeted atoms. Default: 200 kcal/mol
- TMD output frequency: TMD output is saved to disk after every ”n” steps of targeted MD simulation where ”n” is the number entered here. Default: 5000
- TMD Last step: Number of TMD steps. Default: 50000
- Job Name: Name of the job. This is user’s choice. User is strongly recommended to not put spaces or special characters in the job name.
- Temperature (K): Temperature at which Targeted MD simulations are performed. Typically it would be 310 K or something close. (Default: 310 K)
- Minimization steps: Number of steps of gradient-descent minimization of the PDB before initiating targeted MD (Default: 1000).

- DCD frequency: Frames are saved to disk after every ”n” steps of targeted MD simulation where ”n” is the number entered here. Lowering this value will take up more disk space to store the targeted MD trajectory (Default: 5000).

- Energy output frequency: Energy of the system is stored to the log file every ”n” steps where ”n” is the number entered here. Also see ”Pressure output frequency” below (Default: 5000).

- File name: Name of the NAMD input script. This is user’s choice. User is strongly recommended to not put spaces or special characters in the File name.

- Job number: Ignore unless restarting from a previous Targeted MD simulation (job number is set to 0 when ignored). If restarting from job number ’n’, set job number as ’n+1’ (first restart will have job number = 1, next restart 2, etc.)

- Time (ns): Duration for which targeted MD simulation is performed. Timestep used is 2 fs. This means, if user wants to run 1 ns targeted MD, simulation will be performed for 500000 steps, not including the minimization steps described above (Default: 5 ns)).

- Restart frequency: Restart files are saved every ”n” steps where ”n” is the number entered here. Restart files come in handy if the job crashes for some reason. Keeping this value too low (high restart frequency) might make the job slow (Default: 5000).

- XST frequency: Periodic box information are saved every ”n” steps where ”n” is the number entered here. See ”Restart frequency” above for purpose and recommended usage (Default: 5000).

- Pressure output frequency: Pressure of the system is stored to the log file every ”n” steps where ”n” is the number entered here. Also see ”Energy output frequency” above (Default: 5000).

  This is demonstrated in movie TMD_movie.mov. The TMD-driven folding is shown in movie TMD_beforeAndAfter.mov.

**MDFF** :

MDFF refines an initial search model (either from MELD or from MAINMAST+TMD) against the electron density using flexible fitting. This step requires a PDB file, a PSF file, and a map file. Other parameters can be left at default values.

- PSF file: See ”PSF” in Targeted molecular dynamics.
- PDB file: See ”PDB” in Targeted molecular dynamics.
- Map file: EM map. Could be .mrc or .ccp4 formatted.
- GSCALE: Strength of coupling between EM map and MD simulation. Denotes how strongly the map influences the simulation. Higher number means stronger influence (Default: 0.3).
- NUMSTEPS: Number of MDFF steps (Default: 50000).

  This is demonstrated in movie MDFF_movie.mov.

**MELD** : Here, the density-fitted model is refined to improve secondary structure content.

- Substructure Prediction File: Output from PSIPRED
- MDFF-MAINMAST file: This is the output (PDB) from MDFF-MAINMAST
- Sequence: FASTA without header (not needed if starting from a PDB)
- Fraction of secondary structure to trust: User's choice. Recommended value is 70% for globular and 90% for membrane proteins
- Fraction of initial pdb contacts to trust: User's choice. See paper for recommended values (Default: 40%)
- Maximum distance: Only distances closer than this value are kept. Note that the units are nm. Default: 0.8 nm.
- Number of steps: Number of MELD steps (Default: 10000 steps)
- Lowest Temperature: Temperature of the lowest replica (Default: 300 K).
- Number of replicas: Default: 30
- Block size: Restart files are saved after every block (higher block size means restart files saved less frequently). Default: 100 steps (5ns)
- Highest Temperature: Temperature of the highest replica (Default: 450 K).

This is demonstrated in movie MELD_movie.mov. The MELD-driven folding is shown in movie MELD_beforeAndAfter.mov.

Finally, the MELD-refined structure is plugged back into MDFF, and the MDFF-MELD iteration continues till convergence.

### Tables

Table S1: Ubiquitin at 3.0 Å resolution.

| Molprobity Parameters | MELD-ReMDFF | Original (1UBQ) |
|---|---|---|
| Poor rotamers | 3.33 | 8.82 |
| Favoured rotamers | 93.33 | 80.88 |
| Ramachandran outliers | 1.41 | 0.0 |
| Ramchandran favoured | 95.77 | 100 |
| MolProbity score (percentile) | 1.19(99) | 2.26(61) |
| $C_{\beta}$ deviations | 1.43 | 1.43 |
| Bad bonds | 0.33 | 0.0 |
| Bad angles | 0.12 | 1.96 |
| Cis Prolines | 0.0 | 0.0 |
| Clash score (percentile) | 0 | 10.56 |
| EM Ringer | 0.57 | 1.72 |

Table S2: Flippase at 1.8 Å resolution

| Molprobity Parameters | CryoFold | Rosetta | Original (unpublished SFX model) |
|---|---|---|---|
| Poor rotamers | 0 | 0.0 | 0.0 |
| Favoured rotamers | 93.90 | 97.85 | 80 |
| Ramachandran outliers | 0.0 | 1.89 | 0 |
| Ramchandran favoured | 93.40 | 96.23 | 100.00 |
| MolProbity score (percentile) | 0.93 (100) | 1.66(91) | 0.85(100) |
| $C_\beta$ deviations | 3.96 | 0.00 | 0.00 |
| Bad bonds | 0.00 | 0.59 | 0.0 |
| Bad angles | 0.26 | 0.87 | 0.00 |
| Cis Prolines | 0.00 | 0.0 | 0.00 |
| Clash score (percentile) | 0 | 7.29(85) | 1.26(99) |
| EM Ringer | 2.17 | 2.11 | 1.49 |

Table S3: Flippase at 5.0 Å resolution

| Molprobity Parameters | CryoFold | Rosetta | Original (unpublished SFX model) |
|---|---|---|---|
| Poor rotamers | 1.22 | 0.0 | 0.0 |
| Favoured rotamers | 98.78 | 97.85 | 80 |
| Ramachandran outliers | 0.0 | 1.89 | 0 |
| Ramchandran favoured | 94.34 | 96.23 | 100.00 |
| MolProbity score (percentile) | 0.95 (100) | 1.66(91) | 0.85(100) |
| $C_\beta$ deviations | 0.00 | 0.00 | 0.00 |
| Bad bonds | 0.00 | 0.59 | 0.0 |
| Bad angles | 0.09 | 0.87 | 0.00 |
| Cis Prolines | 0.00 | 0.0 | 0.00 |
| Clash score (percentile) | 0 | 7.29(85) | 1.26(99) |
| EM Ringer | 0.579 | 0.787 | 3.89 |

Table S4: TRPV1 soluble domain

| Molprobity Parameters | CryoFold | Rosetta | Original (5IRZ) |
|---|---|---|---|
| Poor rotamers | 4.88 | 0.0 | 0.00 |
| Favoured rotamers | 92.37 | 0.00 | 98.90 |
| Ramachandran outliers | 1.56 | 11.16 | 0.00 |
| Ramchandran favoured | 93.75 | 73.02 | 92.54 |
| MolProbity score (percentile) | 1.67 (90) | 3.56(8) | 1.99 (100) |
| $C_\beta$ deviations | 10.00 | 0.00 | 0.00 |
| Bad bonds | 0.22 | 0.09 | 0.07 |
| Bad angles | 0.28 | 3.66 | 0.00 |
| Cis Prolines | 0.00 | 0.0 | 10.00 |
| Clash score (percentile) | 0 | 7.29(85) | 10.14 |
| EM Ringer | 3.35 | - | 3.26 |

Table S5: TRPV1 deposited models

| Model | Rama Favoured | Rotamer outliers | Rama outliers |
|---|---|---|---|
| Model 1 | 81.20 | 0.37 | 0.16 |
| Model 2 | 93.06 | 4.17 | 0.16 |
| Model 3 | 91.96 | 0.23 | 1.65 |
| Model 4(new, old) | 93.75, 92.30 | 2.75, 3.47 | 1.56, 3.37 |
| Model 5 | 90.94 | 1.62 | 0.17 |
| Model 6 | 89.94 | 0.23 | 0.00 |
| Model 7 | 94.28 | 26.57 | 0.00 |
| Model 8 | 49.84 | 0.00 | 20.13 |
| Model 9 | 53.67 | 0.69 | 18.53 |
| Model 10 | 53.99 | 0.35 | 20.13 |
| Model 11 | 57.51 | 0.69 | 16.61 |
| Model 12 | 55.27 | 0.35 | 13.74 |
| Model 13 | 58.15 | 1.39 | 15.97 |
| Model 14 | 48.56 | 0.00 | 22.68 |
| Model 15 | 53.67 | 0.35 | 20.77 |
| Model 16 | 52.40 | 0.35 | 18.85 |
| Model 17 | 91.51 | 0.20 | 0.35 |
| Model 18 | 49.20 | 0.69 | 23.32 |

Table S6: Rotary substates of mitochondrial $F_1$ - $F_0$ ATPsynthase in *Polytomella* sp.

| State | PDB | Resolution (Å) | Frames | CC | EM Ringer |
|---|---|---|---|---|---|
| I | 6RET | 4.3 | PDB | 0.82 | 1.88 |
| | | | MELD | 0.82 | 1.88 |
| | | | MELD-MDFF | 0.82 | 1.82 |
| II | 6RDQ | 4.0 | PDB | 0.87 | 2.94 |
| | | | MELD | 0.48 | 1.63 |
| | | | MELD-MDFF | 0.82 | 2.82 |
| III | 6RDW | 3.8 | PDB | 0.81 | 3.79 |
| | | | MELD | 0.34 | 1.74 |
| | | | MELD-MDFF | 0.74 | 3.61 |
| IV | 6RDK | 3.7 | PDB | 0.84 | 3.20 |
| | | | MELD | 0.4 | 1.81 |
| | | | MELD-MDFF | 0.8 | 2.96 |

Table S7: Structure based model quality for rotary substates of mitochondrial $F_1$ - $F_0$ ATPsynthase in *Polytomella* sp.

**A**

| Molprobity Parameters | 6RET | MELD (I) | MELD-MDFF (I) |
|---|---|---|---|
| Clashscore | 7.7 | 0 | 0 |
| Poor rotamers (%) | 0.41 | 0.42 | 0.50 |
| Favored rotamers (%) | 91.64 | 91.82 | 92.07 |
| Ramachandran outliers (%) | 0.14 | 0.14 | 0.13 |
| Ramachandran favored (%) | 93.48 | 93.48 | 93.88 |
| MolProbity | 1.85 | 0.93 | 0.91 |
| $C\beta$ deviations (%) | 0.0 | 0.0 | 0.02 |
| Bad bonds (%) | 0.0 | 0.09 | 0.06 |
| Bad angles (%) | 0.03 | 0.10 | 0.08 |
| Cis Prolines (%) | 2.12 | 2.12 | 2.12 |

**B**

| Molprobity Parameters | 6RDQ | MELD (II) | MELD-MDFF (II) |
|---|---|---|---|
| Clashscore | 6.31 | 0.0 | 0.0 |
| Poor rotamers (%) | 0.49 | 4.11 | 3.70 |
| Favored rotamers (%) | 90.20 | 87.13 | 89.45 |
| Ramachandran outliers (%) | 0.21 | 3.10 | 2.06 |
| Ramachandran favored (%) | 94.27 | 90.05 | 92.18 |
| MolProbity | 1.73 | 1.51 | 1.41 |
| $C\beta$ deviations (%) | 0.0 | 5.71 | 5.44 |
| Bad bonds (%) | 0.0 | 3.43 | 3.55 |
| Bad angles (%) | 0.0 | 3.65 | 3.81 |
| Cis Prolines (%) | 2.12 | 2.12 | 2.12 |

**C**

| Molprobity Parameters | 6RDW | MELD (III) | MELD-MDFF (III) |
|---|---|---|---|
| Clashscore | 6.32 | 0.0 | 0.0 |
| Poor rotamers (%) | 0.68 | 4.05 | 3.80 |
| Favored rotamers (%) | 88.19 | 88.21 | 88.08 |
| Ramachandran outliers (%) | 0.24 | 2.60 | 2.50 |
| Ramachandran favored (%) | 93.54 | 90.68 | 91.28 |
| MolProbity | 1.77 | 1.49 | 1.45 |
| $C\beta$ deviations (%) | 0.0 | 5.71 | 6.45 |
| Bad bonds (%) | 0.01 | 3.24 | 3.57 |
| Bad angles (%) | 0.04 | 3.76 | 3.96 |
| Cis Prolines (%) | 2.13 | 2.12 | 2.12 |

**D**

| Molprobity Parameters | 6RDK | MELD (IV) | MELD-MDFF (IV) |
|---|---|---|---|
| Clashscore | 5.69 | 0.0 | 0.0 |
| Poor rotamers (%) | 0.47 | 4.21 | 3.67 |
| Favored rotamers (%) | 92.20 | 87.32 | 89.18 |
| Ramachandran outliers (%) | 0.24 | 3.13 | 2.14 |
| Ramachandran favored (%) | 94.48 | 90.01 | 92.12 |
| MolProbity | 1.69 | 1.52 | 1.41 |
| $C\beta$ deviations (%) | 0.0 | 5.90 | 6.10 |
| Bad bonds (%) | 0.0 | 3.42 | 3.54 |
| Bad angles (%) | 0.0 | 3.91 | 3.87 |
| Cis Prolines (%) | 2.12 | 2.12 | 2.12 |

Table S8: Mg $^{2+}$ Channel CorA at 3.8 Å resolution

| MolprobityParameters | CryoFold | Rosetta | Original (3JCF) |
|---|---|---|---|
| Poor rotamers | 1.07 | 0.0 | 0.0 |
| Favoured rotamers | 94.22 | 98.78 | 98.73 |
| Ramachandran outliers | 1.52 | 1.15 | 0 |
| Ramchandran favoured | 93.62 | 96.54 | 95.39 |
| MolProbity score (percentile) | 0.94 (100) | 1.50(95) | 1.75(87) |
| $C_\beta$ deviations | 1.49 | 0.30 | 0.00 |
| Bad bonds | 0.43 | 0.44 | 0.0 |
| Bad angles | 0.18 | 0.62 | 0.00 |
| Cis Prolines | 0.0 | 0.0 | 0.0 |
| Clash score | 0.0 | 5.11(93) | 7.92(82) |
| EM Ringer | 1.14 | 1.76 | 2.29 |

**Figures**

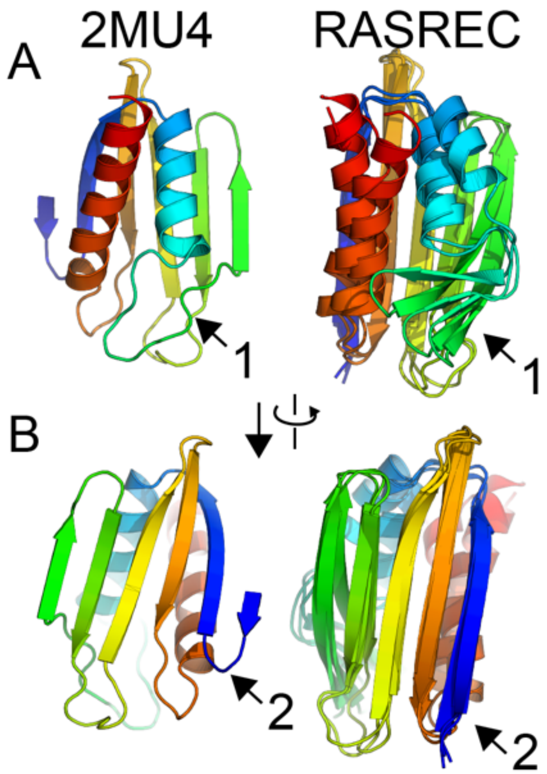

Figure S1: **De novo folding of flpp3 using RASREC Rosetta converged on folds that resemble the structure determined by SFX.** A) The structure of flpp3 shown with rainbow coloring from the N-terminus (blue) to the C-terminus (red) and the superposed output from the top three clusters of the RASREC predictions. The data output shows that RASREC predicted the helices with relatively low convergence. Additionally, RASREC predicted $\beta$-sheets that were not observed in the SFX structure. B) A 90° rotation of the model illustrates that the $\beta$-sheets were predicted relatively well, but the N-terminal sheet was overestimated.

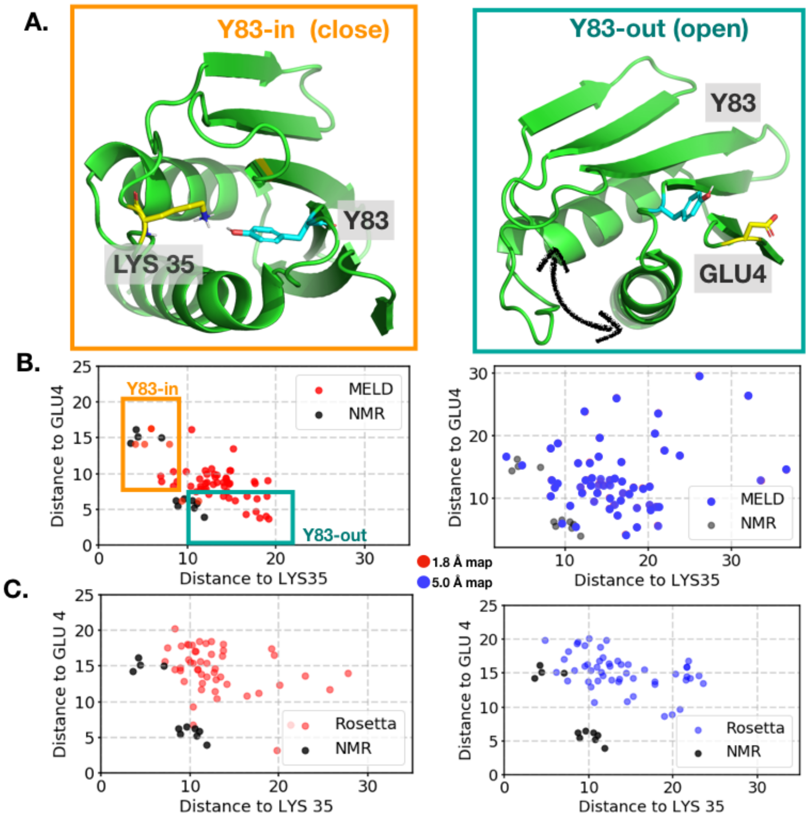

Figure S2: **Flpp3 ensembles from MELD overlapped with the NMR derived ensembles** A) The two predominant states of flpp3 shown in cartoon representation in green. The state Y83-in (left panel) is characterized by the residue Y83 (depicted in cyan stick representation) being buried in the protein proximal to the residue K35 (shown in yellow stick representation). Conversely, the state Y83-out (right panel) is characterized by the residue Y83 being solvent exposed and closer to E4 (depicted in yellow stick representation). B) The ensembles elucidated by MELD sampling (red dots [representing 1.8 Å resolution] and blue dots [5 Å resolution]) and NMR (black dots) projected onto a 2D space spanned by the distance between Y83 and E4 and K35. The ensembles sampled in MELD and NMR overlap for both the Y83-in states (orange rectangle) as well as the Y83-out state (blue rectangle). C) Scatter plot depicting ensembles sampled in by Rosetta and NMR projected onto the distance between Y83-E4 and Y83-K35.

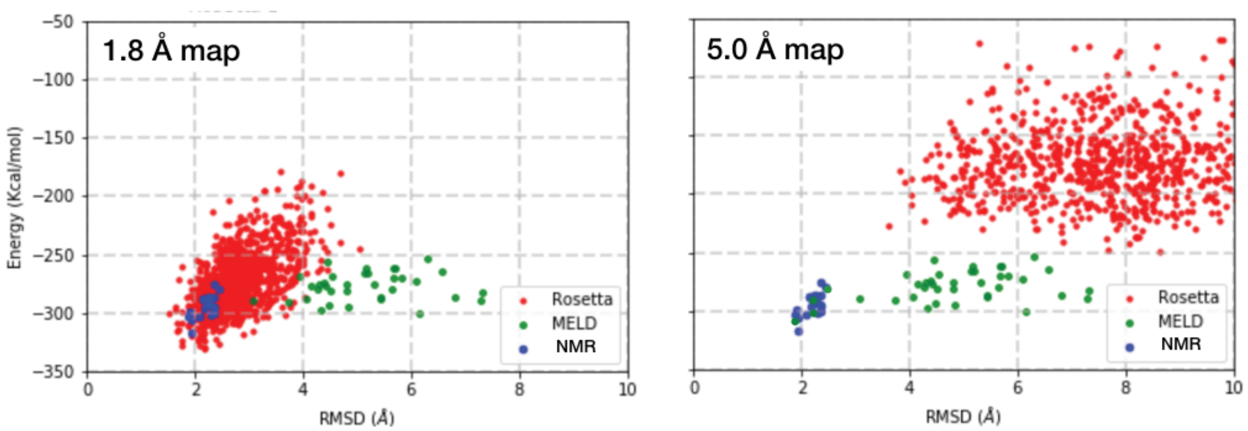

Figure S3: **Model quality of MELD and Rosetta modes for flpp3.** Rosetta relaxation was performed on 50 MELD models (green) and 1100 Rosetta models (red) and 20 structures of Flpp3, PDB 2MU4 (blue). Along the Rosetta energy (Y-axis), MELD models have a same distribution with NMR structures of flpp3. In the structural sampling space (X-axis), MELD models cover the similar RMSD range with Rosetta models.

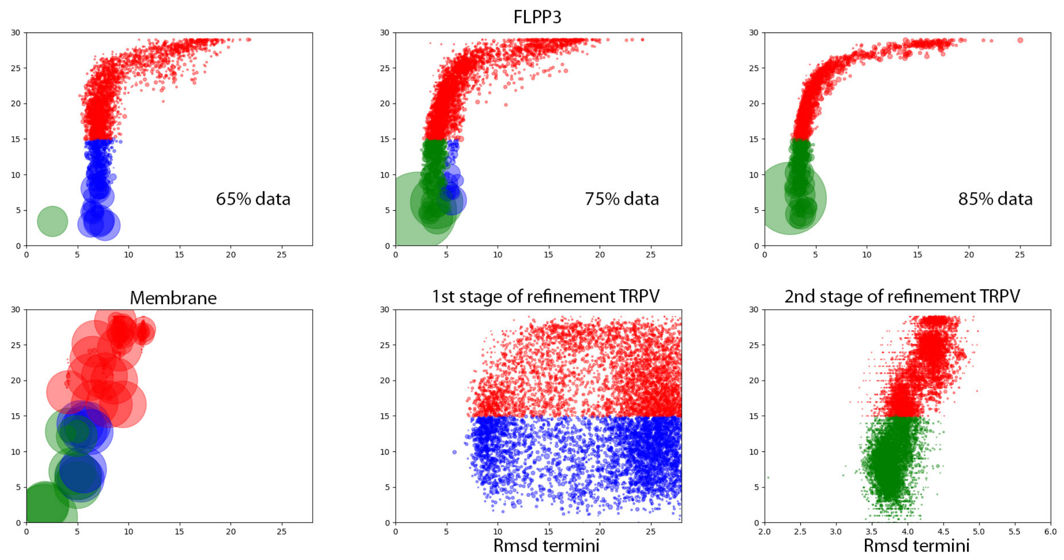

Figure S4: **Typical Funneling landscapes observed in MELD trajectories.** Clusters identified in high replica index (y-axis) are show in red, low replica indexes are colored blue for misfolded and green for correct topology. TOP panels correspond to FLPP3 runs enforcing contacts based on MDFF refinements. Trusting more data converges on large native-like clusters. Bottom left panel corresponds to the membrane protein refinement. Here the funneling effect is observed over a smaller range of RMSD values due to cartesian restraints being enforced. Bottom middle pannels corresponds to the initial refinement of termini conformation in the absence of specific MDFF contact information – a wide range of possible arrangements is observed with no preference, selected structures for MDFF are selected based on cross correlation with the density map. Bottom right panel shows MELD simulations after an MDFF step where MDFF contacts guide the structures towards more native like conformations.

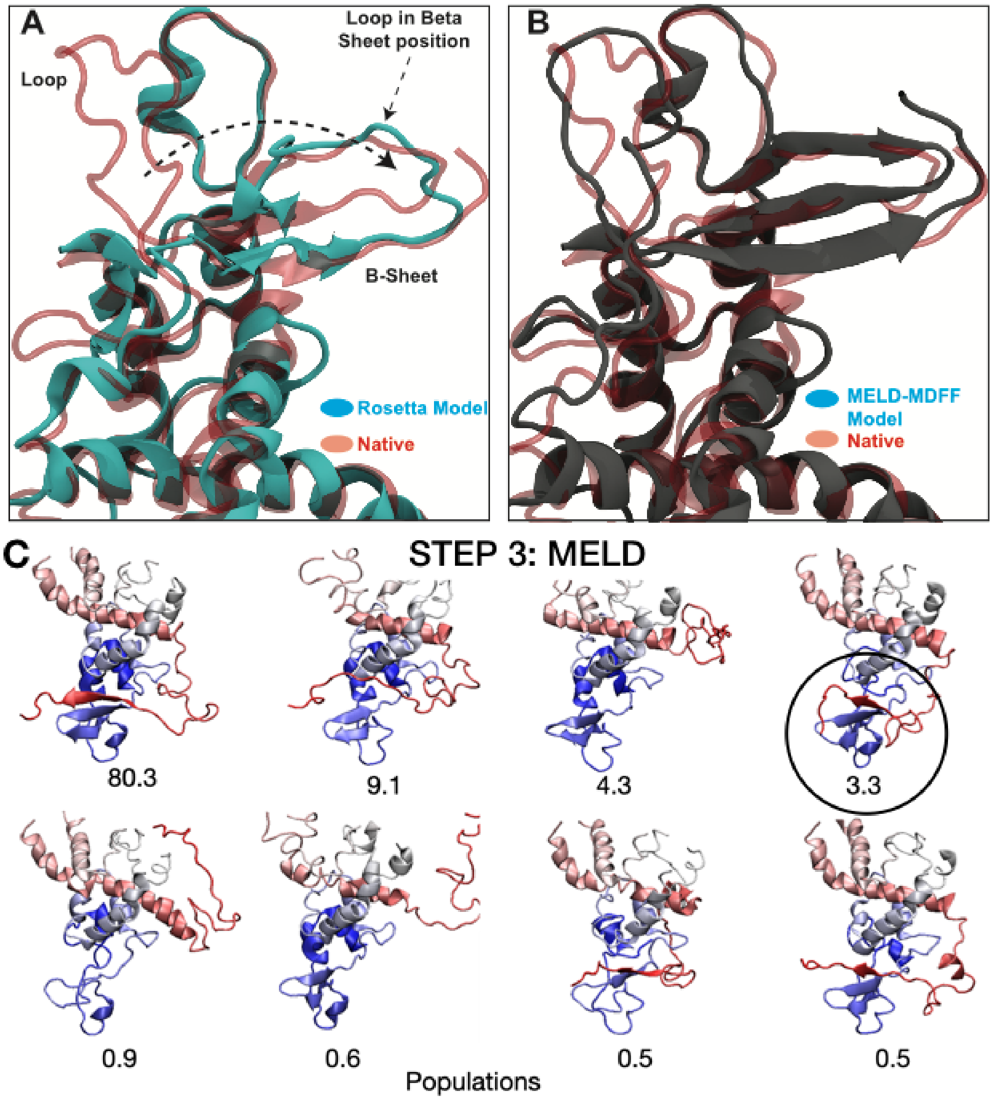

Figure S5: **Rosetta-ES model and CryoFold model for TRPV1.** (A) Comparison between the Rosetta-EM model (cyan cartoon) and the native structure (5IRZ, red transparent) of the cytoplasmic domain of TRPV1. Arrows denote erroneous modeling of the loop on the top of $\beta$ sheets in the Rosetta-EM model. (B) Comparing the CryoFold model (grey cartoon) and the native structure (red transparent cartoon). Note that the CryoFold model agrees well in both the loop and the $\beta$-sheet region with the native model. (C) Population of all the models derived in MELD.

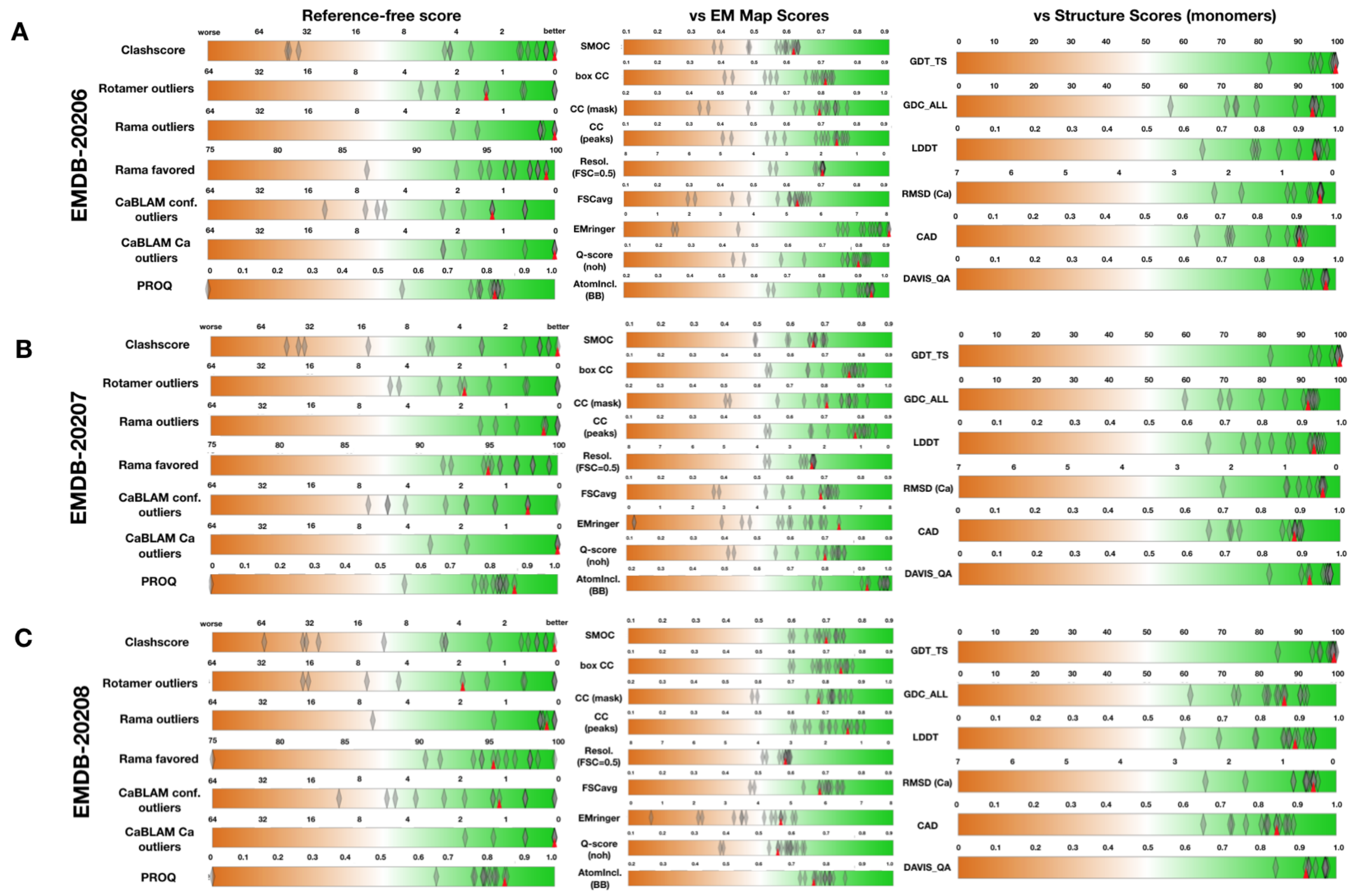

Figure S6: **Model metrics for apoferritin at three different resolutions in 2019 EMDB modeling challenge.** Assessors ranking of 17 teams based on quality of model and fit for three EM density maps (**C**) EMDB-20206 (1.8 Å), (**B**) EMDB-20207 (2.3 Å), (**C**) EMDB-20208 (3.1 Å).The ranking for the CryoFold team (Group 73) is shown by red inverted triangle. Overall the performance of CryoFold was among the top groups in this challenge.

A)

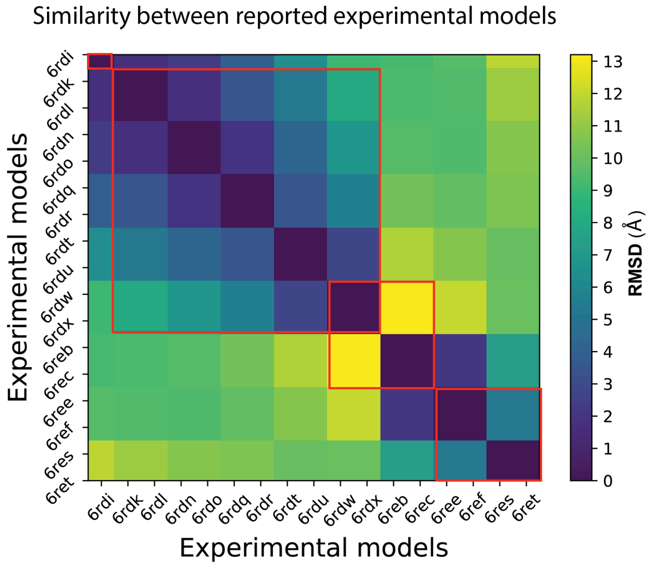

B)

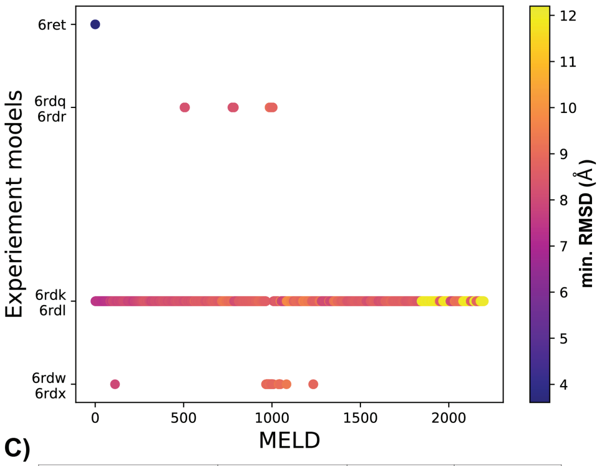

C)

| MELD-MDFF sampling of experimental models | PDB | Initial RMSD | Final RMSD |
|---|---|---|---|
| I | 6RET | 0.0 | 0.0 |
| II | 6RDQ, 6RDR | 7.6 | 2.1 |
| III | 6RDW, 6RDX | 12.0 | 2.8 |
| IV | 6RDK, 6RDL | 8.4 | 1.8 |

Figure S7: **Sampling rotary substates of mitochondrial $F_1$ - $F_0$ ATPsynthase in *Polytomella* sp.** (**A**) RMSD matrix showing the similarity between the different rotary substates deposited to PDB in 2019 by Murphy et al. (**B**) Minimum RMSD based clustering between structures obtained from MELD and experimental models. (**C**) Table for RMSD between initial model from MELD and final model after MDFF refinement with respect to target experimental models states I-IV.

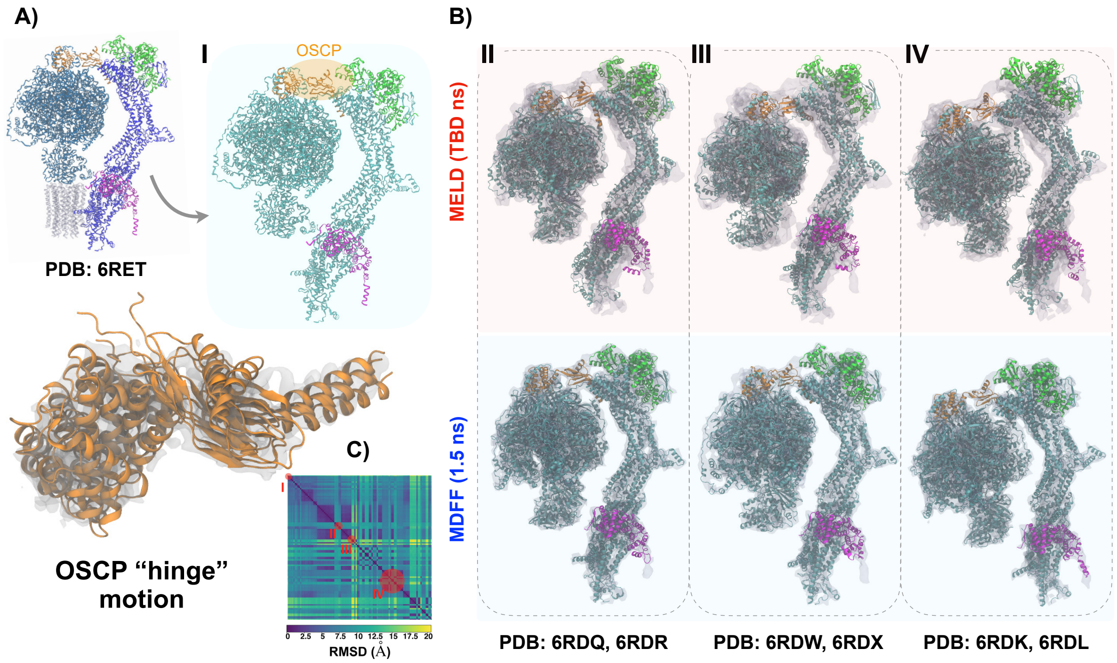

Figure S8: **Modeling of the soluble domain of mitochondrial $F_1$ - $F_0$ ATPsynthase.** **(A)** High resolution structure of a rotary substate of mitochondrial $F_1$ - $F_0$ ATPsynthase in *Polytomella* sp. deposited in 2019 (pdb 6RET), where the components used for MELD and MDFF refinement are illustrated by colors cyan, orange, green, blue and magenta. The *c* ring in the transmembrane region shown in gray was not used for the modeling and refinement. **(B)** Starting from one of the thirteen distinct conformation states (6RET), using RMSD matrices (bottom panel) the MELD trajectory were clustered in four distinct states (I: 6RET, II: 6RDQ, 6RDR, III: 6RDW, 6RDX, IV: 6RDK, 6RDL) marked by red circles. (top panel in light magenta shade) Structures from MELD (125 ns) corresponding to states II, III and IV. (bottom panel in light blue shade) MDFF refinement (1.5 ns) for structures initially obtained from MELD and fitted to their respective cryoEM-map density. The range of the resolution varied between 3.7 - 4.3 Å. **(C)** Conformational changes of the OSCP hinge which is responsible for mediating interdomain flexibility.

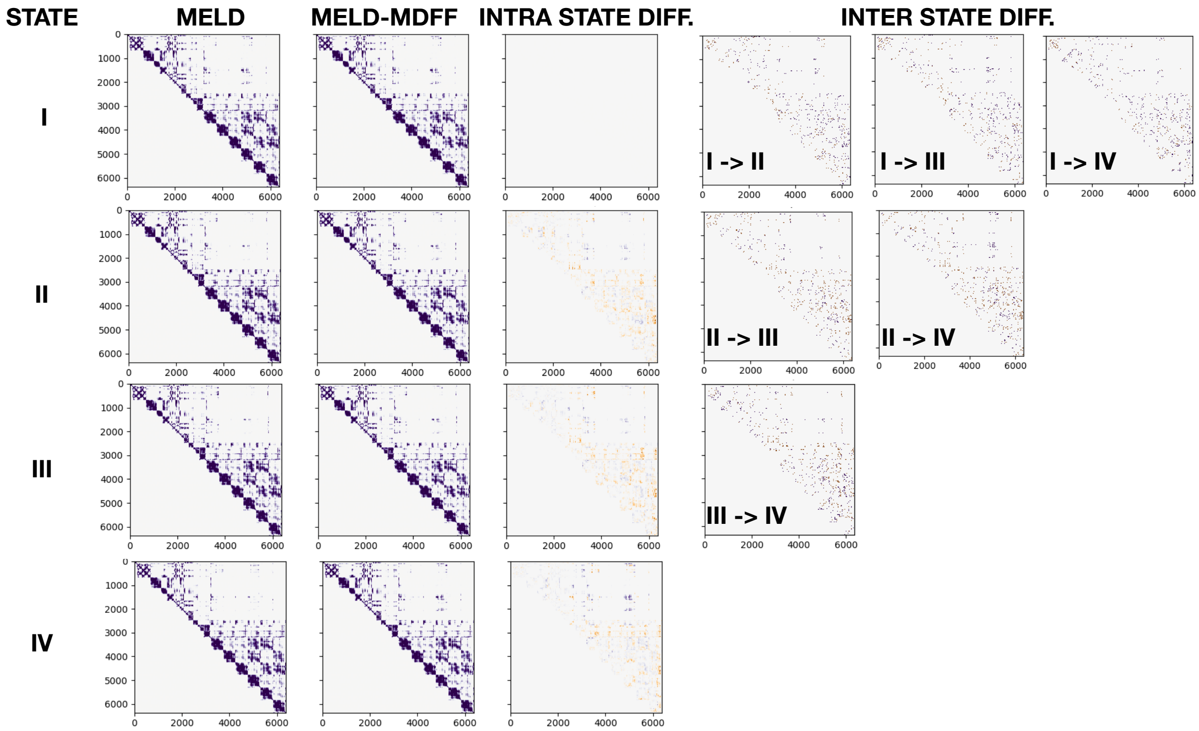

Figure S9: **Pairwise residue contact analysis for experimentally-verified substates I - IV of $F_oF_1$ ATP synthase from Polytomella sp.** The four rows in the figure panel indicate the change in the contact pairs in the initial and final models obtained from different states sampled by cryoFold. The first two columns illustrate the contacts in the final MELD and the final ReMDFF models. The intra-state difference (third column), represents the change in the number of tertiary contacts between the MELD and ReMDFF models within the same state. Highlighting changes in tertiary contacts across states, results in the fourth column illustrate that the interstate difference in contacts is much greater than that within an intrastate MELD-ReMDFF ensemble. These larger inter-state differences suggest that CryoFold captures the distinctive interface contacts of the $F_oF_1$ ATP synthase in each of the states I-IV, starting only with data that guides it towards state I. The broader space of protein conformations (beyond state I) that is captured by CryoFold, stems from MELD's capability of sampling states II, III and IV-like protein-protein contacts within the heterogeneous complex, which is beyond ReMDFF's sampling capabilities. Details of MELD's enhanced sampling of binding contacts is discussed elsewhere . The MDtraj package is used for calculating the distance and contact between residue pairs for the protein molecule. A cutoff of 4 Å is set to calculate the contact pairs.

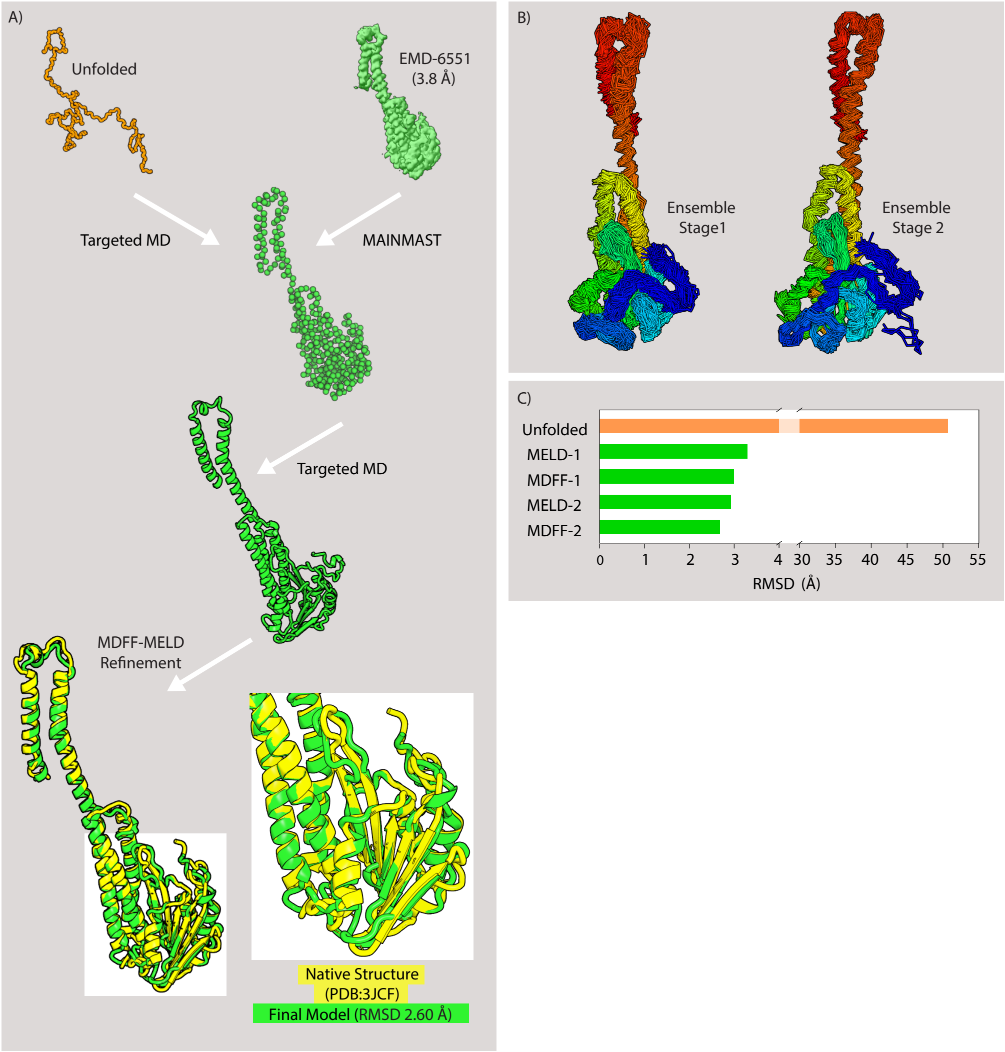

Figure S10: **Modeling transmembrane Magnesium-channel CorA.** The CryoFold protocol on CorA. A C$\alpha$ trace (green spheres) was generated from the Cryo-EM density map (EMD-6551) using MAINMAST. A random coil (orange) was fitted to this map using targeted MD to generate initial search model (green cartoon). Two concurrent rounds of MELD-ReMDFF refinement generated a structure that agrees extremely well with the native structure (yellow). The resultant CryoFold model is at an RMSD of 2.60 Å relative to the native structure and features accurate beta structures. (B) Ensemble of models at MELD refinement stage step1 and step 2. Ensemble models of step3 is showing convergence after three iterations. (C) The evolution of the RMSD of CryoFold models with each MELD-ReMDFF refinement.

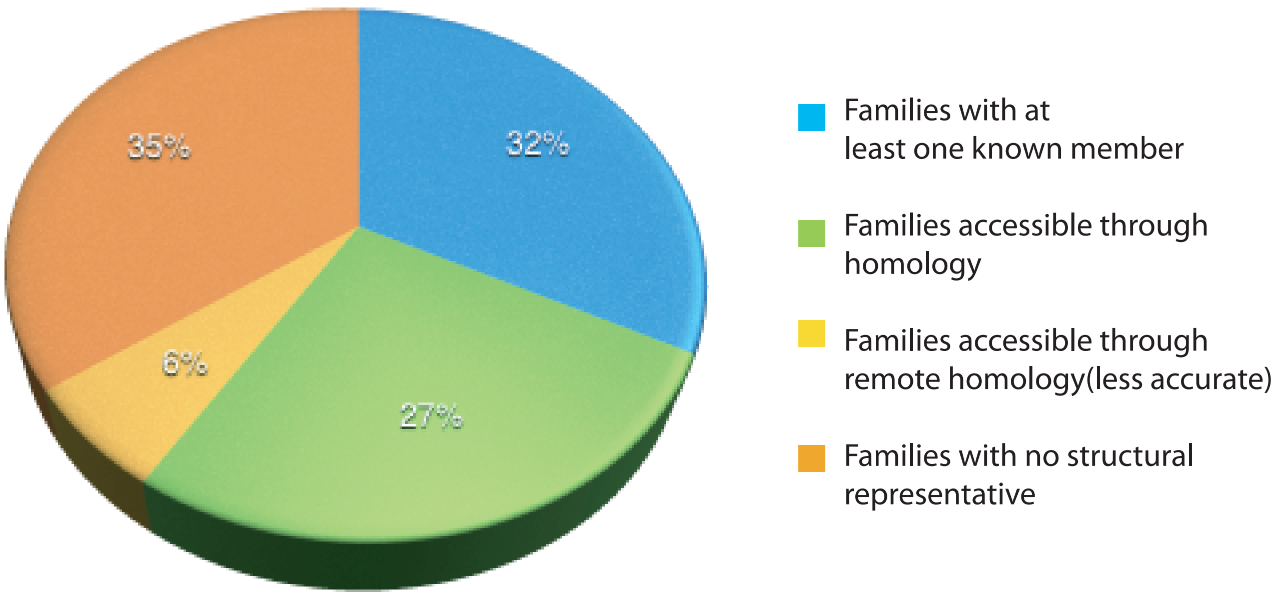

Figure S11: **Percent of protein families with structural representatives.**

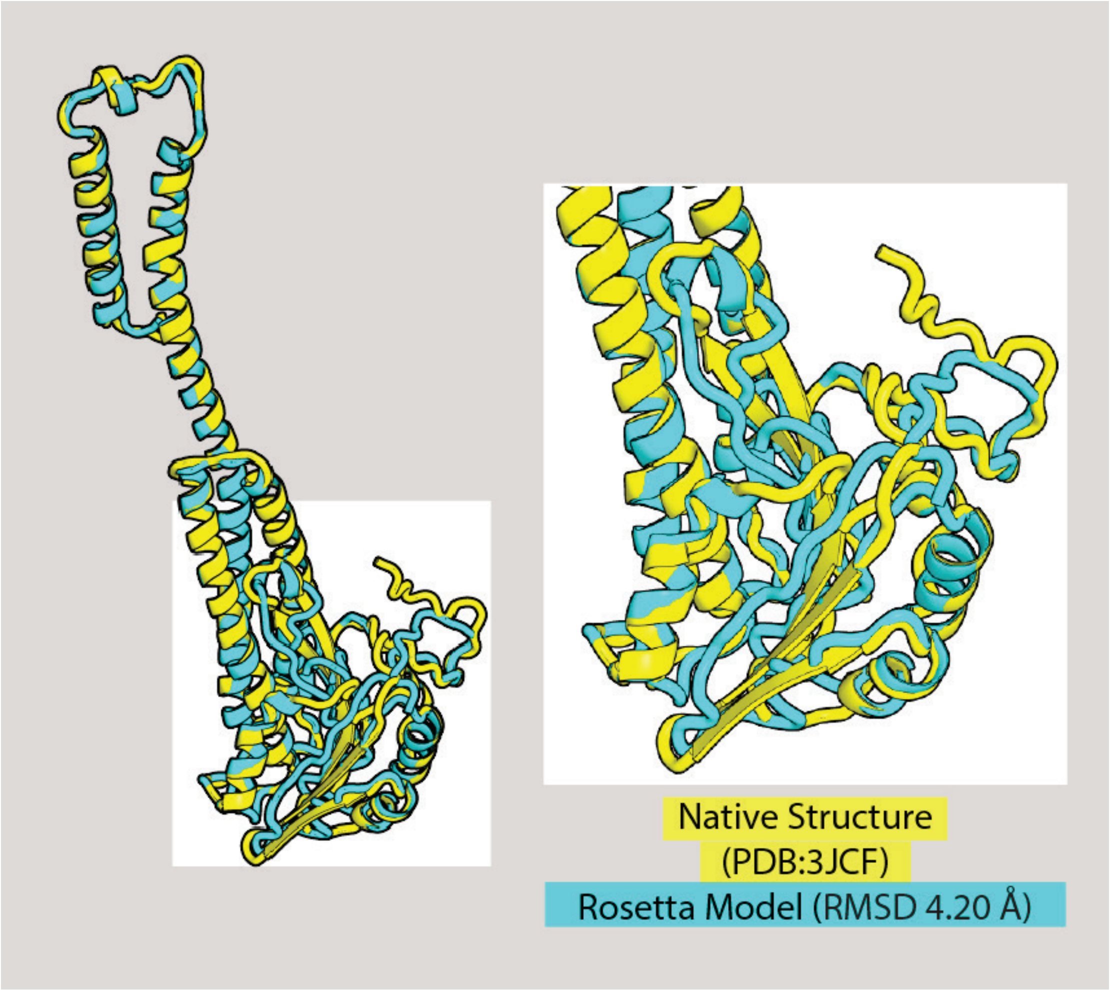

Figure S12: **Modeling transmembrane Magnesium-channel CorA.** The Rosetta-EM model (cyan) and the native structure (yellow). This model is at an RMSD of 4.20 Å with respect to the native structure.

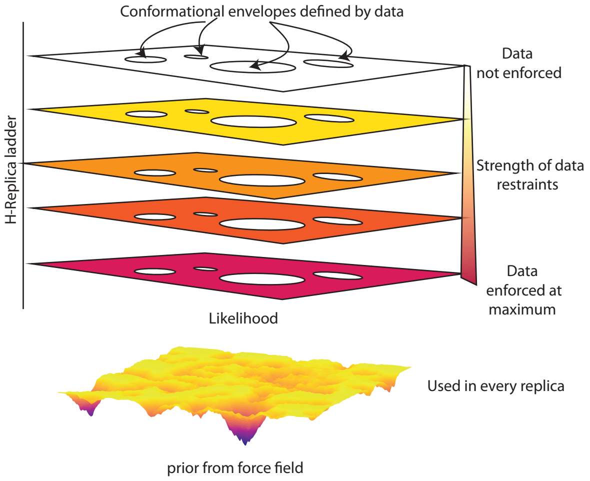

Figure S13: **MELD samples from a posterior distribution biased by agreement with data.** MELD tries to satisfy subsets of noisy data, effectively creating conformational envelopes where there is no bias and other regions that funnel structures towards the closest envelope. A hamiltonian and temperature replica exchange ladder is used where the data is not enforced at the highest replica (where the temperature is also high) and are strongly enforced at the lowest replica ladder.

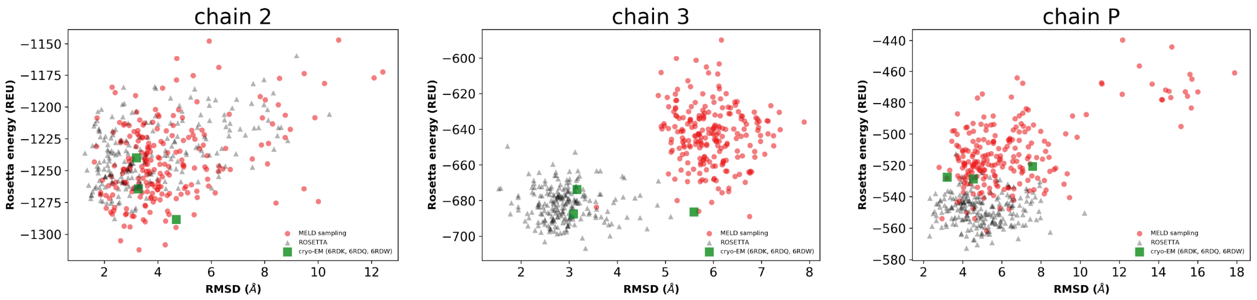

Figure S14: **Energy based model quality of MELD-MDFF trajectory for individual chains in mitochondrial $F_oF_1$ ATPsynthase in *Polytomella* sp.** Three chains (2, 3 and P) were identified based on maximum RMSD to the starting MELD model. Starting from the initial model (6RET), state I, Rosetta relaxation was performed on 220 models from the MELD-MDFF trajectory (red circles). Independently, 220 models were generated using Rosetta-Relax algorithm with the same initial model (gray circles). Finally, Rosetta-relaxation was performed on experimental models PDB: 6RDQ, 6RDW and 6RDK corresponding to states II, III and IV respectively, clustered from the RMSD matrix. Overall, the conformational free energy of MELD-MDFF models agrees very well to experimental models with a larger radius of convergence.

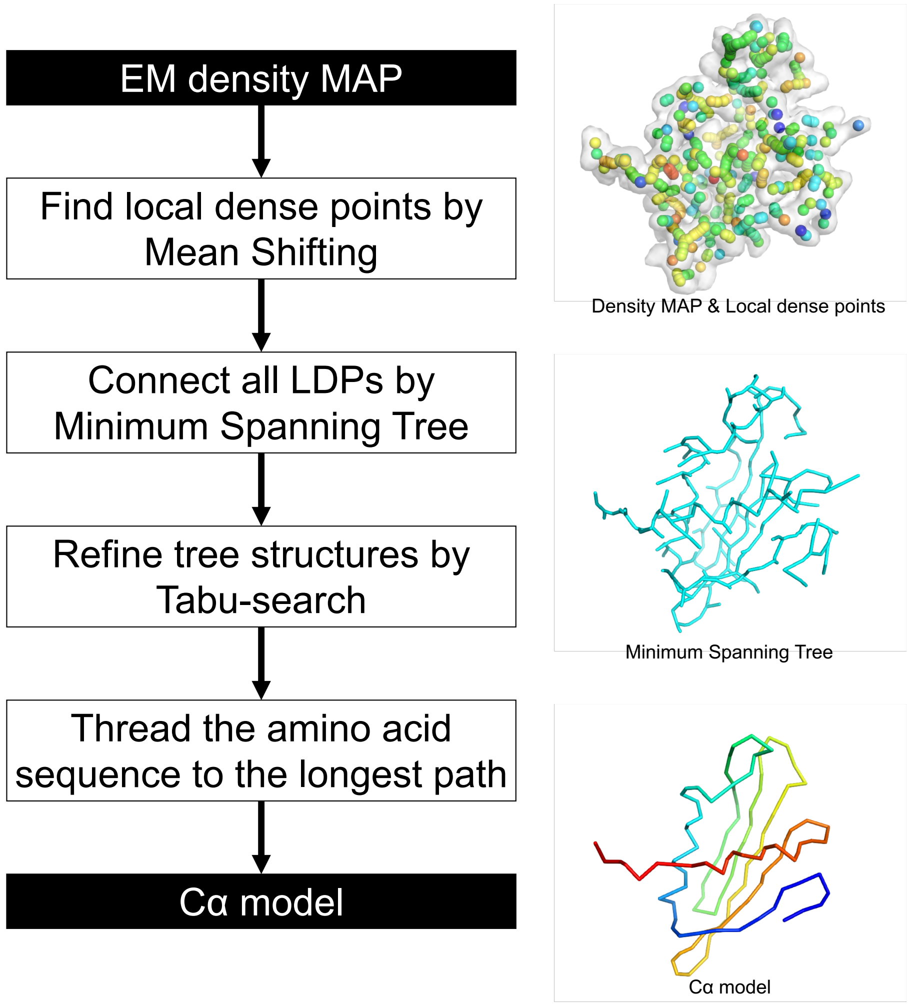

Figure S15: **MAINMAST protocol.** Four steps of MAINMAST algorithm are illustrated with example for flpp3. First, local dense points are identified with the mean shift algorithm. Identified local dense points are connected by minimum spanning tree (MST) (cyan). Using tabu-search, the MST is refined, iteratively. The amino acid sequence of the query protein is mapped on the longest path in the tree. C$\alpha$ models from each generated longest path are ranked with the density–volume matching (threading) score. In the first panel on the right, the local dense points are colored by the scale of density, blue to orange for low to high density. The second panel shows the minimum spanning tree. In the third panel, the chain represents a predicted C$\alpha$ model.

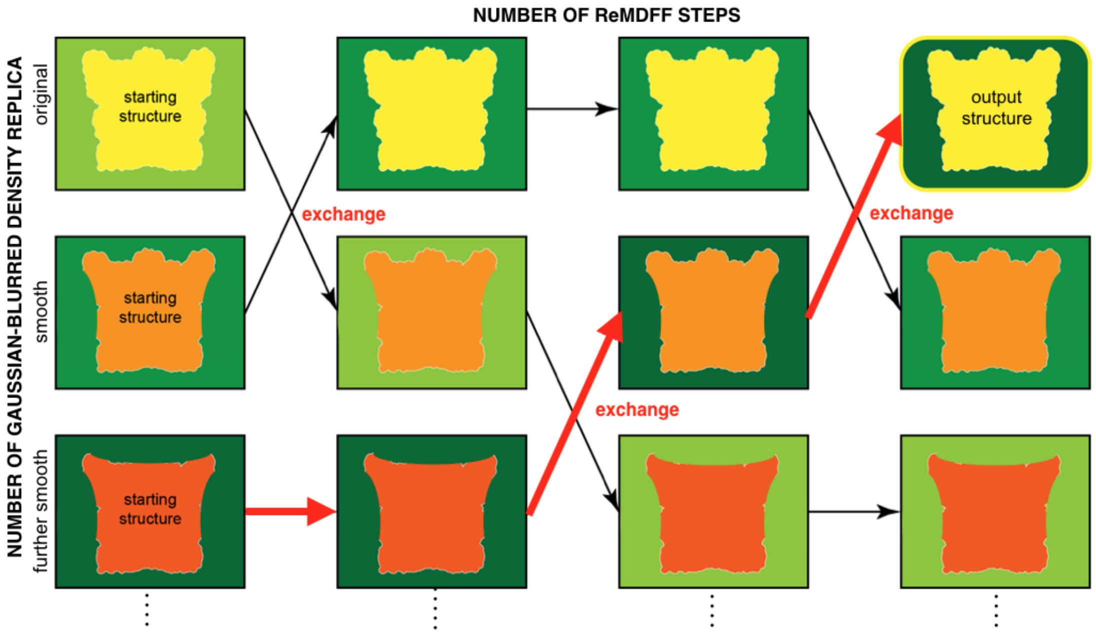

Figure S16: **A simple example of ReMDFF.** Three replicas are included in this schematic. Each replica consists of a molecular structure and a cryo-EM map-based grid potential. Different green boxes represents grid potentials of different resolutions. The structural models as refined at different resolutions are shown in red, orange, and yellow with different hue levels representing changes in the conformation. The arrows indicate the transfer of a grid potential from one replica to another. The output structure is selected from the trajectory visited by the grid potential of the original resolution (dark green).

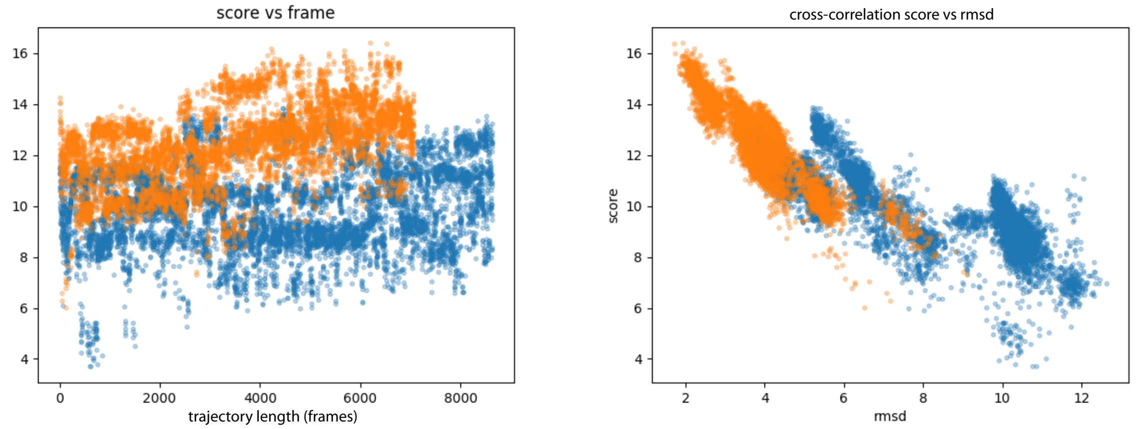

Figure S17: **Analysis of MELD-ReMDFF trajectories.** Cross correlation score vs (A) trajectory length or (B) rmsd to native. The cross correlation score increases through simulation time and is a good indicator of protein quality when compared to rmsd (Å). Colors indicate two starting structures: the blue dots correspond to a trajectory that comes from a low resolution backbone tracing and the orange ones from a high resolution backbone trace.

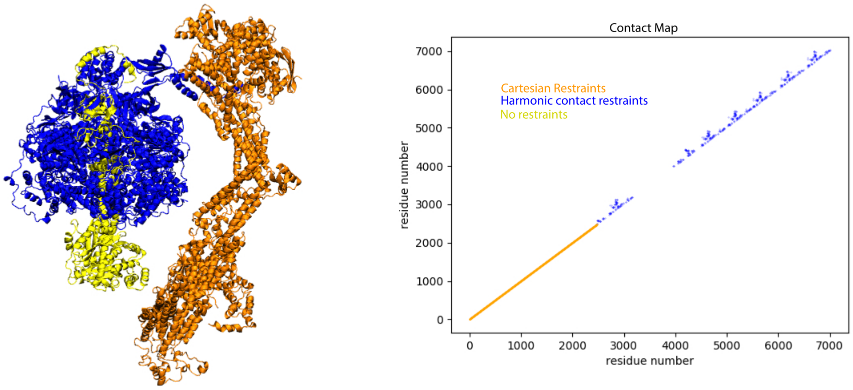

Figure S18: **Data used for the ATP syntahse simulation.** We restrained the orange region with harmonic cartesian restraints on C$\alpha$ atoms, we imposed flat bottom harmonic restraints in the domains corresponding to the blue region (see contact map on the right) to maintain domain architecture and left the atoms in the yellow region free.

11. Chimera. UCSF Computer Graphics Laboratory. San Francisco, CA. http://www.cgl.ucsf.edu/chimera.

21. Lange, O. F. & Baker, D. Resolution-adapted recombination of structural features significantly improves sampling in restraint-guided structure calculation.

*Proteins: Structure, Function, and Bioinformatics* **80**, 884–895 (2012). URL https://doi.org/10.1002/prot.23245.

22. Brandon Frenz, F. D., Ray Y.-R. Wang. Tutorial: Rosetta tools for structure determination in cryoem density (2018). URL https:/faculty.washington.edu/dimaio/files/rosetta_density_tutorial_aug18.pdf.

23. Frenz, B., Walls, A. C., Egelman, E. H., Veesler, D. & DiMaio, F. RosettaES: a sampling strategy enabling automated interpretation of difficult cryo-EM maps. *Nature methods* **14**, 797–800 (2017).
